# Genomic analyses reveal the origin of domestic ducks and identify different genetic underpinnings of wild ducks

**DOI:** 10.1101/2020.02.03.933069

**Authors:** Rui Liu, Weiqing Liu, Enguang Rong, Lizhi Lu, Huifang Li, Li Chen, Yong Zhao, Huabin Cao, Wenjie Liu, Chunhai Chen, Guangyi Fan, Weitao Song, Huifang Lu, Yingshuai Sun, Wenbin Chen, Xin Liu, Xun Xu, Ning Li

## Abstract

Domestic ducks are considered to have been tamed from the mallard or a descendant of the mallard and the spot-billed duck. Domestic ducks show remarkable phenotypic variation in morphology, physiology and behaviour. However, the molecular genetics of the origin and phenotypic variation of ducks are still poorly studied.

Here, we present mallard and spot-billed genomes and perform whole-genome sequencing on eight domestic duck breeds and eight wild duck species. Surprisingly, analyses of these data support a model in which domestic ducks diverged from their closest wild lineage (mallard ducks and spot-billed ducks) at the last glacial period (LGP, 100-300 kilo years ago (Kyr)). The wild lineage further speciated into mallard ducks and spot-billed ducks approximately 70 Kyr, whereas the domestic lineage population decreased through the LGP. A scan of wild duck genomes compared with domestic duck genomes identified numerous loci that may have been affected by positive selection in ancestral wild ducks after their divergence from domestic lineages. Function analyses suggested that genes usually affecting organ development and energy metabolism may involve long-distance flight ability. Further selective sweep analyses identified two genes associated with egg production and three genes related to feeding modulation under selection in domestic ducks. These analyses unravel a distinct evolutionary pattern of ducks and two wild duck *de novo* genomes, thus providing a novel resource for speciation studies.

## Introduction

Domestication and speciation (or population divergence) has drawn a lot attention in evolutionary studies especially since the emergence of new genomic methods with the deluge of empirical sequencing data (Andersson, 2001; Axelsson et al., 2013; Burri et al., 2015; Carneiro et al., 2014; Ellegren et al., 2012; Frantz et al., 2015; Rubin et al., 2012; Rubin et al., 2010; Seehausen et al., 2014). The genome assembly and characterization of genetic variation in many domestic and wild animals have revealed their complex demographic history, thus providing profound knowledges for prerequisite of both domestication and speciation studies (Carneiro et al., 2014; Frantz et al., 2015; Nadachowska-Brzyska et al., 2013; Rubin et al., 2012). We previously drafted a Beijing duck genome and identified genomic variation within this assembly (Yinhua Huang et al., 2013). Despite the genetic variations of duck have been further characterized to explore the genetic variations associated with specific traits (Zhang et al., 2018; Zhou et al., 2018). The domestic origin of ducks and the general genetic basis of the difference between domestic duck and their closest wild relative are still worth to pursue.

There are currently more than 140 wild duck species worldwide, and most species can hybridize with each other (Lavretsky, McCracken, & Peters, 2014). The phylogeographic of a recent radiation in the Aves:Anas were inferred from genomic transect and mallard duck diverged from its closest relative (spot-billed duck) recently was further determined using the mtDNA control region and ODC-6 of nuclear DNA (Zhuravlev & Kulikova, 2014). Traditional views suggest that domestic ducks were originated from mallards or the descendant of mallard duck (*Anas platyrhynchos*, for the sake of simplicity, we referred as mallard in this paper) and spot-billed duck (*Anas zonorhyncha*, refer as spot-billed)(Steve, 1992). Those are mainly based on their similar outlooks, habits, and distributions. Moreover, there are broad phenotypic differences between domestic ducks and their wild relatives, such as loss of flight and migration ability, reduced brain size, and increased body size and fertility. Although researches of steamer ducks on flightless and genetic association with selected traits on domestic ducks have been performed (Zhang et al., 2018; Zhou et al., 2018). They revealed diverse mechanisms from others and It indicating a historical context would be important in understanding functional changes and its ecology factors (Burga et al., 2017; Campagna, McCracken, & Lovette, 2019). However, the demographic inference suffered from lacking of solid history information, too many possibilities, complex history and no robust mutation rates. The historical relationships of ducks, like many other domestic animals, are still unclear (Albarella, 2005; Larson & Fuller, 2014; Madge & Burn, 1988; Richards, 2017; Stern, 2000).

Despite, lots of efforts have committed and traditional domestication hypotheses have been found incompatible with genetic data, and the direct ancestors of domesticated animals are still unclear (Frantz et al., 2015; Freedman et al., 2014). Nevertheless, some genetic analyses were still conducted under the pre-assumption of some untested recoding and questionable hypothesis (Irving-Pease, Frantz, Sykes, Callou, & Larson, 2018).

In this research, we surprised found excellent case of ducks to understand the domestication process. Because the central issue on the time of domestic lineage origin could be derivation through the splitting orders of domestic ducks, mallards and spot-billed. As the provocative of our results that the domestic lineage may diverged from mallard and spot-billed duck prior to the two wild ducks’ splitting, which put forward a time depth longer than human civilization’s origins. Multiple methods were used to bolster our finding that are from solid and basic principles, different aspect of data. We will demonstrate that how those results conducted from different approaches were logically matched with each other. In the end, we characterized the genomic divergence of mallard and spot-billed and the introgression effects to illustrate very recent divergence time between the two wild ducks.

## MATERIALS AND METHODS

### Sample collection

A total of 15 mallard, 14 spot-billed and 8 other species wild ducks (with hunting permission granted by the local government,) together with 114 domestic ducks were analysed in this study. Details in the geographical distribution of samples collected and sequenced can be found in Fig. S1; Table S1 and can be summarized as follows: 9 mallard ducks (9/15) and 8 spot-billed (8/14) from Fenghua, Zhejiang province (121° 39′E 29°67′N); 6 mallard ducks (6/15) and 6 spot-billed (6/14) from Hangzhou, Zhejiang province (120°21’E′ 30°25′N). Among them, 1 female mallard and 1 female spot-billed from Fenghua were used to perform de novo sequencing with Illumina technology (BGI-Shenzhen, China). 8 other species wild ducks were collected from Poyang late, Jiangxi province (116°59′E 28°96′N) and Fenghua, Zhejiang province, separately.

**Table 1.**
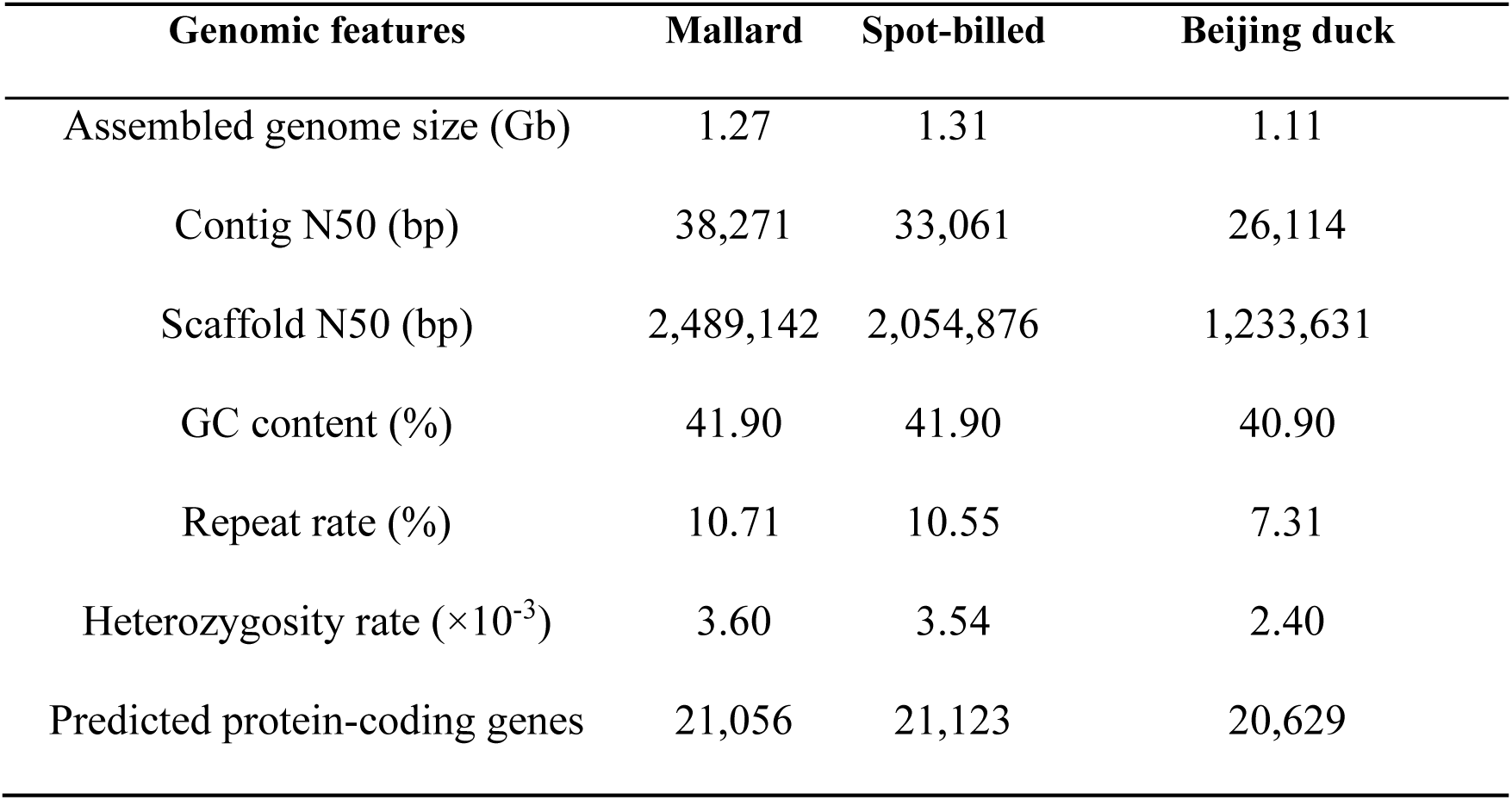
Comparison of features of the mallard, spot-billed and Beijing duck Genomes.

### Assembly and annotation

The assemble of the two wild ducks were performed using the similar pipeline as Beijing duck assembly to exploit the specific genomic feature of wild or domestic duck and access the influence of genome assemble on subsequent variation analysis. Sequences from paired-end reads of 17 libraries for each individual with insert sizes ranging from 250 bp to 20,000 bp were assembled with SOAPdenovo and SSPACE software (Table S1). The super-scaffolds were positioned along the chromosomes based on the duck cytogenetic map and the comparative genomic map between duck and chicken (Yinhua Huang et al., 2013; Y. Huang et al., 2006). Reference gene sets for the mallard genome and the spot-billed genome were retrieved by merging the homologous set, the transcript set (predicted with 21 duck transcriptomes) and the *de novo* set (Yinhua Huang et al., 2013).

### Whole-genome resequencing

114 domestic ducks were chosen from 8 breeds which only Beijing ducks and Shaoxing ducks were sequenced by individual. Other 6 breed samples and 2 outgroups were pools of DNA samples from 10 individuals each in equimolar quantities. 35 wild ducks from eight species (gadwall (n=1), Baikal teal (n=1), pintail (n=1), falcated teal (n=1), common teal (n=1), wigeons (n=3), spot-billed (n=13) and mallard (n=14)) were individual sequenced. DNA for whole genome resequencing was prepared according to the instructions of Illumina technology (BGI-Shenzhen) extracted from blood in each case. Paired-end reads were generated from wild ducks, domestic ducks, Muscovy ducks and Taihu geese then aligned to Beijing duck assembly (BGI_duck_1.0) using the bowtie2 with default parameters. Duplicated reads were removed individually from samples using Picard and realignment process are performed before SNP calling. After removing reads that map to multiple place, we called SNPs using GATK and filtered SNPs according to the coverage with thresholds of ≤ one third of the mean or ≥ three-fold of the mean and MQ value ≤ 28 (DePristo et al., 2011). SNP used in IBS, PCA and genetic structures were called from multiple bam samples by using joint variant calling mode of GATK. We merged individual bam files into one population bam sample and jointly call SNPs with pool samples, or random sample bam reads into same coverage when necessary. SNP validation and other detail process such as SNP pruning see SI Appendix section 4.

### ^∂^a^∂^i simulation and PSMC analysis

For the ^∂^a^∂^i simulation, SNPs frequency of mallard, spot-billed and other 8 domestic populations were obtained from each of the 10 VCF files, then combined by using a way like outer join of SQL. (i.e. include SNPs that were monomorphic within one population but variable in their combined samples (10 populations); SI Appendix). After LD pruning, We then inferred ancestral alleles by using Muscovy ducks and Taihu geese as outgroups using the following thresholds: (1) more than 30-fold coverage when SNP fixed in Muscovy or both outgroups; and (2) the both outgroups were fixed the same directions. Subsequently, we quantized the frequency of each SNP of 8 domestic duck populations into one population and simulated the divergence among spot-billed, mallard and “domestic” ducks using split without migration, split with migration and IM models with 50 bootstraps each (SI Appendix 4.6). *Nref* was calculated with the following format: 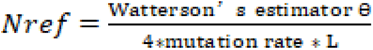; and L represents the length of the SNPs from and diluted by the rate of SNP sampling.

In both PSMC (Li & Durbin, 2011; Lohmueller, Bustamante, & Clark, 2010) and ^∂^a^∂^i simulations, We assumed that the mutation rate per year was 9.97e-10 and the generation time was one year (Nadachowska-Brzyska, Li, Smeds, Zhang, & Ellegren, 2015).

### Positive selection in early ancestors of mallard and spot-billed ducks

To detect selection that occurred in early wild duck lineage (mallard and spot-billed) after their divergence with the domestic lineage, we used a method derived from (Green et al., 2010) that is particularly suited to detect older selective sweeps during or shortly after divergence. We selected SNPs that derived allele existed (including fixed) in mallard or spot-billed. Ancient sweeps were enriched mainly through searching for genomic regions with dearth of derived allele shared with domestics among wilds. (i.e. selected and linked SNPs reached high frequency to fixed. Whereas the allele frequency spectrum of those regions will be recovered by wild duck alleles, that is not shared with domestic lineages, with frequency spectrum towards normal to higher frequency since age old). This method will naturally disregard regions of recent selection or purifying selection in wild because those regions with derived allele skewed towards lower frequency wild alleles and making lower predicted domestic frequency thus leading weak signal (SI Appendix).

We calculated the derived allele frequency of 18,083,967 selected SNPs in both the wild and domestic lineage. As expected, they were highly correlated (Fig. S20) This correlation was used to calculate the expected derived allele frequency in the domestic lineage as a function of the frequency in the wild lineage. log ratio of the sum of observed/expected derived frequency in 20-kb windows with a step size of 10 kb were applied to scanning. After Z-transforming the genome-wide distributions, we arrived at measure S and defined selective sweep regions of the early mallard and spot-billed ancestors using the thresholds of -0.2819 (4-fold standard deviations away from the mean) for autosomes and -0.3089 for sex chromosomes (1.5-fold standard deviations away from the mean). Recent selective sweep regions in the domestic lineage could also reduce the S value; thus, we excluded regions that show lower zHp (CDRs) in the domestic lineage.

## RESULTS

We report high-quality drafts of the mallard and spot-billed and compare them to the genome of Beijing ducks and other birds (Table 1; SI Appendix; Fig. S1; Fig. S3-S11; Table S2-S10). We also generated a total of 1,219 Gb whole-genome resequencing data of 35 wild ducks from eight species, 114 domesticated ducks from six Chinese breeds and two commercial breeds (Cherry Valley and Campbell ducks), and 10 birds from Muscovy ducks and Taihu geese each as outgroups (Fig. S2; Table S1).

Variants were called using GATK for each population and different combinations separately, and several quality filters were applied. In total, 44,852,612 SNPs were identified in the 10 duck populations (eight domestic duck breeds and two wild duck populations), and 20,676,377 and 42,012,846 SNPs were detected in domestic duck and two wild duck populations (the mallard and spot-billed), respectively, with 18,093,115 shared SNPs (Fig. S12). Additionally, mallard and spot-billed ducks shared a comparative number of SNPs with domestic ducks (SI Appendix).

### Domestic duck originated in a lineage distinct from the mallard and spot-billed

We explored the genetic relationships of domestic and wild ducks by performing a principal component analysis (PCA) initially with genome-wide SNPs from 27 Shaoxing and 27 Beijing ducks (individually sequenced domestic ducks) and 35 wild ducks. Expectedly, the domestic ducks (Shaoxing and Beijing ducks) clustered to mallard and spot-billed while clearly separating from other wild species (Fig. 1a; SI Appendix 4.5). Such genetic relationships were also identified by further PCA, and tree analyses based on the IBS (identical by state) distance of those 89 individuals (Fig. 1b; Fig. S13-14). These findings suggest that the mallard and spot-billed have a closer genetic relationship with domestic populations than other wild species.

**Figure 1:**
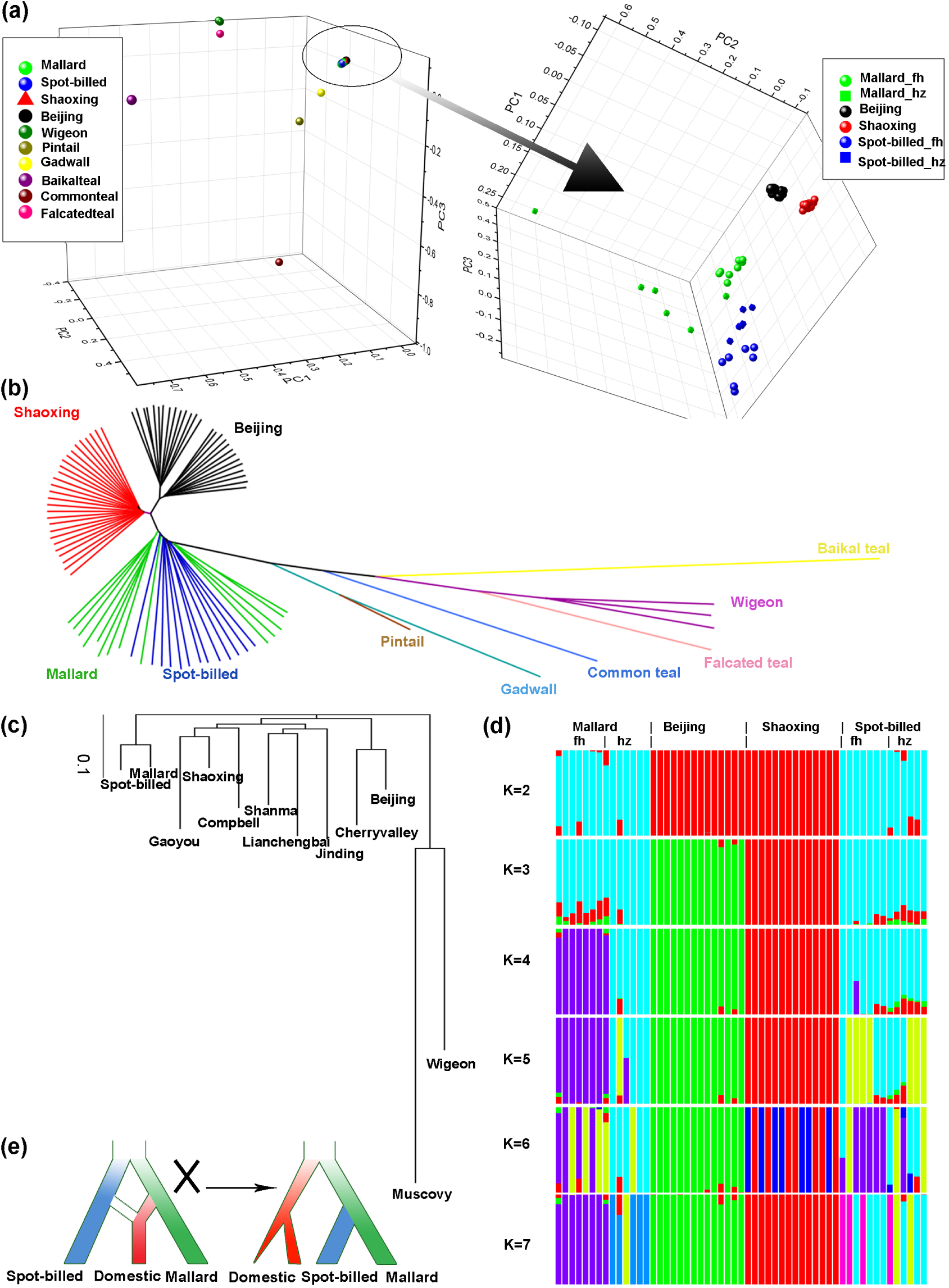
Inference of phylogenetic topology in wild and domestic ducks. Beijing and Shaoxing ducks representing domestic ducks were used in the principal component analysis (PCA), phylogenetic analysis based on identity-by-state (IBS) and genetic structure analyses. “_fh” and “_hz” mean spot-billed and mallard collected from Fenghua and Hangzhou, respectively. (**a**) PCA plot constructed for Principal Component 1 (PC1), PC2 and PC3 using 10,525 high-quality filtered SNPs from 27 Beijing and 27 Shaoxing ducks and 35 wild ducks (one Baikal teal, one gadwall, one pintail, one falcated teal, one common teal, three wigeon, 13 spot-billeds and 14 mallards) shows that domestic ducks are close to spot-billed and mallard wild ducks. (**b**) Neighbour-joining tree based on IBS distance of 80,505,576 SNPs. (**c**) Neighbour-joining tree based on genome wide pairwise fixation index (F_ST_) of 44,852,612 SNPs. (**d**) Genetic structure analyses of 13 spot-billed and 14 mallard wild ducks along with 14 Beijing and 14 Shaoxing domestic ducks using an admixture ranging *K* from two to seven. (**e**) Inferred demographic models for wild and domestic ducks inferred based on the genetic analyses of **a**-**d** (right). The traditional demographic model, which hypothesizes that domestic ducks originated from mallards or the descendant of a mixture of spot-billed and mallard ducks (left), is incompatible with observations from the neighbour-joining trees (**b**-**c**) and genetic data (**d**).

We then asked whether domestic ducks originated from the mallard or the descendants of the mallard and spot-billed ducks. We performed a 3-population test for admixture by taking the mallard and spot-billed as ancestral populations and Beijing and Shaoxing as target populations through a Block Jackknife analysis that corrected for linkage disequilibrium among SNPs (Reich, Thangaraj, Patterson, Price, & Singh, 2009; Rodgers, 1999). Surprisingly, this effort obtained significantly positive scores for these four groups (f3=0.007279, Z score=23.834; Table S11) that did not demonstrate that domestic ducks were descendants of the mallard and spot-billed ducks. However, the mallard specific SNPs (polymorphism present only in mallard and ancestral allele fixed in spot-billed) and spot-billed specific SNPs were almost equally distributed into each of the 8 domestic breeds, which implied equal closely kinship (Fig. S12b).

We further performed genetic structure analysis to estimate ancestry among individuals of the four populations (Beijing, Shaoxing, mallard and spot-billed) with *K* (the number of ancestry components) ranging from two to seven (Alexander, Novembre, & Lange, 2009). This analysis indicated that domestic ducks had a homogeneous genetic background, whereas wild ducks showed a heterogeneous genetic background; this outcome is consistent with the PCA (Fig. 1a, d). However, the wild ancestor duck started to divide into mallard and spot-billed when *K* was increased to four, and it followed the separation between the wild ducks and domestic ducks when *K*=2, which had the lowest cross-validation error (Fig. 1d; Fig. S15; SI Appendix 4.5). These analyses indicated a deep divergence time between domestic lineage and wild than it between mallard and spot-billed, instead of traditional hypothesis (Fig. 1e). F_ST_-based phylogenetic tree also compatible with our hypothesis, and this evolutionary model could also explain the positive f3 and the comparative numbers of mallard/spot-billed specific SNPs observed in each domestic duck breeds. As those SNPs that shared by the wild ancestors and domestic ancestors draft into mallard and spot-billed specific SNPs equiprobably (Fig. 1c; Fig. S12b). Although the resolution of IBS tree and genetic structure seems couldn’t distinguish mallard and spot-billed well, may due to their very recent divergence stage, overall analyses indicated that they were monophyletic (SI Appendix 4.5.2; Fig. S13d,S15b).

To further confirm the evolutionary relationships of mallard, spot-billed and domestic ducks, we then randomly selected and polarized 20,333 unlinked SNPs from these 10 duck populations with two outgroups (Muscovy ducks) for a demographic simulation using the diffusion approach (^∂^a^∂^i) (Methods) (Gutenkunst, Hernandez, Williamson, & Bustamante, 2009). We estimated the divergent time of the domestic ducks (quantized all eight domestic breeds into one population) and two wild duck populations (mallard and spot-billed) using two simple but informative models, with one taking migration into account while the other did not (Fig. 2a; Methods; SI Appendix 4.6). One hundred bootstrap simulations of the two models show that both divergence times between the two wild duck populations and domestic ducks were almost equal (100-300 Kyr) and longer than the corresponding one between mallard and spot-billed ducks (approximately 70 Kyr, Fig. 2b-c; SI Appendix section 4.6; Table S12, Fig. S16-18). Which further support that the ancestor of the domestic duck split from the ancestor of mallard and spot-billed ducks before the two wild ducks’ divergence (Fig. 1e). As ∂ a ∂ i couldn’t distinguish short-divergence from high-migration, a noteworthy consequence of introducing migration factors is the divergence time of model 2 increased. Furthermore, it seems due to ∂ a ∂ i have difficulty to determine longer Ts accompanied by high migration or shorter Ts coupled with low migration rate, thus having simulation on both situations and produce two peaks of estimate Ts.

**Figure 2:**
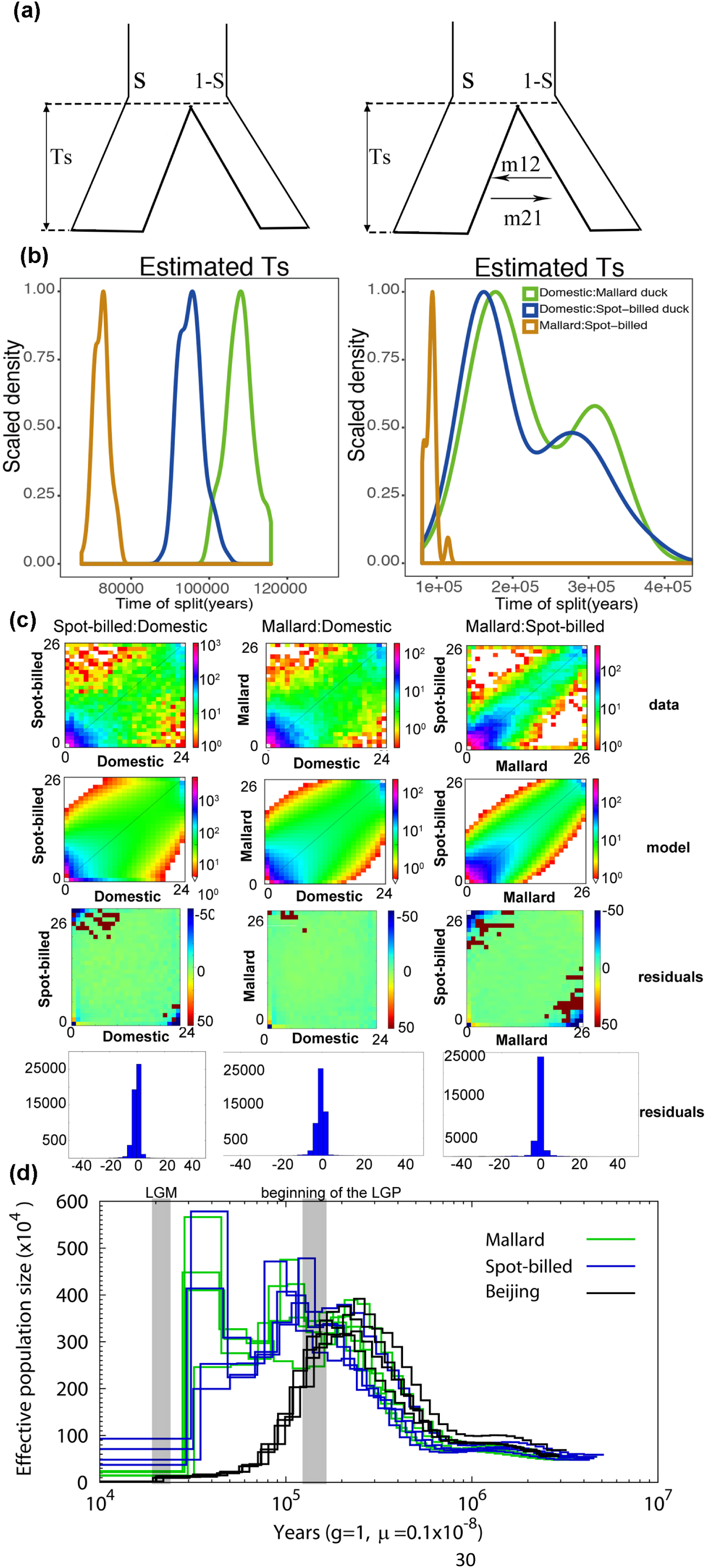
Inference of joint demographic history of ducks. Wild ducks included spot-billed and mallard ducks. Domestic ducks included Beijing, Cherry Valley, Campbell, Gaoyou, Jingding, Lianchengbai, Shaoxing and Shanma. (**a**) Illustration of the two models (with/without migration) used in the history inference through the ^∂∂^a^∂∂^i analysis. Parameters corresponding to the code are provided in the supporting information. (**b**) Distribution of estimated Ts (divergence time) of each model in 100 simulations between spot-billed and mallard (orange), mallard and domestic lineage (green), spot-billed and domestic lineage (blue). (**c**) Joint Site Frequency Spectrum (SFS) for observed data, simulation data and residuals. (**d**) Demographic history inference of mallard duck, spot-billed duck and Beijing duck through the PSMC analysis.

In addition, we performed diploid pairwise sequential Markovian coalescent (PSMC) analysis to reconstruct the demographic history and estimated the effective population size (*N_e_*) change along time among the mallard, spot-billed, Beijing and Shaoxing duck individuals. This analysis suggested that ancestors of these four populations presented a comparative effective population size before 110 Kyr (beginning of the LGP). However, ancestors of the two wild ducks and domestic ducks seemed to be under different demographic histories after the LGP. During this period, the effective populations of Beijing and Shaoxing duck ancestors decreased dramatically and continually. In contrast, the mallard and spot-billed duck ancestors maintain a relative high population size until LGM (Fig. 2d; SI Appendix; Fig. S19). This observation is consistent with previous research (Zhang et al., 2018) and the population structure and ancient introgression may make the curve of wild heterogenous. PSMC can also be informally used to infer divergent times when *N_e_* trajectories that are overlapping diverge and move forward in time towards the present (Freedman et al., 2014; Wang et al., 2016). Interestingly, the divergent time was roughly coincident with the one inferred from ^∂^a^∂^i. Such consistency further encouraged us to infer that the ancestor of domestic duck diverged from the ancestor of mallard and spot-billed at approximately the beginning of the LGP. Similar to the *N_e_* value estimated by the PSMC analysis, we found that the population size of domestic duck decreased whereas that of both wild duck species increased after the split between the wild and domestic population when using ^∂^a^∂^i to simulate a more complex model (i.e., population size changed exponentially after split) of divergence between populations (SI Appendix 4.6).

### Early ancestors of mallard and spot-billed might benefit from adaptive selections

Interestingly, the different population size change trends occurred after divergence between the wild lineage and domestic lineage during the LGP. One possibility might be that natural selection granted the mallard and spot-billed ancestors but not domestic duck ancestors new trait advantages to adjust to environmental change after their split. Consequently, environmental changes led to a decreased domestic ancestor population size but increased wild ancestor population size (Fig. 2d; SI Appendix). We detected positive selection in the early ancestors of these two wild populations after their divergence with the domestic lineage and identified a total of 434 putative selective sweep regions, with a total size of 10.86 Mb, using thresholds of 4 standard deviations from the mean (Fig. 3a-b; Fig. S20, S21; Methods) (Green et al., 2010; Groenen et al., 2012). Only 150k of these putative sweep regions were overlapped with regions subsequently detected by F_ST_ between wild and domestic, further indicated this signal capture different property. Among them, 168 selective sweeps harboured 202 protein coding genes while the remaining sweeps did not contain annotated protein coding genes (SI Appendix; Table S13). Recent studies have indicated that positive selection in intergenic and intragenic regions was associated with phenotypic variation (Carneiro et al., 2014; Groenen et al., 2012; Rubin et al., 2010). We then extended 180 kb in both the upstream and downstream regions that did not contain annotated genes and identified 162 closest genes in 156 of these extended regions (Table S13).

**Figure 3:**
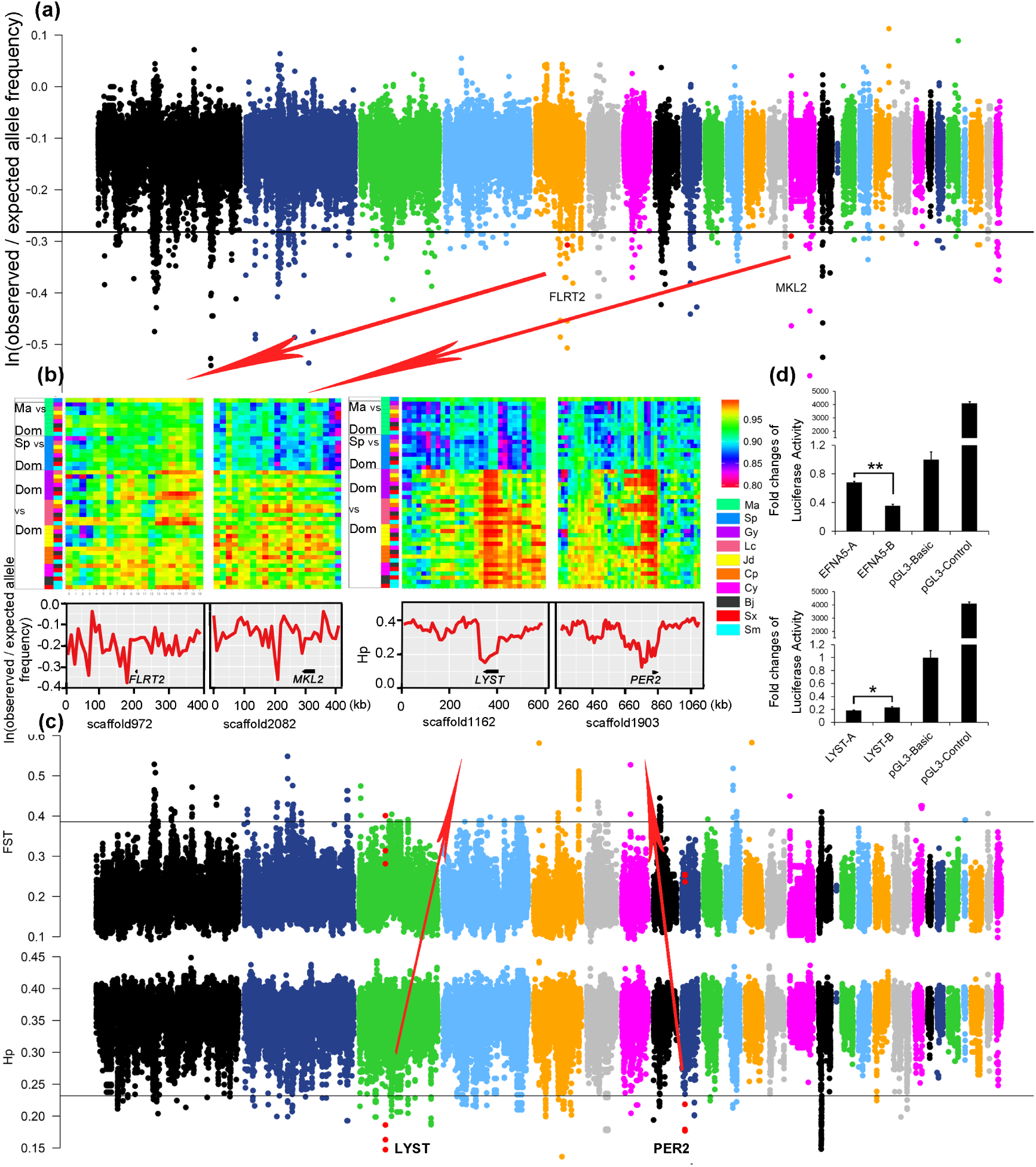
Summary of selective sweep analyses of the early wild duck lineage and domestic duck. (**a**) Distribution of ln(observed/expected derived allele frequency) calculated in a 20 kb window with a sliding size of 10 kb along scaffolds that are arranged according to their alignment to chicken autosomes. The horizontal black line indicates the threshold at ln(observed/expected derived allele frequency)=-0.3136 (mean of ln values - 4 standard deviations). Outlier regions that were visualized in detail by identity scores (IS) and re-computing values in smaller window sizes in (**b**) are highlighted with red dots. (**b**) Details of four putative selective sweep regions around the *FLRT2, MKL2, LYST* and *PER2* genes. The top of each panel shows the degree of haplotype sharing in pairwise comparisons among populations. Coloured boxes (left) indicate the comparison performed on that row. “Ma,” “Sp,” and “Dom,” represent mallard, spot-billed and domestic duck, respectively. The y-axis (downside) of *FLRT2* and *MKL2,* which are associated with retina/heart morphogenesis under selection in wild ancestors, are ln(observed/expected derived allele frequency), whereas the y-axis of *LYST* and *PER2*, which were under selection in domestic duck, are Hp. (**c**) Distribution of F_ST_ between the wild and domestic duck and Hp in the domestic duck, calculated in a 40 kb window with a sliding size of 20 kb along scaffolds that are arranged according their alignment to chicken autosomes. The horizontal black line indicates the threshold at F_ST_=0.3858 or Hp=0.2319 (mean of ln values - 4 standard deviations). Red dots indicate the location of outlier genes (4 standard deviations away from hp means) that were visualized in detail in (**b**). (**d**) Comparison in luciferase activity of major alleles of MRPS27 and EFNA5 in wild (spot-billed and mallard) and domestic (Beijing) duck in DF1 (chicken embryonic fibroblast) cells. “**A**” and “**B**” represent wild duck and domestic duck, respectively. “*” P ≤ 0.05.

Subsequently, we performed a Gene Ontology (GO) analysis of 306 genes (12 of them detected in domestic selection were removed, Methods) in or closest to (extended 180 kb in both upstream and downstream) the selective sweep regions (SI Appendix) and highlighted 343 enriched categories related to 236 biological processes (Table S14, S15). Interestingly, the most frequent categories were related to a series of organs and systems, such as morphogenesis and development of the lung, heart, metanephros, aortic and cardiac muscle and functions of the blood circulatory system, respiratory system, peripheral nervous system and ventricular system development, and rhythmic excitation (Table 2). Because long-distance flight requires adaptive changes in body shape, energy metabolism and spatial navigation, we proposed that the selection of genes associated with the above functions might have enhanced the ability to fly long distances in the ancestor of mallard and spot-billed (SI Appendix). We further investigated functions of genes in putative selective sweep regions by surveying published literature and listed several genes that might be related to flight activity. The selection of the two genes *DNAJA1*, which stimulates ATP hydrolysis (Baaklini et al., 2012), and *MRPS27*, which aids in protein synthesis within mitochondria (Davies et al., 2012), might help supply increased energy for sustained flight. We also noted that *PARK2* was under selection when using a lower threshold (S=-3). *PARK2* promotes the autophagic degradation of dysfunctional depolarized mitochondria, potentially suggesting an adaptation to facilitate the degradation of damaged mitochondria caused by products of an elevated metabolic rate. Previous studies found that some genes play important roles in energy metabolism, such as ferrous iron transport (i.e., *SLC39A14*) or positive fever generation regulation (i.e., *PTGS2*) (Liuzzi, Aydemir, Nam, Knutson, & Cousins, 2006; McCarthy & Kosman, 2015). Positive selection of these genes in the early mallard and spot-billed duck ancestors might have enhanced their ability to transport oxygen in blood and adapt to cold and hypoxic environments at high altitudes during flight. Interestingly, we found that some genes involving retina and forebrain regionalization may implicate functions related to magnetic navigation ability (SI Appendix 4.7.1).

**Table 2.** Top 32 enriched biological process terms among genes under positive selection in the early spot-billed and mallard ancestors.

| Gene ontology term | P-value | Number | Size |
| --- | --- | --- | --- |
| positive regulation of tyrosine phosphorylation of Stat3 protein | 6.29E-05 | 4 | 12 |
| regulation of activin receptor signalling pathway | 0.000382 | 2 | 2 |
| positive regulation of cytosolic calcium ion concentration involved in phospholipase C-activating G-protein coupled signalling pathway | 0.000387 | 3 | 8 |
| detection of virus | 0.001131 | 2 | 3 |
| D-aspartate import | 0.001131 | 2 | 3 |
| metanephros morphogenesis | 0.002232 | 2 | 4 |
| L-glutamate import | 0.002232 | 2 | 4 |
| negative regulation of extrinsic apoptotic signalling pathway in absence of ligand | 0.003441 | 3 | 16 |
| positive regulation of granulocyte macrophage colony-stimulating factor production | 0.003672 | 2 | 5 |
| **respiratory system development** | **0.003672** | **2** | **5** |
| **determination of left/right symmetry** | **0.003701** | **5** | **53** |
| spinal cord motor neuron differentiation | 0.004119 | 3 | 17 |
| **lung morphogenesis** | **0.004119** | **3** | **17** |
| ionotropic glutamate receptor signalling pathway | 0.004872 | 3 | 18 |
| **ventricular cardiac muscle tissue morphogenesis** | **0.004872** | **3** | **18** |
| **dorsal/ventral pattern formation** | **0.005176** | **4** | **36** |
| negative regulation of peptidyl-threonine phosphorylation | 0.005437 | 2 | 6 |
| **serotonin metabolic process** | **0.005437** | **2** | **6** |
| **lung epithelium development** | **0.005437** | **2** | **6** |
| cytokine-mediated signalling pathway | 0.006291 | 4 | 38 |
| nucleotide-excision repair | 0.006613 | 3 | 20 |
| neutrophil chemotaxis | 0.006613 | 3 | 20 |
| negative regulation of epithelial cell proliferation | 0.006904 | 4 | 39 |
| **positive regulation of heart rate** | **0.007514** | **2** | **7** |
| inositol phosphate-mediated signalling | 0.007514 | 2 | 7 |
| **bud elongation involved in lung branching** | **0.007514** | **2** | **7** |
| **limb morphogenesis** | **0.009839** | **3** | **23** |
| long-chain fatty acid metabolic process | 0.00989 | 2 | 8 |
| dicarboxylic acid transport | 0.00989 | 2 | 8 |
| interleukin-6-mediated signalling pathway | 0.00989 | 2 | 8 |
| thrombin receptor signalling pathway | 0.00989 | 2 | 8 |
| **response to wounding** | **0.012416** | **3** | **25** |

In addition, we examined the functional activity of four mutations located in introns, UTRs (Untranslated Region) or downstream of *EFNA5*, *ISL1* and *MRPS27*. Three of the four sites with wild-type sequences showed significantly higher gene expression than the domestic genotype in a luciferase activity assay (Fig. 3d; Fig. S22; Table S16).

### Characterization on the genomic landscape of divergence between mallard and spot-billed consistent with very recent speciation

We estimated that mallard diverged from the spot-billed approximately 70 Kyr using two split models (Fig. 2a-b). This divergence time was consistent with a report counted using mtDNA sequences and less than a corresponding time (approximately 340 Kyrs-2 Myr) between two flycatcher species, which were studied extensively in speciation divergence (Ellegren et al., 2012; Nadachowska-Brzyska et al., 2013; Zhuravlev & Kulikova, 2014). In line with previous study, the fixed difference (d_f_) for 40 kb bins, which is an indication of species divergence, was used to detect the ‘divergence peak’ regions, which showed highly elevated divergence up to 50 times the genomic median. The findings presented a genomic landscape in sharp contrast to the case of flycatchers using the same criterion. Fewer d_f_s (4,386) with a larger proportion (80%) contained in the “divergence peak” are distributed in smaller genomic regions (0.6% of duck genome) than flycatchers (SI Appendix). This difference might because the two wild ducks are in an earlier stage of speciation than flycatchers (Nadachowska-Brzyska et al., 2013). Moreover, almost all “divergent peaks” were located in scaffolds homologous to the chicken “Z” chromosome. This finding may indicate an analogous situation to the flycatchers’ speciation (Discussion section; Appendix 4.7.2; Fig. S23).

Other genomic parameters (F_ST_ (fixation index), dxy (sequence divergence), π (nucleotide diversity) and Tajima’s D) for 40 kb bins showed a similar pattern as observed in many speciation studies. Our observations revealed highly heterogeneous genomic landscapes of differentiation when using elevated F_ST_ (4-fold standard deviations away from the mean, “differentiation islands”) as an indication of genomic divergence (Burri et al., 2015; Ellegren et al., 2012; Han et al., 2017; Lavretsky et al., 2015; Martin et al., 2013)(Fig. 4; Fig. S24). For example, the level of these parameters is significantly different between Z chromosomes and autosomes (Fig. 4a; Table S17). The nucleotide diversity and SNP density were reduced on the differentiation islands of autosomes (Fig. 4c-e; Table S18; SI Appendix 4.7.1). However, dxy was not elevated, and it was even reduced in most of the differentiation islands; This finding is consistent with the observation in flycatchers but different from the case of Darwin’s finches (Fig. S24; Burri et al., 2015; Han et al., 2017). Moreover, Tajima’s D of high differentiation was not reduced in mallard or spot-billed; this finding is different from observations in other species may also be an evidence of their very recent speciation (Burri et al., 2015; Han et al., 2017)(Discussion section; Fig. 4b; Fig. S24).

**Figure 4:**
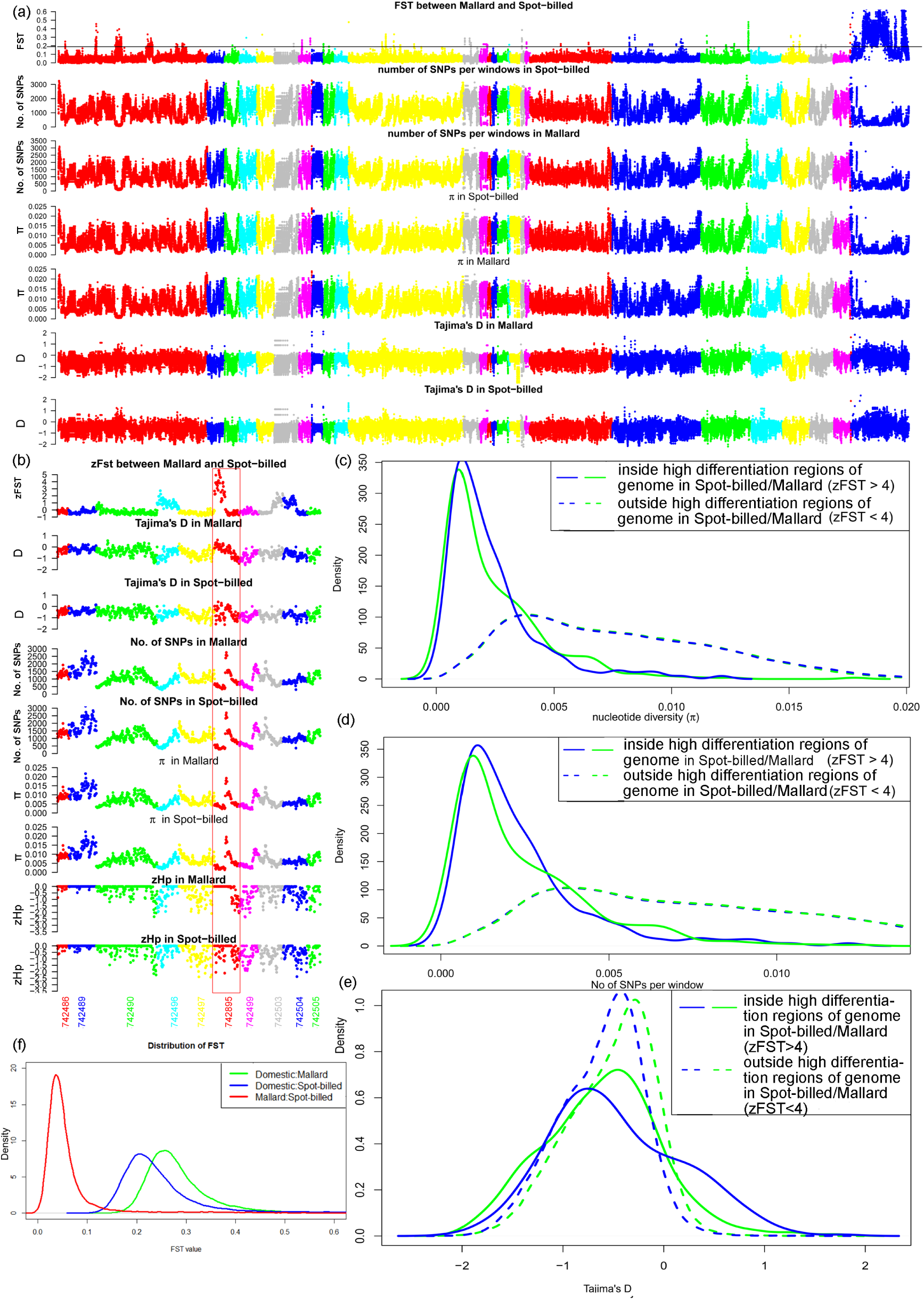
Genomic landscape of species divergence between spot-billed and mallard ducks. (**a**) Distribution of population genomic parameters of the Beijing duck assembly, in 40 kb windows with sliding 20 kb steps. The blue scaffolds in the right were referred to as duck Z scaffolds. The black horizontal line indicates the threshold at F_ST_=0.1888 (mean of F_ST_ + 4 standard deviations, referred to as Z(F_ST_)=4). The plots show the F_ST_ between spot-billed and mallard, number of SNPs per window, nucleotide diversity (π), and Tajima’s D for spot-billed and mallard. **(b)** Distribution of population genomic parameters along ten example scaffolds; this distribution is part of (a). **(c-e)** Distribution of nucleotide diversity (π), number of SNPs per window and Tajima’s D inside/outside high differentiation region autosomes (high differentiation regions, i.e., zF_ST_>=4, corresponding to bins above the horizontal line in (a) autosomes) in mallard and spot-billed ducks. **(f)** Distribution of pairwise F_ST_ for 40 kb windows with sliding 20 kb steps. The red line shows the plot in the first row of (**a**). Blue and green lines show the pairwise F_ST_ between spot-billed/mallard and domestic ducks (combining 8 domestic breeds).

### Identification of selective sweep regions in domestic duck

To detect genomic regions under domestic selection, we searched the duck genome for regions with reduced heterozygosity (Hp) and increased fixation index (F_ST_) using methods similar to the chicken and dog genome (Fig. S25; SI Appendix) (Axelsson et al., 2013; Rubin et al., 2010). We Z-transformed the Hp and F_ST_ distributions of autosomes and sex chromosomes separately and focused on putatively selective sweep regions falling at least four standard deviations away from the mean (Z(Hp)<-4 and Z(Z(F_ST_)>4) (Fig. 3c; Fig. S26-S27; SI Appendix 4.7.2). However, we found that only a small proportion of the selective sweep regions were predicted using both signals (Fig. S28; Table S19, S20), which may attribute to the long time difference in evolutionary history since their split (100-300 Kyr). Therefore, we focused on these 137 regions with extremely low levels of Hp (average length=63.6230 kb, average Hp=0.2100 for autosomes, 0.2470 for sex chromosomes) (SI Appendix). We found that 131 genes and 289 miRNAs were contained in 80 of these CDRs while 47 genes were closest to one of the 33 CDRs when we extended 180 kb both upstream and downstream of these 57 CDRs containing no annotated protein coding genes (Table S19, S21). Further GO analyses indicated that 162 genes in or closest to these 137 CDRs were enriched for 145 biological process categories (p-value<0.05), which represented developmental signalling; the nervous system; lipid, carbohydrate and protein metabolic processes; cell division; and immune responses (Table S22). Among them, two genes (*NLGN1* and *PER2*) are involved in the circadian sleep/wake cycle or associated with egg production (Nakao et al., 2007), three genes (*MRAP2, MC4R* and *RMI1*) are associated with the modulation of feeding behaviour or reducing food intake in response to dietary excess, and four genes (*HSD17B4, HEXB, B4GALT1* and *CYP19A1*) are related to sexual characteristics, male courtship behaviour or androgen metabolic processes (Dores & Garcia, 2015; Sebag, Zhang, Hinkle, Bradshaw, & Cone, 2013)(Fig. S29; Table S19, S21, S22).

Among these candidate domestication genes, we examined the functional activity of two mutations located in the introns of *GRIK1* and *LYST*. The domestic genotype of the *LYST* mutation showed significantly higher gene expression acidity than the wild genotype in a luciferase activity assay (Fig. 3d; Table S16)

### Absence introgression evidence of mallard and spot-billed further support their shallow divergence may not confounding by cross hybridization

Although introgression has been considered in our previous analyses and unlikely affected our inference, the Anseriformes is known for its propensity to interspecific hybridize. To further investigate the evidence of introgression between mallard and spot-billed we stratified the genomic divergence values of each window by ΔDAF (or absolute ΔDAF) (derived allele frequency difference between the two populations) bins. According to recent research on introgression issues in chimpanzee, the chance of derived alleles being introduced through gene flow increases with the frequency in the donor population. Gene flow should be introduced into receptor populations at low frequencies (de Manuel et al., 2016). Therefore, we may expect lower divergence values in these higher ΔDAF (absolute ΔDAF) bins because they represent a higher probability of introgression. We did find decreased genetic distance (FST, dxy) between mallard/spot-billed and domestic ducks in higher ΔDAF bins (Fig. S30a). However, both the FST and dxy values between mallard and spot-billed ducks increased with the ΔDAF (Fig. S30a blue lines). This observation found that evidence of introgression present between wild and domestic but not found between mallard and spot-billed. Furthermore, we found that the mean ΔDAF value did not show a significant difference between high differentiation regions and genomic background; this finding is consistent with recent research showing that gene flow is not the major factor in the formation of genomic islands (Table S23) (Han et al., 2017).

To further decipher the introgression effect and its influence on the evolutionary relationship inference of the domestic duck and the two wild ducks, we performed an ABBA-BABA test (assuming that the mallard, spot-billed, and domestic duck are P1, P2, and P3, respectively) using the Muscovy duck as an outgroup. The analysis suggested that the genetic relationship between mallard and spot-billed ducks are closer than it between the two wild species and the domestic duck (Fig. S31a). And we also found evidence of gene flow between the domestic and wild ducks that are all consistent with the above analysis (D-statistics: -0.2972; jackknife std: 0.4189; Fig. 1d, Fig. S15). To investigate whither gene flow has its effect on this phylogenetic tree, we used ΔDAF to test whether is there evidence that the sharing of this derived allele is caused by introgression. We stratified the number of BBAA (ABBA, BABA) sites by the ΔDAF bins between mallard and spot-billed duck populations (spot-billed and domestic, mallard and domestic, respectively). Most BBAA sites were distributed in the lower ΔDAF bins (-0.4 - 0.4) Because majority of SNPs distributed at lower ΔDAF bins. To eliminated the inhomogeneity distribution across the ΔDAF bins which is caused by phylogeny SNPs, we further measured the distribution of the BBAA ratio among all SNPs (i.e., polymorphic sites in spot-billed and domestic, mallard and domestic, mallard and spot-billed for ABBA, BABA, and BBAA, respectively). If introgression exists between mallard and spot-billed ducks (mallard and domestic, spot-billed and domestic), then the proportion of BBAA sites would likely increase with the ΔDAF (or absolute ΔDAF if negative). We did find that the proportion of BABA and ABBA sites increased with the ΔDAF, indicating that the derived allele sharing between wild (mallard or spot-billed) and domestic populations is partly due to introgression. This finding was consistent with previous analysis (Fig. 1d; Fig. S30). Whereas the distribution of the BBAA proportion across ΔDAF bins showed a different pattern than the BBAA rate, which showed only a small difference across ΔDAF bins, indicating lack of introgression between mallard and spot-billed ducks, and implying that the major reason for excessive derived allele sharing is due to phylogeny rather than introgression (Fig. S31b).

## DISCUSSION

Animal domestication are hot topics at least for two reasons: their important significance in the development of human civilization and supposed to be excellent models to understanding genetic changes underlying the striking morphological and behavioural changes observed in these animals (Carneiro et al., 2014; Frantz et al., 2015; Freedman et al., 2014; (Axelsson et al., 2013; Larson & Burger, 2013). However, the fundamental shift associated with the domestication are still unclear. Our research unexpected found the divergence time between domestic duck lineages and their closely wild relatives surprised longer than the origin of human civilization may provide an excellent case to understand domestication process. PCA, IBS tree, genetic structure, f3-statistic and species-specific alleles’ distribution were combined to inferred the new evolutionary relationship between mallard, spot-billed and domestic lineages, which has already implicated domestic lineage may originated unexpected long time ago. Although incomplete lineage sorting (ILS) and geneflow may confound the analyses. It unlike caused a genome-wide consequence (e.g. The genomic average genetic distance between the mallard and spot-billed ducks (mean F_ST_: 0.0637) significantly lower than that in mallard/spot-billed vs. domestic pairs (mean F_ST_: 0.2832, 0.2498) (Fig. 4b)). Furthermore, ∂a∂i and PSMC analyses that rely on different data sets and different aspects of data, derived divergence times coincided with each other, and indicated similar demography history. Together with our special analyses on introgression, we provided robustness inference about the evolutionary relationships. The scale of divergence time between domestic and wild was largely established on this topology and self-contained logic. It may suggest the domestication of duck was more a natural consequence related to environment and resource rather than singly human dominance (Irving-Pease et al., 2018).

Meanwhile, recent research also suggested that the critical changes at only a few domestication loci may not true, instead of many mutations of small effect, especially those frequency changes at regulatory regions play much more prominent role (Axelsson et al., 2013; Carneiro et al., 2014; Freedman et al., 2014; Rubin et al., 2012). Our long-term domestication process further supported this view and highlight the evolution context for those functional studies. Similar situation may be found in the flightless research that different mechanisms underlying different cases have been revealed (Campagna et al., 2019; Sackton et al., 2019). Before we discuss about flight loss, may be a better understanding on how flight’s acquired and evolved were also benefit. Our research revealed that the important phenotypic difference between wild and domestic duck, long distance flight abilities, may also a long-term evolutionary process that start from their divergence. It further implied the importance of evolutionary context on functional study and the complexity of flight evolution. Flight certainly seems to be linked to evolutionary success and a hard process (Swartz, 1998; Vargas, 2015). Birds’ flight originated from dinosaurian long time ago and made it through the extinction, and evolved to different extent of flight ability. They might give up flight when predators away and food resource plenty, most likely to occurred on islands and in aquatic species, and probably with little chance of evolving flight again (Armistead, 2014; Roff, 1994; Vargas, 2015). Most extinct birds are flightless might be the interpretation for the decline of domestic lineage and the maintain of wild population, especially when whether the domestic ducks lost flight or not successfully acquire it remain unclear (Fig. 2d).

Additionally, we found that duck domestication was accompanied by selection at genes affecting neuronal activities (e.g., *SH3GL2* (Wang et al., 2013)) and lekking behaviour (e.g., *HEXB, B4GALT1* and *HSD17B4*). Moreover, genes associated with lipid, carbohydrate, and protein metabolic processes and food intake control were under selection (e.g., *RMI1*, *MRAP2* and *MC4R* (Rubin et al., 2012)). In summary, an adaption to food-rich and artificial feeding environments with the development of agriculture may have contributed to duck domestication.

In addition, the rapid speciation in mallard and spot-billed, which demonstrates that speciation may occur within only 70 Kyr, provides an excellent resource for speciation studies. All “divergent peaks” located in scaffolds homologous to the ‘Z’ chicken chromosome are similar to the situation in flycatchers. Male plumage traits of the divergence species (mallard/spot-billed duck or collared/pied flycatcher) are quite different, and for flycatcher, both species-specific male plumage traits and species recognition are located on the Z chromosome (Saether et al., 2007; Saetre et al., 2003). Linked selection is suggested to dominantly drive the evolution of genomic differentiation rather than gene flow (Burri et al., 2015; Han et al., 2017; Rettelbach, Nater, & Ellegren, 2019). The lack of elevated dxy in “differentiation islands” and other analyses supported this suggestion. Furthermore, Tajima’s D is not reduced in differentiation islands potentially due to new strong positive selection (Fig. 4; Fig. S24). Tajima’s D is used to detect the selection of new mutations and correlated to the difference between π and the expectation of π, which is related to the number of SNPs (Misawa & Tajima, 1997). Due to the short divergence time, the mallard and spot-billed did not deposit many new mutations that increased population mutation rate (θ_w_) after selection (new mutations increase diversity at a low frequency thus would mainly increase the θ_w_ rather than nucleotide diversity (π) to reduce the Tajima’ D). Thus, the results in the allele frequency spectra did not skew towards rare alleles, whereas the collared and pied flycatchers (approximately 340 Kyr-2 Myr) and Darwin’s finches show reduced Tajima’s D in their genomic divergent islands due to the long-time interval after initial selection, which may be associated with the driving force of speciation (Han et al., 2017; Nadachowska-Brzyska et al., 2013).

In the end, we argue that our article has presented a distinct genetic resource between wild and domestic ducks. We also note that evidence in the genetic structure showed that mallard and spot-billed (especially the spot-billed duck) have interspecific similarities to domestic ducks (Fig. 1d and Fig. S15; especially when K=3,4). This finding may pose a significant threat to the genetic diversity of wild stocks.

## Availability of supporting data and materials

### Data availability

The genome assembly has been deposited in GenBank PRJNA392350. The duck, Taihu goose and Muscovy duck genome resequencing data are deposited under the BioProject PRJNA315043.

### Funding

The sequencing of the spot-billed and mallard genomes was funded by the National Basic Research Program of China (2013CB835200) and the National Natural Science Foundation of China (31471176). The resequencing of whole genomic sequences was funded by the National High Technology Research and Development Program of China (2010AA10A109) and the National Key Technology Support Program of China (2017SKLAB06-2).

### Author contributions

N.L. designed the project. Z.L.L, H.F.L., L.C., H.B.C., W.T.S., E.G.R and R.L collected and purified DNA samples. Q.W.L., Y.Z, G.Y.F., H.F.L., Y.S.S., W.B.C., X.L. and X.X. performed genome assembly, gene annotation and gene family evolution of the spot-billed and mallard genomes. R.L performed population genetics analysis. E.G.R. and X.X.W. performed luciferase activity analysis. R.L. and Q.W.L. wrote the manuscript. R.L. revised the manuscript.

## Acknowledgements

We would like to thank Dr Xiaojun Yang (Kunming Institute of Zoology) for identifying the wild duck species. We also thank Dr Minghui Chen, Dr Yiqiang Zhao, Prof. Xiaoxiang Hu, and Dr Jia Li from China Agricultural University; Prof. Zhonghua Zhang from the Chinese Academy of Agricultural Sciences; and Prof. Mingzhou Li from Sichuan Agricultural University for their helpful discussions and comments.

## References

Albarella, U. (2005). Alternate fortunes? The role of domestic ducks and geese from Roman to Medieval times in Britain. Documenta Archaeobiologiae III. Feathers, Grit and Symbolism (ed. by G. Grupe & J. Peters), 249–58.

Alexander, D. H., Novembre, J., & Lange, K. (2009). Fast model-based estimation of ancestry in unrelated individuals. Genome Res, 19(9), 1655–1664. doi:10.1101/gr.094052.109

Almathen, F., Charruau, P., Mohandesan, E., Mwacharo, J. M., Orozco-terWengel, P., Pitt, D., … Burger, P. A. (2016). Ancient and modern DNA reveal dynamics of domestication and cross-continental dispersal of the dromedary. Proc Natl Acad Sci U S A, 113(24), 6707–6712. doi:10.1073/pnas.1519508113

Andersson, L. (2001). Genetic dissection of phenotypic diversity in farm animals. Nat Rev Genet, 2(2), 130–138. doi:10.1038/35052563

Axelsson, E., Ratnakumar, A., Arendt, M. L., Maqbool, K., Webster, M. T., Perloski, M., … Lindblad-Toh, K. (2013). The genomic signature of dog domestication reveals adaptation to a starch-rich diet. Nature, 495(7441), 360–364. doi:10.1038/nature11837

Baaklini, I., Wong, M. J., Hantouche, C., Patel, Y., Shrier, A., & Young, J. C. (2012). The DNAJA2 substrate release mechanism is essential for chaperone-mediated folding. J Biol Chem, 287(50), 41939–41954. doi:10.1074/jbc.M112.413278

Botigue, L. R., Song, S., Scheu, A., Gopalan, S., Pendleton, A. L., Oetjens, M., … Veeramah, K. R. (2017). Ancient European dog genomes reveal continuity since the Early Neolithic. Nat Commun, 8, 16082. doi:10.1038/ncomms16082

Burri, R., Nater, A., Kawakami, T., Mugal, C. F., Olason, P. I., Smeds, L., … Ellegren, H. (2015). Linked selection and recombination rate variation drive the evolution of the genomic landscape of differentiation across the speciation continuum of Ficedula flycatchers. Genome Res, 25(11), 1656–1665. doi:10.1101/gr.196485.115

Burga, A., Wang, W., Ben-David, E., Wolf, P. C., Ramey, A. M., Verdugo, C., … Kruglyak, L. (2017). A genetic signature of the evolution of loss of flight in the Galapagos cormorant. Science, 356(6341). doi:10.1126/science.aal3345

Campagna, L., McCracken, K. G., & Lovette, I. J. (2019). Gradual evolution towards flightlessness in steamer ducks. Evolution, 73(9), 1916–1926. doi:10.1111/evo.13758

Carneiro, M., Rubin, C. J., Di Palma, F., Albert, F. W., Alfoldi, J., Barrio, A. M., … Andersson, L. (2014). Rabbit genome analysis reveals a polygenic basis for phenotypic change during domestication. Science, 345(6200), 1074–1079. doi:10.1126/science.1253714

Cruickshank, T. E., & Hahn, M. W. (2014). Reanalysis suggests that genomic islands of speciation are due to reduced diversity, not reduced gene flow. Molecular Ecology, 23(13), 3133–3157. doi:10.1111/mec.12796

Davies, S. M., Lopez Sanchez, M. I., Narsai, R., Shearwood, A.-M. J., Razif, M. F., Small, I. D., … Filipovska, A. (2012). MRPS27 is a pentatricopeptide repeat domain protein required for the translation of mitochondrially encoded proteins. FEBS letters, 586(20), 3555–3561.

DePristo, M. A., Banks, E., Poplin, R., Garimella, K. V., Maguire, J. R., Hartl, C., … Daly, M. J. (2011). A framework for variation discovery and genotyping using next-generation DNA sequencing data. Nat Genet, 43(5), 491–498. doi:10.1038/ng.806

de Manuel, M., Kuhlwilm, M., Frandsen, P., Sousa, V. C., Desai, T., Prado-Martinez, J., … Marques-Bonet, T. (2016). Chimpanzee genomic diversity reveals ancient admixture with bonobos. Science, 354(6311), 477–481. doi:10.1126/science.aag2602

Dores, R. M., & Garcia, Y. (2015). Views on the co-evolution of the melanocortin-2 receptor, MRAPs, and the hypothalamus/pituitary/adrenal-interrenal axis. Mol Cell Endocrinol, 408, 12–22. doi:10.1016/j.mce.2014.12.022

Ellegren, H., Smeds, L., Burri, R., Olason, P. I., Backstrom, N., Kawakami, T., … Wolf, J. B. (2012). The genomic landscape of species divergence in Ficedula flycatchers. Nature, 491(7426), 756–760. doi:10.1038/nature11584

Frantz, L. A., Mullin, V. E., Pionnier-Capitan, M., Lebrasseur, O., Ollivier, M., Perri, A., … Larson, G. (2016). Genomic and archaeological evidence suggest a dual origin of domestic dogs. Science, 352(6290), 1228–1231. doi:10.1126/science.aaf3161

Frantz, L. A., Schraiber, J. G., Madsen, O., Megens, H. J., Cagan, A., Bosse, M., … Groenen, M. A. (2015). Evidence of long-term gene flow and selection during domestication from analyses of Eurasian wild and domestic pig genomes. Nat Genet, 47(10), 1141–1148. doi:10.1038/ng.3394

Freedman, A. H., Gronau, I., Schweizer, R. M., Ortega-Del Vecchyo, D., Han, E., Silva, P. M., … Novembre, J. (2014). Genome sequencing highlights the dynamic early history of dogs. PLoS Genet, 10(1), e1004016. doi:10.1371/journal.pgen.1004016

Green, R. E., Krause, J., Briggs, A. W., Maricic, T., Stenzel, U., Kircher, M., … Paabo, S. (2010). A draft sequence of the Neandertal genome. Science, 328(5979), 710–722. doi:10.1126/science.1188021

Groenen, M. A., Archibald, A. L., Uenishi, H., Tuggle, C. K., Takeuchi, Y., Rothschild, M. F., … Megens, H.-J. (2012). Analyses of pig genomes provide insight into porcine demography and evolution. Nature, 491(7424), 393–398.

Gutenkunst, R. N., Hernandez, R. D., Williamson, S. H., & Bustamante, C. D. (2009). Inferring the joint demographic history of multiple populations from multidimensional SNP frequency data. PLoS Genet, 5(10), e1000695. doi:10.1371/journal.pgen.1000695

Han, F., Lamichhaney, S., Grant, B. R., Grant, P. R., Andersson, L., & Webster, M. T. (2017). Gene flow, ancient polymorphism, and ecological adaptation shape the genomic landscape of divergence among Darwin’s finches. Genome Res, 27(6), 1004–1015. doi:10.1101/gr.212522.116

Harr, B. (2006). Genomic islands of differentiation between house mouse subspecies. Genome Research, 16(6), 730–737.

Huang, Y., Li, Y., Burt, D. W., Chen, H., Zhang, Y., Qian, W., … Li, J. (2013). The duck genome and transcriptome provide insight into an avian influenza virus reservoir species. Nature genetics, 45(7), 776–783.

Huang, Y., Zhao, Y., Haley, C. S., Hu, S., Hao, J., Wu, C., & Li, N. (2006). A genetic and cytogenetic map for the duck (Anas platyrhynchos). Genetics, 173(1), 287–296. doi:10.1534/genetics.105.053256

Irving-Pease, E. K., Frantz, L. A. F., Sykes, N., Callou, C., & Larson, G. (2018). Rabbits and the Specious Origins of Domestication. Trends Ecol Evol, 33(3), 149–152. doi:10.1016/j.tree.2017.12.009

Larson, G., & Fuller, D. Q. (2014). The Evolution of Animal Domestication. Annual Review of Ecology, Evolution, and Systematics*, Vol* 45, *45*, 115–136. doi:10.1146/annurev-ecolsys-110512-135813

Lavretsky, P., Dacosta, J. M., Hernandez-Banos, B. E., Engilis, A., Jr., Sorenson, M. D., & Peters, J. L. (2015). Speciation genomics and a role for the Z chromosome in the early stages of divergence between Mexican ducks and mallards. Mol Ecol, 24(21), 5364–5378. doi:10.1111/mec.13402

Lavretsky, P., McCracken, K. G., & Peters, J. L. (2014). Phylogenetics of a recent radiation in the mallards and allies (Aves: Anas): inferences from a genomic transect and the multispecies coalescent. Mol Phylogenet Evol, 70, 402–411. doi:10.1016/j.ympev.2013.08.008

Li, H., & Durbin, R. (2011). Inference of human population history from individual whole-genome sequences. Nature, 475(7357), 493–496. doi:10.1038/nature10231

Liuzzi, J. P., Aydemir, F., Nam, H., Knutson, M. D., & Cousins, R. J. (2006). Zip14 (Slc39a14) mediates non-transferrin-bound iron uptake into cells. Proc Natl Acad Sci U S A, 103(37), 13612–13617. doi:10.1073/pnas.0606424103

Lohmueller, K. E., Bustamante, C. D., & Clark, A. G. (2010). The effect of recent admixture on inference of ancient human population history. Genetics, 185(2), 611–622. doi:10.1534/genetics.109.113761

Madge, S., & Burn, H. (1988). Waterfowl: an identification guide to the ducks, geese, and swans of the world: Houghton Mifflin.

Martin, S. H., Dasmahapatra, K. K., Nadeau, N. J., Salazar, C., Walters, J. R., Simpson, F., … Jiggins, C. D. (2013). Genome-wide evidence for speciation with gene flow in Heliconius butterflies. Genome Res, 23(11), 1817–1828. doi:10.1101/gr.159426.113

McCarthy, R. C., & Kosman, D. J. (2015). Iron transport across the blood–brain barrier: development, neurovascular regulation and cerebral amyloid angiopathy. Cellular and molecular life sciences, 72(4), 709–727.

Misawa, K., & Tajima, F. (1997). Estimation of the amount of DNA polymorphism when the neutral mutation rate varies among sites. Genetics, 147(4), 1959–1964.

Nadachowska-Brzyska, K., Burri, R., Olason, P. I., Kawakami, T., Smeds, L., & Ellegren, H. (2013). Demographic divergence history of pied flycatcher and collared flycatcher inferred from whole-genome re-sequencing data. PLoS Genet, 9(11), e1003942. doi:10.1371/journal.pgen.1003942

Nadachowska-Brzyska, K., Li, C., Smeds, L., Zhang, G., & Ellegren, H. (2015). Temporal Dynamics of Avian Populations during Pleistocene Revealed by Whole-Genome Sequences. Curr Biol, 25(10), 1375–1380. doi:10.1016/j.cub.2015.03.047

Nakao, N., Yasuo, S., Nishimura, A., Yamamura, T., Watanabe, T., Anraku, T., … Yoshimura, T. (2007). Circadian clock gene regulation of steroidogenic acute regulatory protein gene expression in preovulatory ovarian follicles. Endocrinology, 148(7), 3031–3038. doi:10.1210/en.2007-0044

Ni Leathlobhair, M., Perri, A. R., Irving-Pease, E. K., Witt, K. E., Linderholm, A., Haile, J., … Frantz, L. A. F. (2018). The evolutionary history of dogs in the Americas. Science, 361(6397), 81–85. doi:10.1126/science.aao4776

Reich, D., Thangaraj, K., Patterson, N., Price, A. L., & Singh, L. (2009). Reconstructing Indian population history. Nature, 461(7263), 489–494. doi:10.1038/nature08365

Richards, J. (2017). www.waterfowl.org.uk. Retrieved from http://www.waterfowl.org.uk/index.html

Rodgers, J. L. (1999). The Bootstrap, the Jackknife, and the Randomization Test: A Sampling Taxonomy. Multivariate Behav Res, 34(4), 441–456. doi:10.1207/S15327906MBR3404_2

Rubin, C. J., Megens, H. J., Martinez Barrio, A., Maqbool, K., Sayyab, S., Schwochow, D., … Andersson, L. (2012). Strong signatures of selection in the domestic pig genome. Proc Natl Acad Sci U S A, 109(48), 19529–19536. doi:10.1073/pnas.1217149109

Rubin, C. J., Zody, M. C., Eriksson, J., Meadows, J. R., Sherwood, E., Webster, M. T., … Andersson, L. (2010). Whole-genome resequencing reveals loci under selection during chicken domestication. Nature, 464(7288), 587–591. doi:10.1038/nature08832

Saether, S. A., Saetre, G. P., Borge, T., Wiley, C., Svedin, N., Andersson, G., … Qvarnstrom, A. (2007). Sex chromosome-linked species recognition and evolution of reproductive isolation in flycatchers. Science, 318(5847), 95–97. doi:10.1126/science.1141506

Saetre, G. P., Borge, T., Lindroos, K., Haavie, J., Sheldon, B. C., Primmer, C., & Syvanen, A. C. (2003). Sex chromosome evolution and speciation in Ficedula flycatchers. Proc Biol Sci, 270(1510), 53–59. doi:10.1098/rspb.2002.2204

Sebag, J. A., Zhang, C., Hinkle, P. M., Bradshaw, A. M., & Cone, R. D. (2013). Developmental control of the melanocortin-4 receptor by MRAP2 proteins in zebrafish. Science, 341(6143), 278–281. doi:10.1126/science.1232995

Seehausen, O., Butlin, R. K., Keller, I., Wagner, C. E., Boughman, J. W., Hohenlohe, P. A., … Widmer, A. (2014). Genomics and the origin of species. Nat Rev Genet, 15(3), 176–192. doi:10.1038/nrg3644

Stern, D. L. (2000). Evolutionary developmental biology and the problem of variation. Evolution, 54(4), 1079–1091.

Steve, M. (1992). Waterfowl: an identification guide to the ducks, geese and swans of the world. Houthton mifflin in harcout, 3.

Wang, G. D., Zhai, W., Yang, H. C., Fan, R. X., Cao, X., Zhong, L., … Zhang, Y. P. (2013). The genomics of selection in dogs and the parallel evolution between dogs and humans. Nat Commun, 4, 1860. doi:10.1038/ncomms2814

Wang, G. D., Zhai, W., Yang, H. C., Wang, L., Zhong, L., Liu, Y. H., … Zhang, Y. P. (2016). Out of southern East Asia: the natural history of domestic dogs across the world. Cell Res, 26(1), 21–33. doi:10.1038/cr.2015.147

Warmuth, V., Eriksson, A., Bower, M. A., Barker, G., Barrett, E., Hanks, B. K., … Manica, A. (2012). Reconstructing the origin and spread of horse domestication in the Eurasian steppe. Proc Natl Acad Sci U S A, 109(21), 8202–8206. doi:10.1073/pnas.1111122109

Xiang, H., Gao, J., Yu, B., Zhou, H., Cai, D., Zhang, Y., … Zhao, X. (2014). Early Holocene chicken domestication in northern China. Proc Natl Acad Sci U S A, 111(49), 17564–17569. doi:10.1073/pnas.1411882111

Zhuravlev, Y. N., & Kulikova, I. V. (2014). Waterfowl Population Structure: Phylogeographic Inference. Achievements in the Life Sciences, 8(2), 123–127.

Armistead, H. T. (2014). Flight Ways: Life and Loss at the Edge of Extinction. Library Journal, 139(14), 138–138.

Larson, G., & Burger, J. (2013). A population genetics view of animal domestication. Trends Genet, 29(4), 197–205. doi:10.1016/j.tig.2013.01.003

Rettelbach, A., Nater, A., & Ellegren, H. (2019). How Linked Selection Shapes the Diversity Landscape in Ficedula Flycatchers. Genetics, 212(1), 277–285. doi:10.1534/genetics.119.301991

Roff, D. A. (1994). The Evolution of Flightlessness - Is History Important. Evolutionary Ecology, 8(6), 639–657. doi:Doi 10.1007/Bf01237847

Sackton, T. B., Grayson, P., Cloutier, A., Hu, Z., Liu, J. S., Wheeler, N. E., … Edwards, S. V. (2019). Convergent regulatory evolution and loss of flight in paleognathous birds. Science, 364(6435), 74–78. doi:10.1126/science.aat7244

Swartz, S. (1998). Taking wing: Archaeopteryx and the evolution of bird flight. Science, 281(5375), 355–356. doi:DOI 10.1126/science.281.5375.355

Vargas, A. (2015). Flying Dinosaurs: How Fearsome Reptiles Became Birds. Journal of Field Ornithology, 86(2). doi:10.1111/jofo.12103

Zhang, Z., Jia, Y., Almeida, P., Mank, J. E., van Tuinen, M., Wang, Q., … Qu, L. (2018). Whole-genome resequencing reveals signatures of selection and timing of duck domestication. Gigascience, 7(4). doi:10.1093/gigascience/giy027

Zhou, Z., Li, M., Cheng, H., Fan, W., Yuan, Z., Gao, Q., … Jiang, Y. (2018). An intercross population study reveals genes associated with body size and plumage color in ducks. Nat Commun, 9(1), 2648. doi:10.1038/s41467-018-04868-4

