## supplemental text for "Genomic analyses reveal the origin of domestic ducks and identify different genetic underpinnings of wild ducks"

**Supporting information for: Genomic analyses reveal the origin of domestic ducks and identify different genetic underpinnings of wild ducks (SI Appendix)**

Rui Liu[1,*], Weiqing Liu[2,3,*], Enguang Rong[1], Lizhi Lu[4], Huifang Li[5], Li Chen[4], Yong Zhao[3,6], Huabin Cao[7], Wenjie Liu[1], Chunhai Chen[2], Guangyi Fan[2,6,8], Weitao Song[6], Huifang Lu[3], Yingshuai Sun[3], Wenbin Chen[2,9], Xin Liu[2,6,9], Xun Xu[2,6,9], Ning Li[1,#]

[1]State Key Laboratory for Agrobiotechnology, China Agricultural University, Beijing, 100094, China. [2]BGI-Shenzhen, Shenzhen 518083, China. [3]BGI-Wuhan, Wuhan 430075, China. [4]Institute of Animal Sciences and Veterinary Medicine, Zhejiang Academy of Agricultural Sciences, Hangzhou 310021, China. [5]Institute of Poultry Science of Jiangsu, Yangzhou 225125, China. [6]BGI-Qingdao, Qingdao 266555, China. [7]Jiangxi Provincial Key Laboratory for Animal Health, Institute of Animal Population Health, College of Animal Science and Technology, Jiangxi Agricultural University, Nanchang 330045, China. [8]State Key Laboratory of Quality Research in Chinese Medicine, Institute of Chinese Medical Sciences, University of Macau, Macao, China. [9]China National GeneBank-Shenzhen

[*]These authors contributed equally to this work

### Contents Tables

Supplementary text ... 6
1. Genome sequencing and assembly ... 6
1.1 Genome sequencing ... 7
1.2 Raw data filtering and genome size estimation ... 7
1.3 Genome assembly and super-scaffold construction ... 8
1.4 Quality evaluation of the genome assembly ... 8
2. Repeat prediction, gene structure and gene functional annotation ... 8
2.1 Transposable elements annotation ... 8
2.2 Variation of CR1 (chicken repeat 1) transposon among ducks ... 9
2.3 Gene structural annotation ... 9
2.4 Gene functional annotation ... 10
3. Evolutionary analysis ... 10
3.1 Gene family clustering ... 10
3.2 Phylogenetic tree reconstruction and divergent time estimation ... 10
3.3 Segmental duplications ... 11
4. Resequencing and population genomic analyses ... 11
4.1 Genome sequencing, Polymorphism identification, accuracy verification and its combine between populations ... 11
4.2 Domestic duck shared a comparative number of SNPs with the mallard duck and spot-billed duck ... 12
4.3 Estimation of nucleotide diversity and population mutation rate ... 12
4.4 Analysis of linkage disequilibrium (LD), identical by state (IBS), principal component analysis (PCA), f3-statistics and population structure ... 12
4.5 PCA, genetic structure and PSMC analysis ... 12
4.6 ∂a∂i simulation ... 14
4.6.1 Devising strategy to figure out the splitting order of domestic lineage and the two wild duck species. ... 14
4.6.2 Data sets and infrastructure ... 16
4.6.3 Comparisons with previous studies ... 17
4.7 Selection analyses and genomic islands ... 18
4.7.1 Detail functional analysis on candidate selected genes in early ancestor of mallard and spot-billed ducks. ... 18
4.7.2 Detail analysis on genomic islands of mallard and spot-billed duck divergence. ... 19
4.7.3 Detail analysis on identification of candidate domestication regions ... 20
4.7.4 Evolutionary relationship of domestic duck, mallard wild duck and spot-billed wild duck ... 22
4.8 Estimation of genetic diversity and distance ... 22
4.9 select candidate casual mutation to verify its functional activity ... 22

#### Supplementary text

##### 1. Genome sequencing and assembly

We generated 262.45 Gb (~209-fold) pair-end reads of a female mallard duck and 275.20 Gb pair-end reads (~220-fold) of a female spot-billed duck, respectively (Table S2,Fig. S2-S3). Using methods similar to those applied to the Beijing duck genome (Yinhua Huang et al., 2013), we generated a high-quality mallard assembly consisting of 61,591 scaffolds and covering 1.27 Gb, and estimated this assembly having a heterozygosity rate $3.61 \times 10^{-3}$ higher than the corresponding in Beijing duck genome ($2.40 \times 10^{-3}$). The contig N50 and scaffold N50 values of the mallard genome assembly were 38.27 Kb and 2.49 Mb, respectively (Table 1; Table S3). We also generated a high-quality spot-billed genome assembly containing 67,685 scaffolds and covering 1.31 Gb. Similarly, we found that spot-billed assembly had a high heterozygosity rate ($3.54 \times 10^{-3}$). The contig N50 and scaffold N50 values of the spot-billed assembly were 33.06 Kb and 2.05 Mb, respectively (Table 1; Table S3). We then estimated coverage of the mallard duck and spot-billed duck assemblies with alignment to seven finished BACs (completed independently using Sanger sequencing technology) and 319,996 duck ESTs assembled in our previous duck genome project using BLAT (Yinhua Huang et al., 2013; Huang et al., 2011). This analysis suggested that 7 BACs covering 640 Kb on chromosomes 1, 3 and 4 were aligned over more than 92% of their lengths (Fig. S4-S5, Table S4), and greater than 98% of 319,996 ESTs were aligned to both the mallard duck and spot-billed duck assemblies (Table S5-S6).

We annotated transposable elements (TE) of the mallard duck and spot-billed duck assemblies using a combined pipeline. This effort found that the mallard duck and spot-billed duck assemblies contained ~136 Mb and ~139 Mb TE sequences, respectively, accounting for about 10% of their assemblies, respectively (Table S7). Among those TE elements, the LINE (long interspersed elements) type CR1 (chicken repeat 1) transposons are of the most abundance, a situation similar as ones in Beijing duck and chicken. Phylogenetic analysis of CR1 in mallard duck, spot-billed duck, Beijing duck and chicken revealed two duck-specific CR1 clades (clade 1 and 3), in which large variation of copy numbers were shown between the two wild ducks and Beijing duck. This implies that, as a main class of TE, CR1 evolved rapidly among different avian species and distinct duck lineages. (Fig. 1a; Fig. S6).

We predicted 21,056 and 21,123 protein-coding genes, which constitutes approximately 2.21% of the mallard genome and 2.13% of the spot-billed genome, using the BGI pipelines (Table S8). These numbers are slightly larger than the corresponding in the domestic duck (Beijing duck) genome (20,629 protein-coding genes) (Table 1). Of the 21,056 mallard genes, 12,817 were mapped to categories established by the Gene Ontology (GO) project, 15,878 had orthologs in the Kyoto

Encyclopedia of Genes and Genomes (KEGG) database, and 14,894 were supported by the duck ESTs produced in our previous duck genome project (Fig. S7). These numbers are similar to these of the spot-billed, where 12,815 genes were mapped to GO project, 15,803 had orthologs in KEGG database and 14,827 were supported by the duck ESTs (Fig. S8).

We then constructed gene families using the above mallard duck and spot-billed reference gene sets, our previous Beijing duck gene set, together with a combined outgroup gene sets from chicken, turkey, flycatcher, ostrich, mouse and human (Fig. S9, Table S9). Interestingly, two wild ducks were similar and their ancestors were clustered to the domestic (Beijing) duck (Fig. 1b). We further examined large-scale differences in gene complements between one domestic duck and two wild ducks. We found that 1,305 gene families were specific to wild duck and this number was larger than the corresponding of Beijing duck (497 gene families) (Fig. S10). Among 19,405 gene families detected in the above three duck genomes, 25 were significantly expanded in wild duck. GO enrichment analysis showed that those expanded genes were significantly over represented in biological processes like chromatin organization, cell-cell adhesion, ATP synthesis and immune response. This implies enhanced genome regulation, energy metabolic activity and response to stimulation along with the speciation of the two wild ducks, which might be beneficial to their adaption to new environments (Table S10; Fig. 1c-d).

##### 1.1 Genome sequencing

We extracted genomic DNA with standard molecular biology techniques from blood and random fragmented DNA for library construction. For short insert size libraries, 5 μg of DNA were fragmented to 250-800bp, end-repaired, A-tailed and ligated to Illumina paired-end adapters (Illumina). The ligated fragments were size selected at 250, 350 and 800 bp on agarose gel and amplified by PCR to yield the corresponding short insert size libraries. For mate-pair library construction, 20-40μg DNA were sheared to the desired insert size using nebulization for 2 kb or HydroShear (Covaris) for 5 kb, 10 kb and 20 kb. Then the DNA fragments were end-repaired using biotinylated nucleotide analogues and circularized by intramolecular ligation Circular DNA molecules were sheared with Adaptive Focused Acoustic (Covaris) to an average size of 500 bp. Biotinylated fragments were purified with Dynabeads® M-280 Streptavidin beads (Invitrogen), end-repaired, A-tailed and ligated to Illumina paired-end adapters, PCR amplification and agarose gel electrophoresis for size-selection. Then sequence all these libraries with Illumina Hiseq2000 platform.

##### 1.2 Raw data filtering and genome size estimation

To reduce the effect of sequencing error or low-quality reads for genome assembly, we took a series of checking and filtering steps on raw reads. We filtered: 1) Reads having an ‘N’ over 10% of their length; 2) Reads with more than 40 bp low quality base (quality score ≤ 7); 3) Reads with more than 10 bp adapter sequences (allowing ≤ 3 bp mismatches); 4) Short insert size paired-end reads that were overlapped (≥ 10bp); 5) Redundant duplicated reads generated by PCR amplification in the library construction (Read1 and Read2 of two pairs of paired-end reads were identical, respectively). In total, 275.20 and 262.45 Gb of data were kept for D2B and D2L *de novo* genome assembly respectively.

Genome size can be estimated using K-mer analysis method, the principle is briefly depicted as follows: A K-mer is an artificial sequence division of K bp iteratively from sequencing reads. A read with L bp in length composes (L-K+1) K-mers. The frequency of each K-mer can then be calculated from the input data. The resulted K-mer frequencies should generally follow a Poisson distribution, except for a high proportion of low frequencies due to sequencing errors. The genome size G can be estimated as G= K_num/K_expect, where K_num is the total number of K-mer, and K_expect is the expected value of the poisson distribution. We performed K-mer analysis (K=17) using about 21-27X filtered data, and estimated the mallard and spot-billed genome size to be about 1.15Gb and 1.14Gb, respectively (Fig. S1).

##### 1.3 Genome assembly and super-scaffold construction

At first, we assembled the two genomes using SOAP *de novo*(Green et al., 2010) for contig and scaffold construction. Detailed assembly steps and the data used in each step: 1) Constructed *de Bruijn* graph with reads from the short insert size libraries using K-mer size 29 bp, which means the parameter –K is set as 29. 2) Simplified the *de Bruijn* graph for contigs construction by removal tips, merging bubbles, removal the low coverage of the connection and resolve the small repeats. 3) Constructed scaffolds with short pair-end and mate-pair libraries step by step. 4) Filled in gaps inside the constructed scaffolds with gap closure step.

##### 1.4 Quality evaluation of the genome assembly

We made use of known EST and BAC sequences to evaluate the accuracy of genome assembly. We mapped 7 BACs(Huang et al., 2011) and 319,996 ESTs of Beijing duck onto the two assemblies using BLAT (Kent, 2002) with default parameters. We found that, for those 7 BACs totally spanning 640 Kb, more than 92% of the spanning were aligned well to the assemblies (Fig. S3 and S4). In addition, around 94% of the ESTs were mapped to individual scaffold of the assemblies (Table S6).

Excessive heterozygosity at regional homologous chromosomes usually lead to failure of collapsing the two haplotype sequences, and thus results in redundant sequences in the assembly. To determine whether there is any case of redundant sequences in our assemblies, we performed a self-to-self alignment of the assemblies. Then found the total length of the scaffolds with the coverage rate is more than 50% is 11.3Mb and 10.3Mb in female mallard and a female spot-billed genome, respectively. That analysis indicated there was almost no redundant sequence in the two assemblies and we masked these scaffolds with coverage rate more than 50% for further deep analysis.

#### 2. Repeat prediction, gene structure and gene functional annotation

##### 2.1 Transposable elements annotation

We annotated transposable elements (TE) using the following steps:
1) We first scanned for repeat elements using LTR_FINDER(Xu & Wang, 2007) with the default parameters. The resulted TE sequences were used as TE database for RepeatMasker(Chen, 2004) search in step 2.
2) We then used RepeatMasker to identify repeat elements against the Repbase database(Jurka et al., 2005) as well as the TE sequences founded in step1.
3) We further searched the assembly against the protein repetitive database provided in RepeatMasker using RepeatProteinMask.
4) At last, we combined all the three set of predictions (two from step2 and one from step3) to generated an integrated repeat annotation result for each genome assembly.
5) Tandem repeats were identified by Tandem repeat finder(Benson, 1999) using the defaults of “Match=2, Mismatch=7, Delta=7, PM=80, PI=10, Minscore=50, and MaxPeriod=12”.

##### 2.2 Variation of CR1 (chicken repeat 1) transposon among ducks

From the result of repetitive elements annotation in section 2.1, we found that CR1 (chick repeat 1), a subtype of LINE retrotransposon, accounts for both the majority of both TE content and variation of TE content among mallard, spot-billed duck, Beijing duck and chicken assembly. In fact, we found there were more than 90Mb (7.4%) of CR1 subtype repeat in two wild ducks, but only approximately 60Mb (5.5%) in Beijing duck. Therefore, we further studied the variation of CR1 among those four species in a perspective of phylogeny. We downloaded non-LTR retrovirus reverse transcriptase (RT) of chicken retrotransposon CR1, complete consensus sequence from NCBI. We extracted the 252 amino acids (aa) long RT_nLTR_like protein sequences from the consensus, and then mapped it to the four genome assemblies using tblastn. We identified 2579, 2294, 617, 1737 RT_nLTR_like domains in the four genome assemblies (evalue < 1e-5, identity >50%, alignment length >200bp), respectively. We then performed multiple sequence alignment of all the identified RT_nLTR_like domains using MUSCLE(Edgar, 2004) (version 3.8.31). Based on the alignment, a phylogenetic tree was subsequently reconstructed using FastTree.

##### 2.3 Gene structural annotation

First, we applied the homolog based gene prediction method using protein datasets of *Anas platyrhynchos, Gallus gallus, Struthio camelus, Meleagris gallopavo, and Ficedula albico*. All the protein sequences were firstly mapped to the assembly using BLASTN with the parameters as “-e 1e-5 –F F –m 8”. Then the aligned protein sequences were realigned to the assembly using GeneWise(Birney, Clamp, & Durbin, 2004) for accurate spliced alignments.
Second, we used Beijing Duck transcriptome reads to polish our target gene annotation. We mapped the transcriptome reads to the two wild duck genome assemblies using TopHat(Trapnell, Pachter, & Salzberg, 2009) with its default settings and then constructed transcripts using Cufflinks. The process is the same as one preformed in the Beijing Duck genome project(Y. Huang et al., 2013).

Finally, we combined all the gene models (five homologous gene models sets and the transcript gene models set) by GLEAN(Elsik et al., 2007) to generate an consensus gene set.

##### 2.4 Gene functional annotation

We aligned protein sequences of the final gene set to various protein databases with known functional knowledge, including InterPro(Hunter et al., 2009) (iprscan_4.7) (profilescan, blastprodom, hmmsmart, hmmpanther, hmmpfam, fprintscan, patternScan), Gene Ontology(Ashburner et al., 2000), Swiss-Prot(Schmitt, Gueguen, Desmarais, Bachere, & de Lorgeril, 2010), TrEMBL(Schmitt et al., 2010) and KEGG(Kanehisa & Goto, 2000) (release76), to identify conserved domains, putative biological functions and molecular pathways they may involve. In total, there are more than 87% of the annotated protein-coding gene models had at least one hit with known function (Table S6).

#### 3. Evolutionary analysis

Mallard and spot-billed ducks, together with five other avian species (*Gallus gallus, Struthio camelus, Meleagris gallopavo, Ficedula albico and Anas platyrhynchos*, which is the Beijing duck) and two mammals (*Homo sapiens*, *Mus musculus*), were used in evolutionary analysis.

##### 3.1 Gene family clustering

Protein-coding genes for *Gallus gallus, Struthio camelus, Meleagris gallopavo, and Ficedula albico* were downloaded from Ensembl release 80. The gene sets for *Anas platyrhynchos* was obtained from the BGI inner database. For gene loci with alternative splicing isoforms, only the transcript with the longest translation product was retained. We carried out an all-to-all alignment of all the collected protein sequences using BLASTP (E-value < 1E-7), and conjoined fragmental alignments using Solar(Yu, Zheng, Wang, Wang, & Su, 2006). Then a simplified version of Treefam(Ruan et al., 2008) methodology was used to cluster individual sequence into families based on the conjoined alignment (Table S11).

##### 3.2 Phylogenetic tree reconstruction and divergent time estimation

After gene family clustering, single copy genes were selected to reconstruct phylogenetic tree of those nine species. Multiple sequence alignment of each gene family was performed by MUSCLE(Edgar, 2004) (version 3.8.31). Phase1-base degenerate sites were extracted and concatenated to generate a super alignment matrix. We built phylogenetic tree using MrBayes(Huelsenbeck & Ronquist, 2001) which takes advantage of both codon-based and amino acid-based algorithms and adjusts them to the topology of the species tree, to form a more accurate consensus tree according to phase1-base degenerate site. Divergence time of species were estimated using molecular clock model implemented by PAML mcmctree(Z. Yang, 1997).

##### 3.3 Segmental duplications

Segmental duplications (SDs) are duplicated blocks of genomic DNA typically ranging in size from 1–200 kb(McPherson et al., 2001). To explore any difference in the content of SDs among two wild duck and Beijing duck genome, we detected SDs in their genome assembly using a whole-genome-alignment based method. We performed self-to-self whole genome alignment of these three assemblies using BLASTZ with the parameters: T=2 C=2 H=2000 Y=3400 L=6000 K=2200. Repetitive elements were masked in advance. We then filtered out self-alignment results (>0.85 similarity and >1000 aligned bp) and obtained more credible align block pairs. Finally, we realigned and filtered these block pairs (>0.9 similarity and >1000 aligned bp), producing a final version of SD regions in each genome. In summary, we identified 59.64 Mb (coving 4,367 genes) and 66.05 Mb (coving 4,864 genes) segmental duplications in mallard and spot-billed genome respectively, which are much larger than Beijing duck's 8.39 Mb (coving 1,133 genes). We further did pair-wise alignment among SDs detected in all these three assemblies to identify SDs specific to each of them.

#### 4. Resequencing and population genomic analyses

##### 4.1 Genome sequencing, Polymorphism identification, accuracy verification and its combine between populations

Genomic DNA libraries were prepared according to the manufacturer's instructions (Illumina). One DNA sample or pools of DNA samples from 10 individuals for each six duck breeds were used to generated pair-end reads with the Illumina technology (Table S1). Pair-end reads were mapped to the Beijing duck assembly (BGI_duck_1.0) using the bowtie2 with default parameters. After removing the reads which map to multiple place, we called SNPs using the Genome Analysis Toolkit (GATK) and filtered SNPs according to the coverage of resequencing data with thresholds of ≤ one third of the mean or ≥ three-fold of the mean and MQ value ≤ 28(DePristo et al., 2011)

We produce BAM file for each individual sample or population pool according its sequencing strategy (Table S11). VCF files for each population were using "UnifiedGenotyper" of GATK by taken all processed BAM files of individual samples as input, or taken pool BAM file of a population as input. And all BAM files were processed by "REMOVE_DUPLICATES", "use unique mapped reads" and "IndelRealigner". In the functional validation section, 10 SNP sits were selected to be used to experimental verification its functional activity. 9 of these 10 were true SNPs but one was INDEL instead. We think that these results could reflect the high accuracy of SNP calling in some extent. In the following selection analyses or ∂a∂i simulation, all SNPs are from regions that homologous to chicken autosomes. We combine the VCF files by using a way like outer join of SQL (Structured Query Language) (https://github.com/vitogump/life). For the variations that present in one

VCF but absent in the another one, we fill it as it fixed as ref allele when the coverage (information from BAM) of this site reached the threshold.

##### 4.2 Domestic duck shared a comparative number of SNPs with the mallard duck and spot-billed duck

Detailed analyses suggested that domestic ducks shared large numbers of SNPs (18,093,115) with wild duck, and hold comparative numbers of common SNPs with the mallard (15,904,435) and spot-billed (16,062,465) ducks. Moreover, we found that the numbers of mallard specific SNPs (heterozygous in mallard and homozygous in spot-billed) are comparative to the numbers of spot-billed specific SNPs (homozygous in mallard and heterozygous in spot-billed) in each of eight tested domestic duck breeds (Fig. S12).

##### 4.3 Estimation of nucleotide diversity and population mutation rate

We estimated nucleotide diversity ($\pi$) and Watterson's estimator $\Theta$ using the PoPoolation package, which corrects biases causing by pooling and sequencing errors, in 40 kb non-overlapping windows(Karlsson et al., 2007; Kofler et al., 2011). We required per position a minimum coverage of 13,13,13,6 and 6 a maximum coverage of 150,150,500,100 and 100 for mallard, spot-billed, Beijing, Shaoxing, and other 8 domestic population each, and set min-count and min-covered-fraction to be 3 and 0.6 for all.

##### 4.4 Analysis of linkage disequilibrium (LD), identical by state (IBS), principal component analysis (PCA), f3-statistics and population structure

LD level in Beijing, Shaoxing, mallard duck and spot-billed duck populations were measured through calculating the correlation coefficient ($r^2$) of alleles using the PLINK(Purcell et al., 2007).

The parameters were set as follows: --ld-window-kb 500 --ld-window 99999 --ld-window-r2 0 --r2. To reduce effect of LD, SNPs were pruning by LD and subsequently used for the PCA, f3-statistics and population structure analyses (Fig. S11). We constructed phylogenetic trees by using neighbor-joining method with MEGA based on IBS distance matrix data of all individuals calculated by the PLINK using the above sampling SNPs(Tamura et al., 2011). Principal component analysis (PCA) was performed using the GCTA64(J. Yang, Lee, Goddard, & Visscher, 2011). The f3-statistics were calculated using the ADMIXTOOLS and genetic structure was inferred using the admixture program.

##### 4.5 PCA, genetic structure and PSMC analysis

###### 4.5.1 Technical details

We use Beijing and Shaoxing ducks to illustrate the relationship with wild as a representation of domestic ducks when preforming analysis require individual samples such as genetic structure analysis:

When PCA analysis were performed on the unit of population genomic SNP, Fig. S14**b** shows that first principle component separate Muscovy duck (outgroup) with eight domestic duck populations and two wild duck (mallard and spot-billed). The second principle component separate wild with domestic populations, and Shaoxing, Beijing ducks were cluster together with all other domestic duck with no any special. Fig. S14**c** further shows that Beijing, Shaoxing and other 6 domestic population are adjacent to each other with almost equally "distance" and clustered according to their geographical distributions.

SNPs used in both PCA and genetic structure analysis were filtered using PLINK(Purcell et al., 2007) (--indep-pairwise 500 50 0.4 --maf 0.05). First, we want to know the genetic relationship of all 8 wild duck species and these domestic populations as mentioned in main text Fig. 2a, which provided evidence that mallard duck and spot-billed duck having a far more closely genetic relationship with domestic duck than other wild species. To eliminate the unequal simple size effect, assembles difference effect and autosomes/sex difference effect, we supplemented PCA analysis by using same sample size for each population, with both mallard duck assemble and Beijing duck assemble (Fig. S14). These results further supported our conclusion.

Second, in our genetic structure analysis, it is interesting that domestic lineage separated from wild duck first, and even subdivided into Beijing and Shaoxing previous than the mallard duck and spot-billed duck's subdivision. To further investigate this phenomenon and eliminate the coverage effect. We perform genetic structure analysis by using equal sample size and same coverage of each population with different populations combinations (Fig. S15). These conclusions are consistent.

Third, in our PSMC analysis, SNPs were filtered according to the coverage of resequencing data at their position with thresholds of 10- to 100-fold for Beijing, spot-billed and mallard and 5- to 80-fold for Shaoxing. Use the parameter "-N30 -t5 -r5 -p "4+30*2+4+6+10" ". When we cutting the coverage of Beijing individuals into the same with Shaoxing individuals, they show the same curve with Shaoxing ducks (Fig. S18). So, we propose that all domestic ducks should have the same PSMC curves.

Cutting sequence coverage was performed by samtools(H. Li et al., 2009) (samtools view -bhs).

##### 4.5.2 Monophyletic taxon of mallard and spot-billed

IBS tree root on different point or without root exhibit different clusters. Sometimes even Shaoxing ducks were not shown as a monophyletic taxon that is obviously not true (Fig. S12f,g,h). Meanwhile, all kinds shape of the tree didn't indicate mallards and spot-billed are monophyletic (Fig. 1b). We think this is because 1) mallards and spot-billed have a very close relationship 2) that almost can't distinguish through IBS tree. To confirm whether samples were properly assigned to their species. We noticed the IBS tree that mallard samples did cluster by the two places where we collected samples from---Fenghua and Hangzhou. And spot-billed were also clustered. With 2 or 3 exception individuals: No. 12 female mallard form Hangzhou were clustered with spot-billed and No. 22 male spot-billed were clustered with mallards (female mallards have similar look with spot-billed and male and female were also looks similar). Whereas, 1. PCA exhibited more precious division that mallard and spot-billed ducks were separated as expected, regarding species or place (Fig.1 a right; Fig. S13d). e.g.

Fig. S13d shows that PC1 separated domestic from the two wild and PC2 separated mallard from spot-billed. PC3 further divided the mallard and spot-billed ducks separately according to collection place. 2. Genetic structure analysis also shows distinct pattern of mallard and spot-billed (Fig.1d K=7). Although some mallard samples are closer to spot-billed when K=4, fewer spot-billed samples were similar with those mallard samples as K increase from 5 to 6. It could be interpreted as mallard and spot-billed have very similar genetic ancestry. This similarity is more likely due to their very recent divergence rather than hybridization. Because introgression would make individuals consist of different part of color like many previous researches on different cases (Fig. S15b)(M. Li et al., 2013). The recent divergence was further supported by the genomic features in subsequent speciation analysis and lack evidence of introgression between mallard and spot-billed, that were supported by distribution of ΔDAF across FST/DXY and ABBA-BBAA ratio bins. Moreover, Fig. S23-S24 shows the characters that discussed in genomic divergence of speciation section unlikely caused by random select two groups from mallard and spot-billed samples. Thus, we conclude mallard and spot-billed are monophyletic.

#### 4.6 ∂a∂i simulation

##### 4.6.1 Devising strategy to figure out the splitting order of domestic lineage and the two wild duck species.

There are two ways to compare the two phylogenetic trees (spot-billed, (mallard, domestic)) and (domestic, (mallard, spot-billed)). First, as some literates did, simulated the three populations split model and calculated the likelihoods to compare the fit of each model (Zhao et al., 2013).

We think this is unsuitable to our date. Because this would take too many parameters into model that would significantly increase the difficult to simulate and is very time consuming. Furthermore, the likelihoods are influenced by multiple parameters. We hardly give a property demographic model and constraint of those parameters, since there is lack of reliable evidence (beside genetic data) about the real evolutionary history of ducks. Moreover, both PCA and structure analysis shows that wild duck populations are structured that is challenging with ∂a∂i because the ∂a∂i model treats populations each as distinct homogenous entities. So, the best way is start with simplifying model and making it solid and clear to be get convergent parameters to make sure the first essential question. That is the splitting order. Because this could be determined by self-contained logic from multiple evidence (genetic structure, PSMC et al.) that alleviate the dependence on exactly divergence time and mutation rate.

We used an alternated way by simulating paired divergence and comparing the distribution of divergence time. If (Fig. 1e left) is the correct evolutionary relationship, then we expected longer divergence time between mallard duck and spot-billed than the ones between the two wild ducks and the domestic ducks. Otherwise if (Fig. 1e right) is the correct tree topology. We would expect the divergence time between two wild duck populations and domestic population were similar and longer than the one between mallard duck and spot-billed duck.

As we only focus on divergence time, we first used an extreme simple model with only two parameters (Fig. 2a left). This way would eliminate the influence between different parameters. Especially taking migration rate into simulation would have a significate impact in divergence time which could be purely due to mathematical property of ∂a∂i software.

The distribution of each estimated parameter and the confident intervals were presented in Fig. S16 and Table S12, respectively. And the comparison of divergence time between the domestic and mallard/spot-billed duck, mallard duck and spot-billed duck were shown in Fig. 2b left.

We then add migration parameters into model (Fig. 2a right). The distribution of each estimated parameter was presented in Fig. S17 that show bimodality distributions rather than normal, so we don't calculate the confident intervals. And the comparison of divergence time between the domestic and mallard/spot-billed duck, mallard duck and spot-billed duck under this model were shown in Fig. 2b right. The maximum likelihood Ts between domestic and mallard/spot-billed duck, mallard duck and spot-billed duck are 333,504 years ago, 334,738 years ago and 74,915 years ago respectively.

Combined the two models' result, we inferred that the ancestor of wild duck diverged from domestic duck about 100Kyr~300Kyr, and subsequently diverged into mallard duck and spot-billed duck about 70Kyr.

In conclude, dadi simulation on both models, considering or not considering the migration, are support the phylogenetic tree that is inferred from our previous population structure analysis. Although dadi software cannot distinguish short-divergence with low migration from long divergence with high-migration, this usually cause the estimate Ts not convergent or convergent into two peaks (smaller Ts accompanied by high migration or bigger Ts coupled with low migration rate). In our second simulations (with migration), estimate Ts does increased, when compared with first model, due to introduce migration. But the Ts still convergent into two peaks, higher estimated Ts with higher estimated migration rate whereas shorter Ts with lower migration rate. A reasonable explanation could be: Although dadi have difficult to determine longer or shorter Ts due to low/high migration rate estimation, thus having simulation on both situation and produce two peaks of estimate Ts, the Ts between mallard and spot-billed is still significant smaller than Ts between wild and domestic one in any case.

In additional, we add parameters to allow population change exponentially (IM_2 model). Clearly, the different between IM model and "split with migration" model is that: after divergence (during Ts), both populations could change population size exponentially. We found that many of simulations with IM model can't get the convergent estimates or get estimates with some parameters hit the boundary, especially for divergence between mallard duck and spot-billed duck. This may be because as the number of parameters (2 for "split_no_M" model, 4 for "split_M" and

7 for "IM_2" model) increased, the correlation between parameters (which is a purely mathematical property of) will significantly increase the difficulty to getting the correct estimates. And the two wild populations are highly structured and exist many fixed difference sites which is a particular challenge for dadi. Furthermore, the distribution of each parameters shown two or more peaks. So, we hardly estimated results. But we found that 29 of 39, 53 of 68 bootstrap fits yield increased domestic population size change after they split with mallard duck/spot-billed duck. All bootstrap fits yield increased wild duck population size change.
(data not show)
Statement on combining 8 domestic populations and two: 1) before the simulation presented in main text, we also separately used one of Beijing, Shaoxing and other 6 domestic populations (number of each sites' genotype of pool populations were inferred from frequency) to simulate divergence with mallard/spot-billed. These simulations got similar result as well as the estimated divergence time. It's also no big difference with the simulation that combine domestic.2) PCA and tree analysis supported.

**4.6.2 Data sets and infrastructure**

For the ∂a∂i simulation analysis, each runs based on different bootstraps data sets. The likelihoods distribution of each runs are presented in Fig. S18. The SNPs were filtered according to the coverage of resequencing data at their position with thresholds of 20-fold for pooled data and a total number of genotyped alleles of more than 50, 36, 26, and 28 for Beijing, Shaoxing, mallard duck and spot-billed duck populations, respectively. The SNPs (single-nucleotide polymorphisms) were sampled randomly per 100 kb from the SNPs identified in wild populations (mallard and spot-billed) and domestic populations (Cherry Valley, Campbell and 6 Chinese duck breeds).

Three models' code list:

```
def split_nm(params,ns,pts):
    s,Ts=params
    xx=dadi.Numerics.default_grid(pts)
    phi=dadi.PhiManip.phi_1D(xx)
    phi=dadi.PhiManip.phi_1D_to_2D(xx,phi)
    phi=dadi.Integration.two_pops(phi,xx,Ts,nu1=s,nu2=(1-s))
    fs=dadi.Spectrum.from_phi(phi,ns,(xx,xx))
    return fs
```

model code1: split without migration

```
def split_m(params,ns,pts):
    s,Ts,m12,m21=params
    xx=dadi.Numerics.default_grid(pts)
    phi=dadi.PhiManip.phi_1D(xx)
    phi=dadi.PhiManip.phi_1D_to_2D(xx,phi)
    phi=dadi.Integration.two_pops(phi,xx,Ts,nu1=s,nu2=(1-s),m12=m12,m21=m21)
```

```
    fs=dadi.Spectrum.from_phi(phi,ns,(xx,xx))
    return fs
```

model code2: split with migration

```
def IM_2(params,ns,pts):
    s,nu1,nu2,TS,m12,m21=params
    xx=dadi.Numerics.default_grid(pts)
    phi=dadi.PhiManip.phi_1D(xx)
    phi=dadi.PhiManip.phi_1D_to_2D(xx,phi)
    nu1_func=lambda t: s *(nu1/s)**(t/TS)
    nu2_func=lambda t: (1-s) *(nu2/(1-s))**(t/TS)
   phi=dadi.Integration.two_pops(phi,xx,TS,nu1=nu1_func,nu2=nu2_func,m12=m1
2,m21=m21)
    fs=dadi.Spectrum.from_phi(phi,ns,(xx,xx))
    return fs
```

model code2: split with migration and population exponentially change.

full software infrastructure include SNP datasets preparation and simulations is provided in https://github.com/vitogump/life.

**4.6.3 Comparisons with previous studies**

Some may think L should be the total length of sequence (variant and invariant) that was used for variant calling. According to the author of dadi and the principle of this software, if researcher didn't use all variants from the L region, they should scale the length by the rate of the SNP selected or the simulation would conduct mislead results which some research may make. As a long divergence time between domestic and their wild relative looks like a highly unlike result (but it is possible), this may lead many research take some trick to avoid this result. Our principle for simulation is that: selected some parsimonious models that are most informative to the issue we concerned (1. what's the order of divergence of mallard, spot-billed and domestic lineage, 2. Then try to point a rough time intervals of the two divergence time) as discussed above; used a range of initial values for each parameter which more than biologically reasonable ranges, let them naturally converge through simulation process and try neither let a setting boundary restrict the estimates, nor cutting a hard bound by removing some estimates. i.e. After model determination, let software determining the parameters itself and keep every simulation results. In the first look we used the same methodology (dadi simulaitons) on similar data conducted very different findings. However, some research used rather complex models to try to include the information about bottle neck after domestication and the detail divergence between specific breeds in domestic, and judge which model is the best simultaneously, by only judging likelihood value of this kind value. In fact, some domestic research (some used dadi) have carried out similar result (divergence time between domestic and wild) with us, however, this kind results were discarded artificially by claiming "very deep divergence (~150 kyrs) is not possible for domesticated species" in some of them (Freedman et al., 2014; Wang et al., 2013). Our research provocative accepting a deep divergence time is encouraged by the special situation of ducks that the two closest wild relatives of domestic ducks may have a very short divergence time, even shorter than the domestic lineage splitting from them. This was illuminated by previous analysis. The feature of genome divergence between mallard and spot-billed are also in accordance with a very recent divergence time.

##### 4.7 Selection analyses and genomic islands

In the $d_f$ calculation, we required all sampled individuals were sufficient covered for each df sites.

Whole genome alignment to chicken were performed by LASTZ(Harris, 2007).

###### 4.7.1 Detail functional analysis on candidate selected genes in early ancestor of mallard and spot-billed ducks.

We detected positive selections in the early ancestors of these two wild populations using methods invented in early modern humans and extended in European and Asian wild boards pig (Green et al., 2010; Groenen et al., 2012). SNPs shared between wild and domestic lineages are mainly ancient variations presented before their splitting. Derived frequency in domestic can be a function of frequency in wild. Ancient sweeps in wild will fixed selected and linked variations, whereas new variations will reshape the regions' nucleated diversity. Those SNP will not be shared with domestic and mostly not skewed towards lower frequency spectrum. This signal mainly utilized middle to high frequency spectrum of wild alleles that would produce high prediction of domestic frequency, especially for those regions undergone ancient selection that wild derived allele absented in domestic lineage. Joint distribution of nucleotide diversity and S value further confirm those candidate regions detected by S were not enriched to low nucleotide diversity (prone to be caused by recent selection)(Fig. S20g). As described in previous research, local recombination don't interferes the power of this methods. We use a relative narrow window width to detect ancient selection because duck genomes' have a higher recombination rates and we prone a high sensitive. We filtered variation that passed the following criteria:

1) No CpG sites
2) More than 30-fold coverage when SNP fixed in Muscovy or both outgroups
   And both outgroups were fixed the same directions.
3) At least 5 individuals were observed in each population or 10 reads were observed for pool populations
4) At least 20 reads were observed in bam when the site was treated as fixed to ref allele in a population
5) Derived allele was detected in wild lineage

For windows with less than 10 variation sites were discarded.

In addition to the functional analysis presented in main text. An overrepresented term of positive selection related to "dorsal/ventral pattern formation", "determination of left/right symmetry" also imply selection on genes which making their body more suitable for flying. overrepresented on terms "response to wounding", "egg activation", "acute inflammatory response to antigenic stimulus", "detection of virus" may indicate that selection on genes increased mallard and spot-billed ducks' survivability in wild. We note that P-values associated with each GO term are not robust to multiple testing. This may due to the small size of each GO term which leading to relatively large number of GO terms (8750 in total, 1050 GO terms contain at least 1 candidate selected gene). Furthermore, ancient selection may associate with long-distance flight ability evolution may involve widespread functional changes. Although our detected selected region are enrich to ancient sweeps, long-term evolution after that may also obscure our signals. However, our main intent was not to identify specific great significance terms but to reveal a trend that top serval terms associated with functions involved in long-range migration ability.

In addition, we found that at least 32 genes (e.g. *IL11RA*, *PRKAA1*, *LPAR1*, and *NS1BP*) involved into immune response to infections and three of them (i.e, *ADAP*, *RIG-I* and *CX3CR1*) played critical roles against influenza A virus infections (Table S15)(Barber, Aldridge, Webster, & Magor, 2010; C. Li et al., 2015). Therefore, it seemed that positive selections on immune genes might help the early mallard and spot-billed duck ancestor keep evolutionary equilibrium with influenza A viruses.

Some cryptochrome-containing cells within the retina are active at night when the birds perform magnetic orientation (L. Q. Wu & Dickman, 2012). In some migratory birds, a distinct part of the forebrain, where primarily processes input from the eye, is highly active at night(Liedvogel et al., 2007). We also found three genes associated with retinal ganglion cell axon guidance (*EFNA5*, *ISL1* and *EPHA7* (S=-3)) and two genes (*CHRD*, *DIXDC1* and *WNT7B*) related to forebrain regionalization or progenitor cell division were under selection in the early mallard duck and spot-billed duck ancestor (Fig. 3d; Fig. S21)(Anderson, Lawrence, Stottmann, Bachiller, & Klingensmith, 2002; F. Wu et al., 2015). Gene family that function on short wavelength-sensitive opsin (Cryptochromes, which have been suggested to form the basis of light-dependent magnetic compass in birds, is excited by blue (short wavelength) light) expanded in wild ducks may also indicated an enhancement in (Table S9)(Liedvogel et al., 2007).

**4.7.2 Detail analysis on genomic islands of mallard and spot-billed duck divergence.**

To elucidate genomic properties and speciation of these two wild duck species, we firstly calculated how many sites all mallard ducks homozygous for one allele and all spot-billed ducks homozygous for another (which we referred the density of these fixed difference sites as $d_f$) in 40kb sliding windows with 20kb step size along duck scaffolds (Methods). This effort found that a small number of dfs (4,386) and only 26 SNPs of these located within protein-coding regions distributed in small genomic regions (758 of totally 52,215 windows) between mallard and spot-billed. We refer the windows showing highly elevated divergence up to 50 times higher than the genomic mean as “divergent peak”. Divergent peaks contain most (3,522, 80%) of these dfs that distribute in 348 (0.6% of the whole duck genome) windows. This is sharp contrast to the case in Ficedula flycatchers, where using the same criterion, that 25% (59,936) dfs were distributed in the large genomic divergent islands covering 2.7% of the flycatcher genome. Such large difference in number of dfs and the pattern of genomic divergent islands in two these species might due to that the wild duck (the mallard and spot-billed duck) has a shorter divergence time and is in an earlier stage than the Ficedula flycatcher.

As our sequence depth (~10X) is higher than the flycatchers' research (~5X), which may leading to the reduce of the dfs. We also random sample the reads into about 5-fold coverage of duck genome and the distribution of dfs remain almost the same (data not shown).

In order to study the genomic divergence in duck autosomes, we then estimated the population differentiation, nucleotide diversity through calculating $F_{ST}$ (fixation index), dxy (sequence divergence), π (nucleotide diversity) and Tajima's D values in sliding window that homologous to chicken autosomes. (Methods; Fig. 4a; Table S16). Scaffolds homologous to chicken Z chromosome showed higher level of $F_{ST}$ (5.5-fold) and Tajima's D (1.5-fold), but smaller number of SNP density (0.5-fold) and lower level of π (0.6-fold) than autosomes (Table S16, Fig. S23). These observations are consistent with change of $F_{ST}$, Tajima's D and SNP density in flycatcher and wild ducks (Mexican and mallard duck)(Ellegren et al., 2012; Lavretsky et al., 2015). Analysis on autosomal differentiation regions shows that the distribution of $F_{ST}$ between the mallard duck and spot-billed ranged from -0.0157 to 0.7785 with an average of 0.04930346 and distributed in a narrow peak, lower than $F_{ST}$ between the wild ducks and domestic ducks ranged from 0.0880 to 0.8912 with an average of 0.2100 and distributed in a wide peak (Fig. 4b). Detailed analysis indicates that small part of genomic regions of the mallard duck and spot-billed duck have $F_{ST}$ values being away from the mean $F_{ST}$ at least 4 standard deviations (we referred these regions as high differentiation islands). These higher $F_{ST}$ regions showed lower level of nucleotide diversity (π mallard: 0.0024 ± 0.0022; π spot-billed: 0.0023 ± 0.0020) and SNP density (360 ± 305 and 334 ± 280 SNP per window in the mallard duck and spot-billed duck respectively) than genomic backgrounds (π mallard: 0.0076 ± 0.0042; π spot-billed: 0.0077 ± 0.0043. 1134 ± 569 and 1161 ± 597 SNP per window in the mallard duck and spot-billed duck respectively) in both mallard duck and spot-billed duck (Fig. 4c-f; Fig. S23; Table S17). This is similar to the case in flycatchers, Darwin's finches and ducks, where high differentiation islands are characterized by reduced levels of nucleotide diversity and polymorphisms (Ellegren et al., 2012; Groenen et al., 2012; Han et al., 2017; Lavretsky et al., 2015). A noteworthy consequence of these coinciding features was that both Tajima's D of these high differentiation islands was not reduced in both mallard duck and spot-billed, whereas it is different with the case in flycatchers that Tajima's D of high differentiation islands was significantly reduced (Fig. 4f; Table S17). Moreover, sequence divergence between mallard duck and spot-billed didn't exceed background levels in high differentiation islands that is consistent with the observation in flycatchers, whereas Darwin's finches' study show elevated dxy in high differentiation islands (Discussion Section; Fig. S23)(Han et al., 2017).

##### 4.7.3 Detail analysis on identification of candidate domestication regions

Limited by many factors, we used very conservative strategy, may not the very advance in methods, that has been well discussed and succeed applied in previous reports, to detect genomic regions under domestic selection. We first used the

Z-transformed distributions of Hp in domestic and wild ducks, we find no apparent functional overlap (data not shown). We chose to set the thresholds at zHp<=−4 and zHp<=−2 for autosomes and sex-chromosomes separately. We applied this thresholds because Z chromosome including a reduction in effective populations size and recombination rate as previous discussed (Axelsson et al., 2013). Although the skewed distribution of heterozygosity scores may indicate a confounding between signals of selection and genetic draft (Fig. S26, S27). We used combining sequence data from all domestic pools to alleviate this problem and extreme tails of the distributions should be enriched for true signals of selection. We then combine zHp with Z-transformed $F_{ST}$ signals. We perform the statistic test on this outlier approaches using 4 times of $zF_{ST}$ and zHp to define selective region, 4 times represent the extreme ends of the distribution of z $F_{ST}$ and zHp, and P-value is less than 0.05, $zF_{ST}$ ($P(zF_{ST} >= 4) = 0.00548$), zHp($P(zHp <= -4) = 0.00322$). Two signals combined identified 137 regions with extremely low levels of Hp (average length = 63.6230 kb, average Hp = 0.2100 for autosomes; Hp = 0.2470 for sex-chromosomes) and 145 regions with significantly elevated $F_{ST}$ values (average length = 58.7670 kb, average $F_{ST}$ = 0.5390 for autosomes, average $F_{ST}$ = 0.394 for sex-chromosomes). We focused on putatively selective sweep regions falling at least four/two standard deviations away from the mean of the Z(Hp) and Z($F_{ST}$) of autosomes and sex-chromosomes separately. However, only a small proportion putative selective sweeps regions were predicted using both extremely low levels of Hp and significantly elevated $F_{ST}$ values (Fig. S28). Such observation might be partly attributed to that the domestic lineage and wild lineage (mallard duck and spot-billed ancestors) were under different evolutionary patterns after they had diverged about 100-200 Kyr. For example, the wild lineage seemed to benefit from positive selection and increased their effective population sizes once or twice times, while the domestic lineage dramatically declined their effective populations sizes during LGP (Fig. 2d,3a; Fig. S19). This complex demographic history of the domestic and wild lineage might affect the $F_{ST}$ distribution.

When we counting the heterozygosity of these populations (Hp) in 40-kb windows, those windows containing less than 10 SNPs were not included in eight duck breeds respectively. We further calculated the Hp mean of these breeds for each window. Similarly, we calculated fixation index ($F_{ST}$) between each domestic duck breeds and the wild duck population (combine spot-billed and mallard) using an estimator introduced in dog in 40-kb windows, and counted the $F_{ST}$ of eight domestic duck breeds for each window(Karlsson et al., 2007). After that, we Z-transformed the Hp and $F_{ST}$ to Z(Hp) and Z($F_{ST}$), and defined regions with Z(Hp) or Z($F_{ST}$) being four standards deviations for autosomes or two standard deviations for sex-chromosomes from the mean of the genome-wide (Z(Hp) as putative selective sweep regions.

In Hp calculation, we didn’t require min SNP numbers because the region with extreme little SNP could be Runs of homozygosity (ROH) that is also our interest.

For the regions under early stage selection in wild lineage or under recent selection in domestic lineage. We performed more detail analysis to visualize the genetic variations in Fig. S21, S29.

The comparison between the details of selection regions in Fig. S20 reflect that the early stage selections mostly couldn't be visualized by current variations directly. But regions under recent selection in domestic lineage showed clear trace (Fig. S29).
GO enrichment analysis were performed by using Fisher's Exact Test (see https://github.com/vitogump/life).

###### 4.7.4 Evolutionary relationship of domestic duck, mallard wild duck and spot-billed wild duck

Such inference was supported by both phylogenetic comparative analysis and population genetic analysis. Despite the incomplete lineage sorting (ILS) and gene flow may confound the analyses, the different SNP sets used in genetic structure analysis, f3-statistics, SNPs distributions analysis and tree construction that was sampled from genome-wide would avoid these confounding effects in phylogenic inference(Harr, 2006). Furthermore, Genetic structure show each individual of both the mallard duck and spot-billed duck almost consistent of purely one ancestry component. This is different from the case of admixture, where individuals of one population with components of other populations' ancestry (de Manuel et al., 2016; M. Li et al., 2013; Prado-Martinez et al., 2013), thus indicating no evidence of introgression between the mallard and spot-billed duck (Fig. 1d; Fig. S15). Subsequent ∂a∂i and PSMC simulations that rely on different data sets and theory reconstructed the perfect matched demographic history further support the robustness of our inference. In addition, the subsequent observations of $F_{ST}$ on domestication and speciation analyses are consistent with or could be best explained by this phylogenic inference (Fig. 4b).

##### 4.8 Estimation of genetic diversity and distance

Nucleotide diversity (π) was calculated using the vcftools(Danecek et al., 2011). Tajima's D was calculated using the Popgenome R package(Pfeifer, Wittelsburger, Ramos-Onsins, & Lercher, 2014) Pairwise $F_{ST}$ was calculated in 40-kb windows with a sliding size of 20-kb. Neighbor-joining tree based on pairwise $F_{ST}$ values were constructed with phylip-3.696 "neighbor" function by taking the distance matrix of pairwise $F_{ST}$ and integrating neighbor-joining trees of 27,374 windows.

##### 4.9 select candidate casual mutation to verify its functional activity

For the sake of simplicity, we catalog the selection regions by its overlap with CDS, UTR, intron, intergenetic region. And rank SNPs by the ΔAF between wild lineage and domestic lineage. we selected the SNPs that located in UTR, intron, intergenetic region and closed to the genes that may play important functional in the duck evolution or domestication.
Supplementary Figures

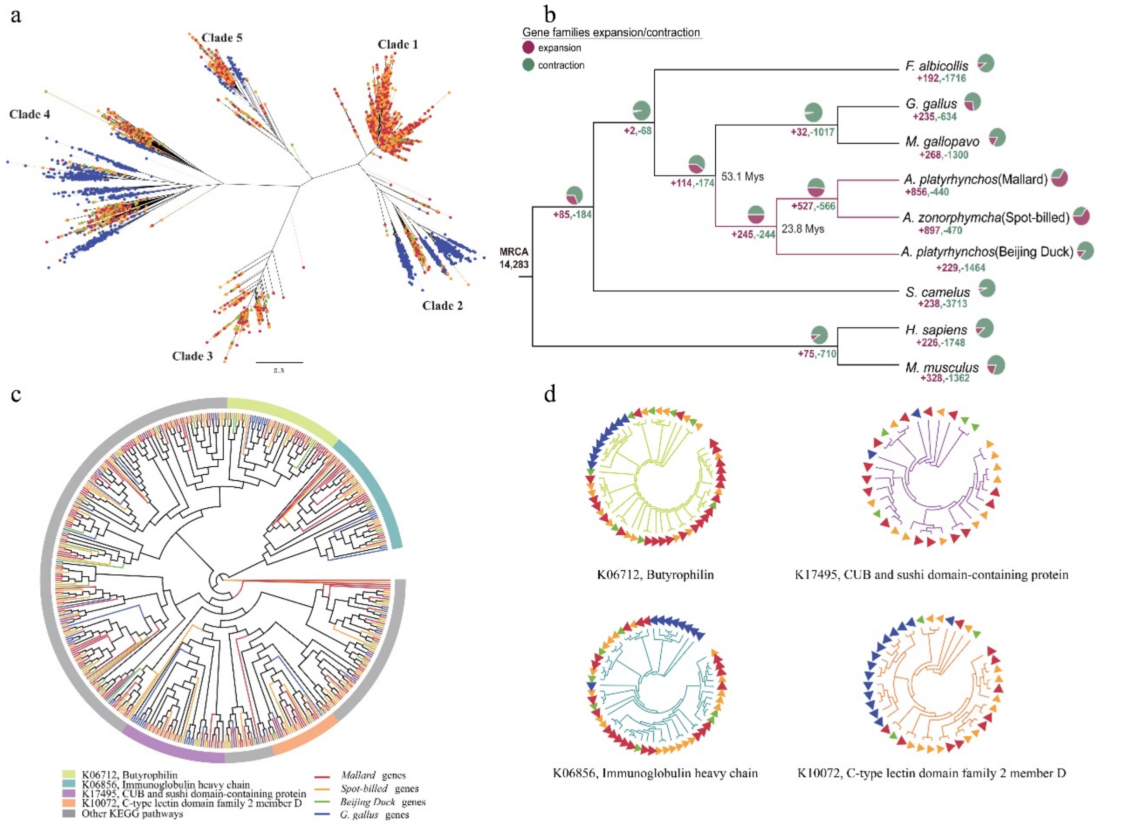

**Fig. S1: Expansion and contraction of transportable element and gene in mallard, spot-billed and Beijing duck genome.**

Green, red, yellow and blue in a and c panel represent Beijing duck, mallard, spot-billed and chicken, respectively. (**a**) Phylogeny of CR1 retrotransposons in four bird genomes. (**b**) Numbers of gene family losses and gains across ten vertebrates. Pie chart and numbers nearby indicate the number of expanded and contracted gene families in each branch. (**c**) Phylogeny of gene families expanded in spot-billed and mallard. Top four families ranked by the number of gene gained are indicated in color arcs. KEGG pathway annotation of those four families are listed in the bottom. Other families are indicated in grey color. (**d**) Detailed phylogeny of four gene families expanded in spot-billed and mallard duck as indicated in (**c**).

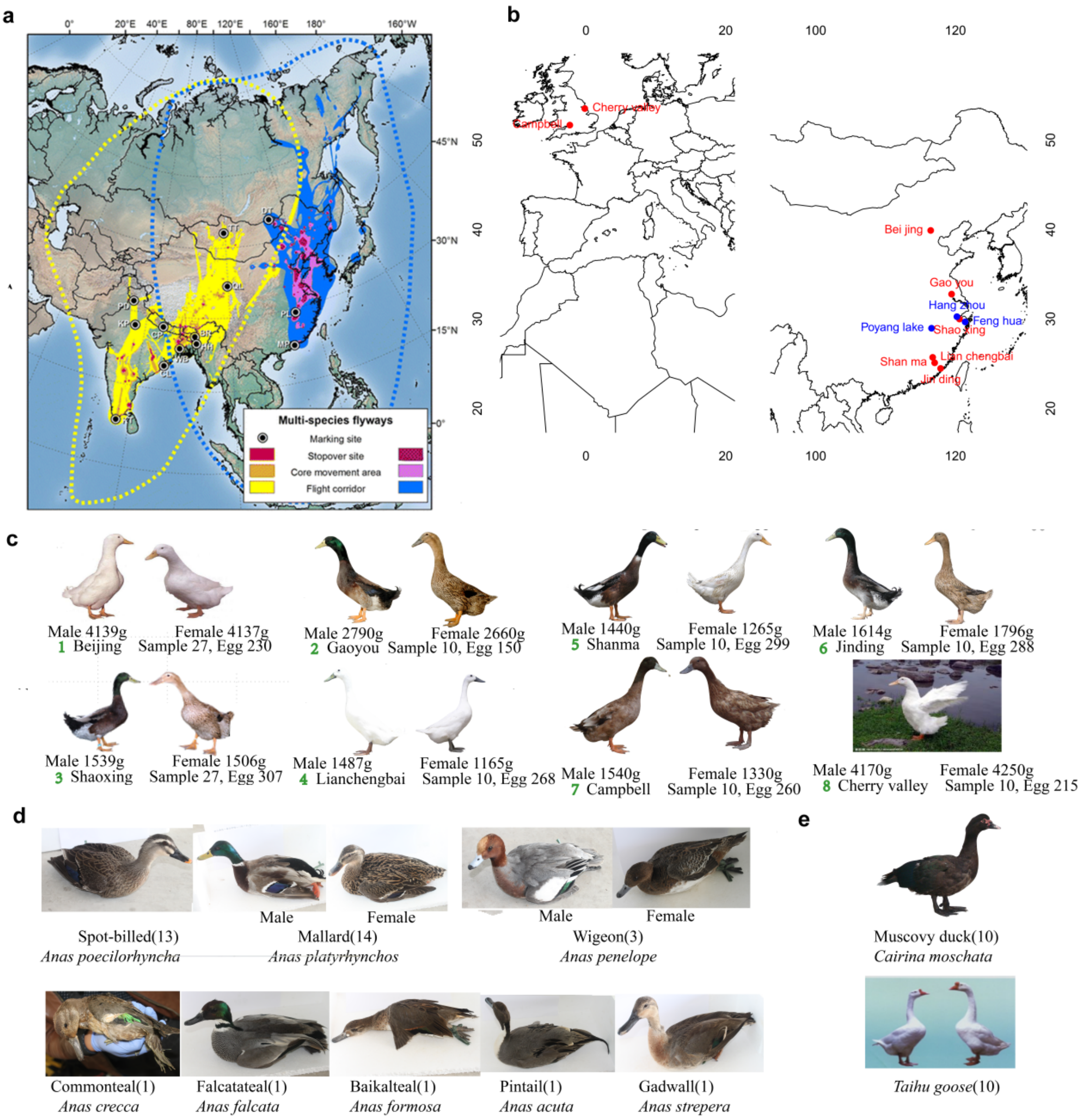

**Fig. S2. Sample distribution and phenotypic variation in domestic and wild ducks.**

Photos were taken from the sequenced samples except Taihu goose and Cherry valley being download from web. (**a**) Estimated migration routes of Anatidae in Central Asian Flyway (CAF) and East Asian-Australasian Flyway (EAAF)**(Palm et al., 2015)**. Regions in the flyway of CAF displayed in yellow-red and for EAAF displayed in blue-purple. From darkest to lightest, colors represent 50%, 75% and 99% cumulative probability contours. (**b**) Geographical distribution of wild duck and domestic duck samples. Blue dots represent the place where wild duck samples collected from. Compare to **a**, our wild duck samples were collection from the main migration route of the Anatidae flyway. (**c**) Phenotype of the eight domestic duck breeds. Number of individuals, average weight (male/female), average eggs production per year were marked. (**d**) Photos and sample numbers of the eight wild duck species. (**e**) Photos and sample number of Muscovy duck and Taihu goose used as outgroup.

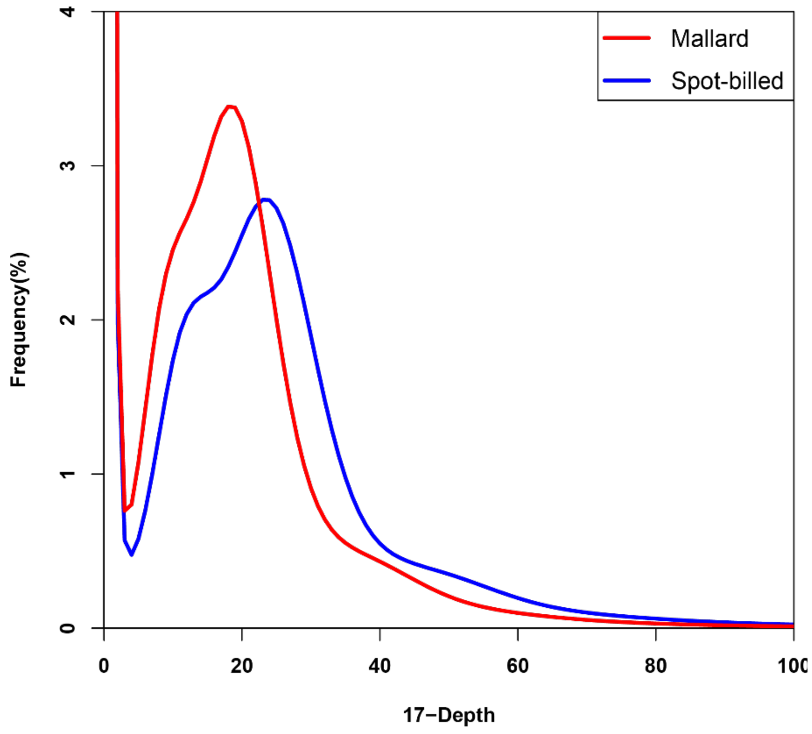

**Fig. S3. Distribution of 17-mer frequency in the corrected pair-end reads of the mallard and the spot-billed genome.**

Only the reads from short-size libraries (< 500 bp) were included in this analysis. We identified 22,312,838,068 kmers using 21-fold data of mallard and 28,137,616,208 kmers using 27-fold data of spot-billed. The D2B and D2L genome size can be estimated as G= K_number (total kmer number) / K_depth (the volume peak), which is estimated to be 1.14Gb and 1.15Gb, respectively.

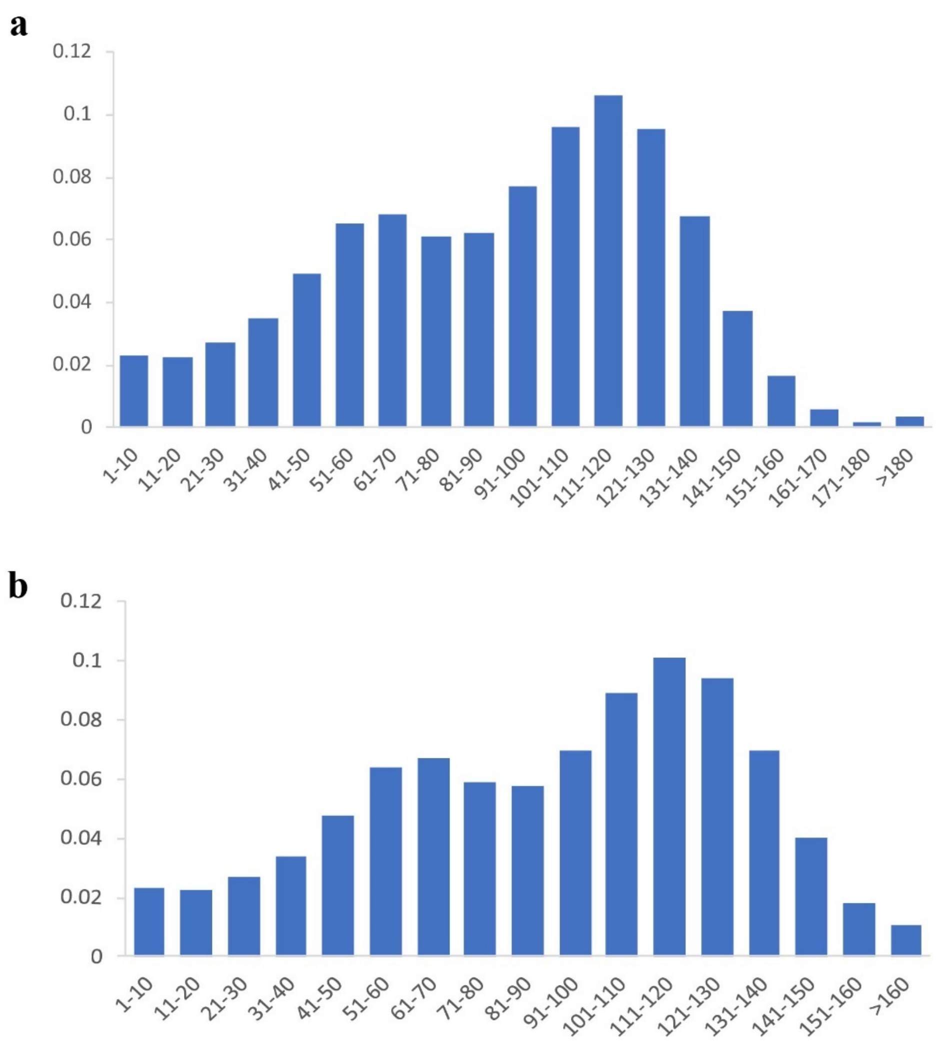

**Fig. S4. Distribution of the sequencing depth of the mallard and the spot-billed assemblies.**

(a) the mallard assembly. (b) the spot-billed assembly.

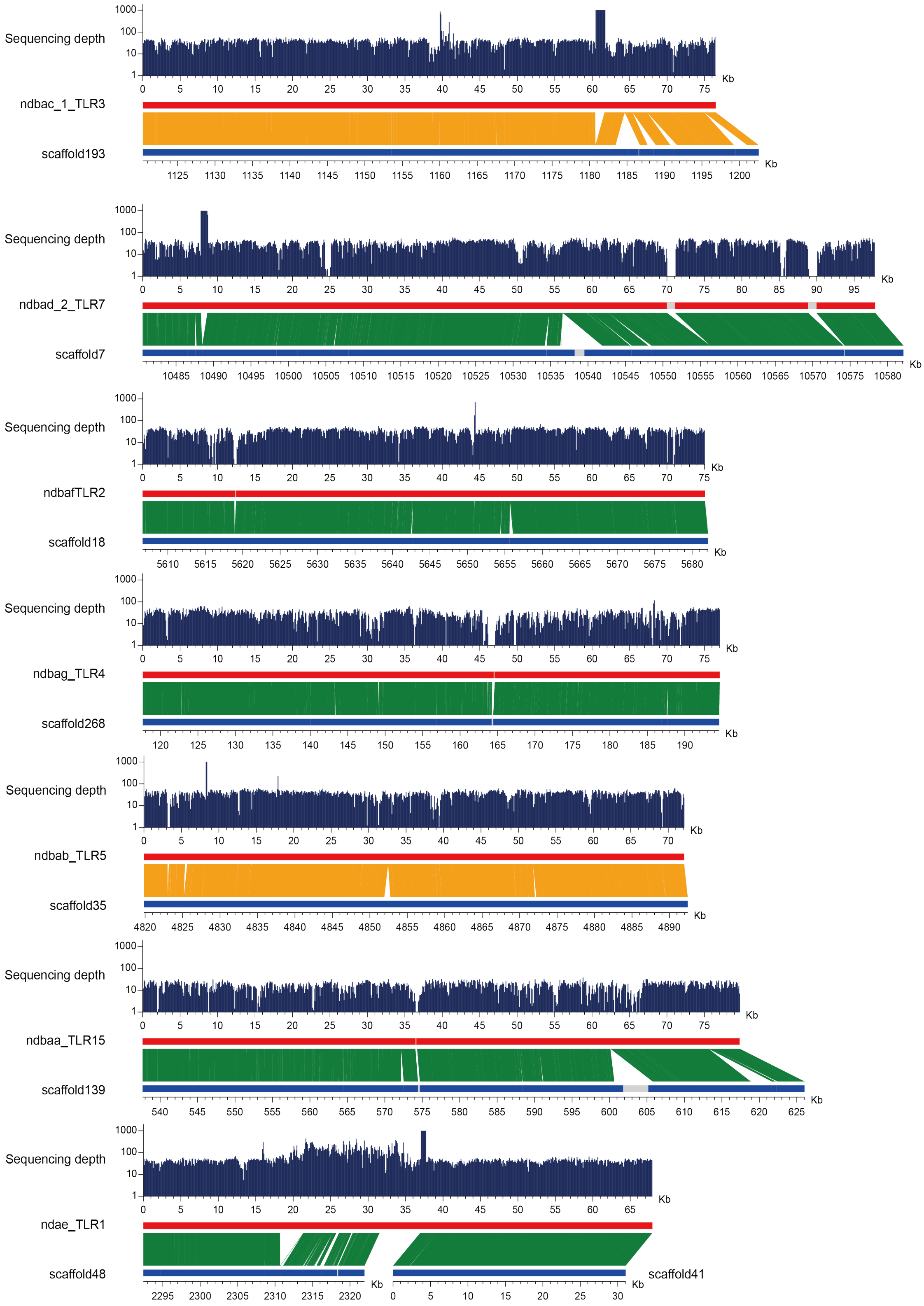

**Fig. S5. Evaluation of the mallard assembly with sequences of seven BACs.** Seven BACs were from chromosome 1, 3 or 4. The BAC sequences and assembly scaffolds are shown in red and dark blue, respectively. Orange represents positively and green was reversely aligned, the remaining unclosed gaps on the scaffolds are marked as white blocks.

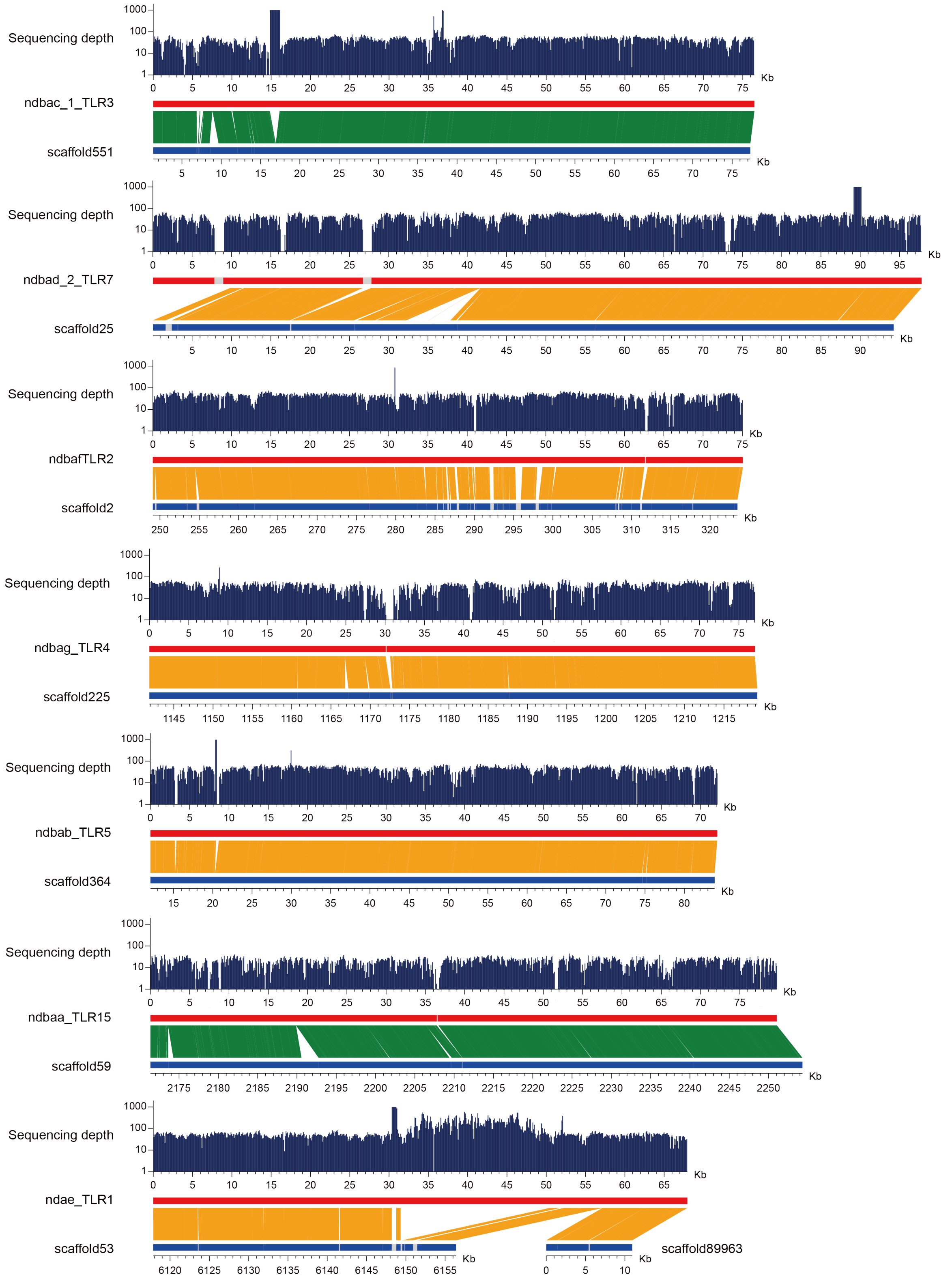

**Fig. S6. Evaluation of the spot-billed assembly with sequences of seven BACs.** Seven BACs were from chromosome 1, 3 or 4. The BAC sequences and assembly scaffolds are shown in red and dark blue, respectively. Orange represents positively and green was reversely aligned, the remaining unclosed gaps on the scaffolds are marked as white blocks.

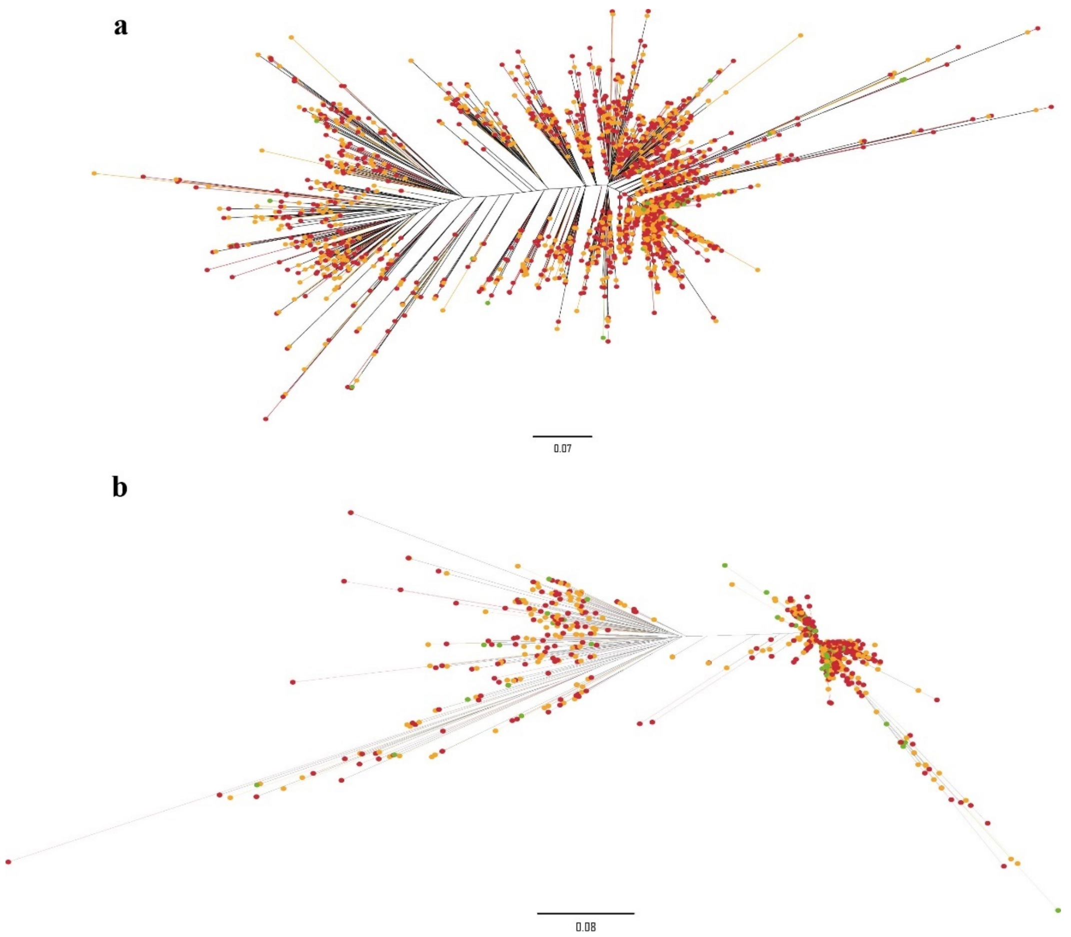

**Fig. S7. Phylogeny of two CR1 retrotransposon clade in four bird genomes.**
Green, red and yellow represent Beijing duck, mallard and spot-billed, respectively.

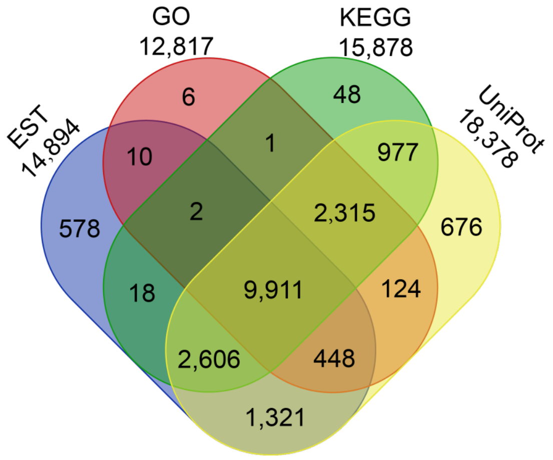

**Fig. S8. Venn diagram showing the mallard reference genes annotated using three databases or supported by EST.**
Total 21,056 mallard genes supported by 319,996 ESTs of Beijing duck and functioned annotation with three databases. And 19,041 genes were supported by one of them.

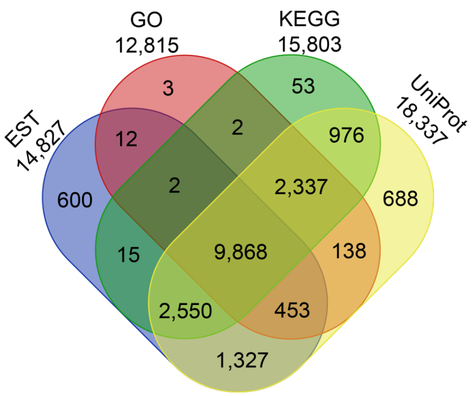

**Fig. S9. Venn diagram showing the spot-billed reference genes annotated using three databases or supported by EST.**

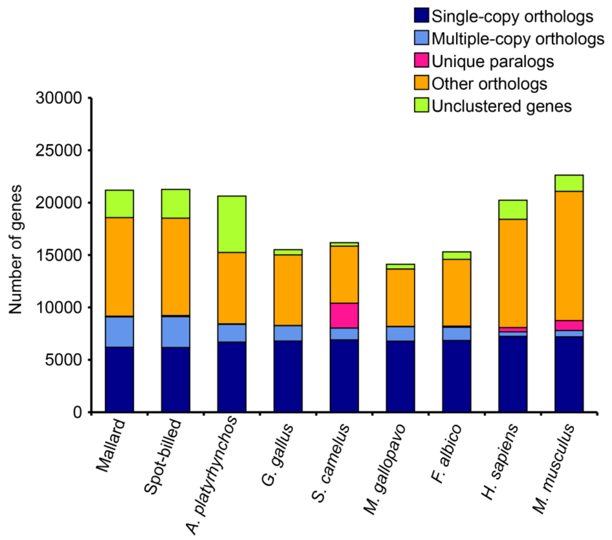

**Fig. S10. Protein homologous comparison among the reference gene sets of nine species by TreeFam.**

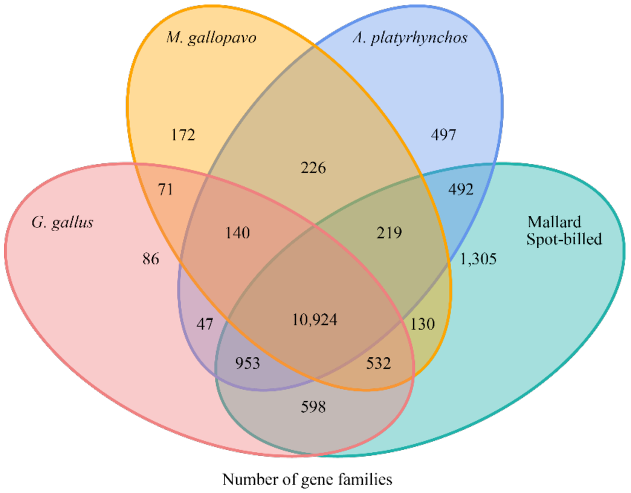

**Fig. S11. Venn diagram shows gene family clusters in five species by TreeFam.**

The number of unique and shared gene families among five genomes.

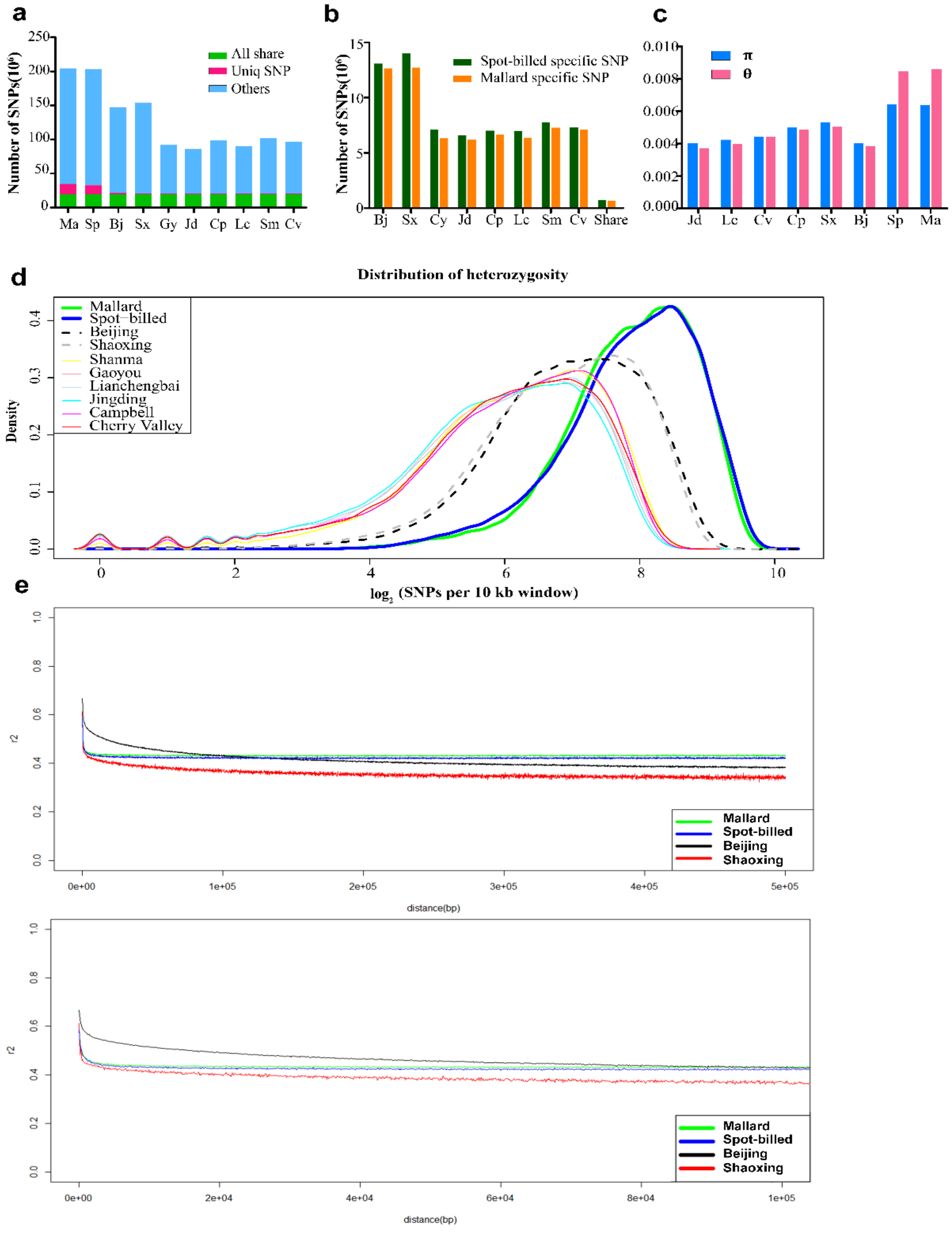
a
Number of SNPs(10⁵)
All share
Uniq SNP
Others
Ma Sp Bj Sx Gy Jd Cp Lc Sm Cv
b
Number of SNPs(10⁶)
Spot-billed specific SNP
Mallard specific SNP
Bj Sx Cy Jd Cp Lc Sm Cv Share
c
π
θ
Jd Lc Cv Cp Sx Bj Sp Ma
d
Distribution of heterozygosity
Density
Mallard
Spot-billed
Beijing
Shaoxing
Shanma
Gaoyou
Lianchengbai
Jingding
Campbell
Cherry Valley
log2 (SNPs per 10 kb window)
e
r2
Mallard
Spot-billed
Beijing
Shaoxing
distance(bp)

f

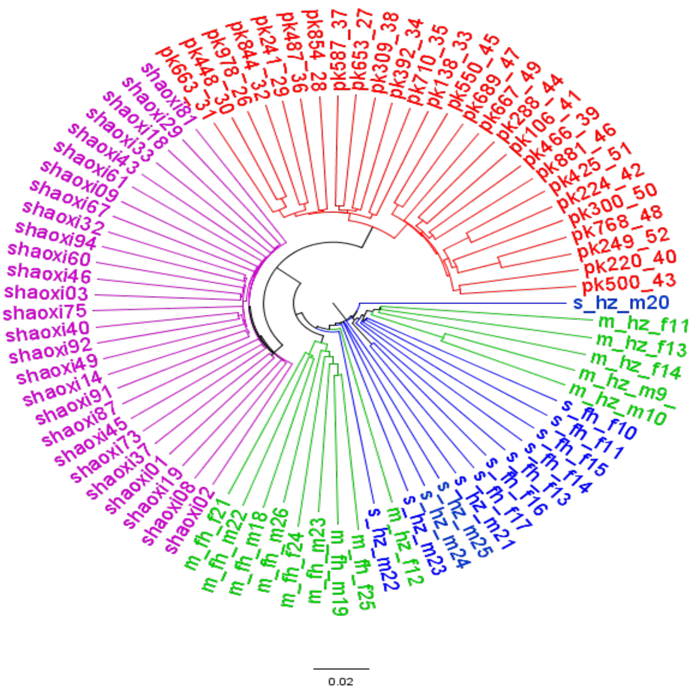
pk663_31
pk448_30
pk978_26
pk844_32
pk241_29
pk487_36
pk854_28
pk587_37
pk653_27
pk309_38
pk392_34
pk710_35
pk138_33
pk550_45
pk689_47
pk667_49
pk288_44
pk106_41
pk466_39
pk881_46
pk425_51
pk224_42
pk300_50
pk768_48
pk249_52
pk220_40
pk500_43
s_hz_m20
m_hz_f11
m_hz_f13
m_hz_f14
m_hz_m9
m_hz_m10
s_fh_f10
s_fh_f11
s_fh_f15
s_fh_f14
s_fh_f13
s_fh_f16
s_fh_f17
s_hz_m21
s_hz_m25
s_hz_m24
s_hz_m23
m_hz_f12
s_hz_m22
m_fh_f25
m_fh_m19
m_fh_m23
m_fh_f24
m_fh_m26
m_fh_m18
m_fh_m22
m_fh_f21
shaoxi02
shaoxi08
shaoxi19
shaoxi01
shaoxi37
shaoxi73
shaoxi45
shaoxi87
shaoxi91
shaoxi14
shaoxi49
shaoxi92
shaoxi40
shaoxi75
shaoxi03
shaoxi46
shaoxi60
shaoxi94
shaoxi32
shaoxi67
shaoxi09
shaoxi61
shaoxi43
shaoxi33
shaoxi18
shaoxi29
shaoxi81
0.02

g

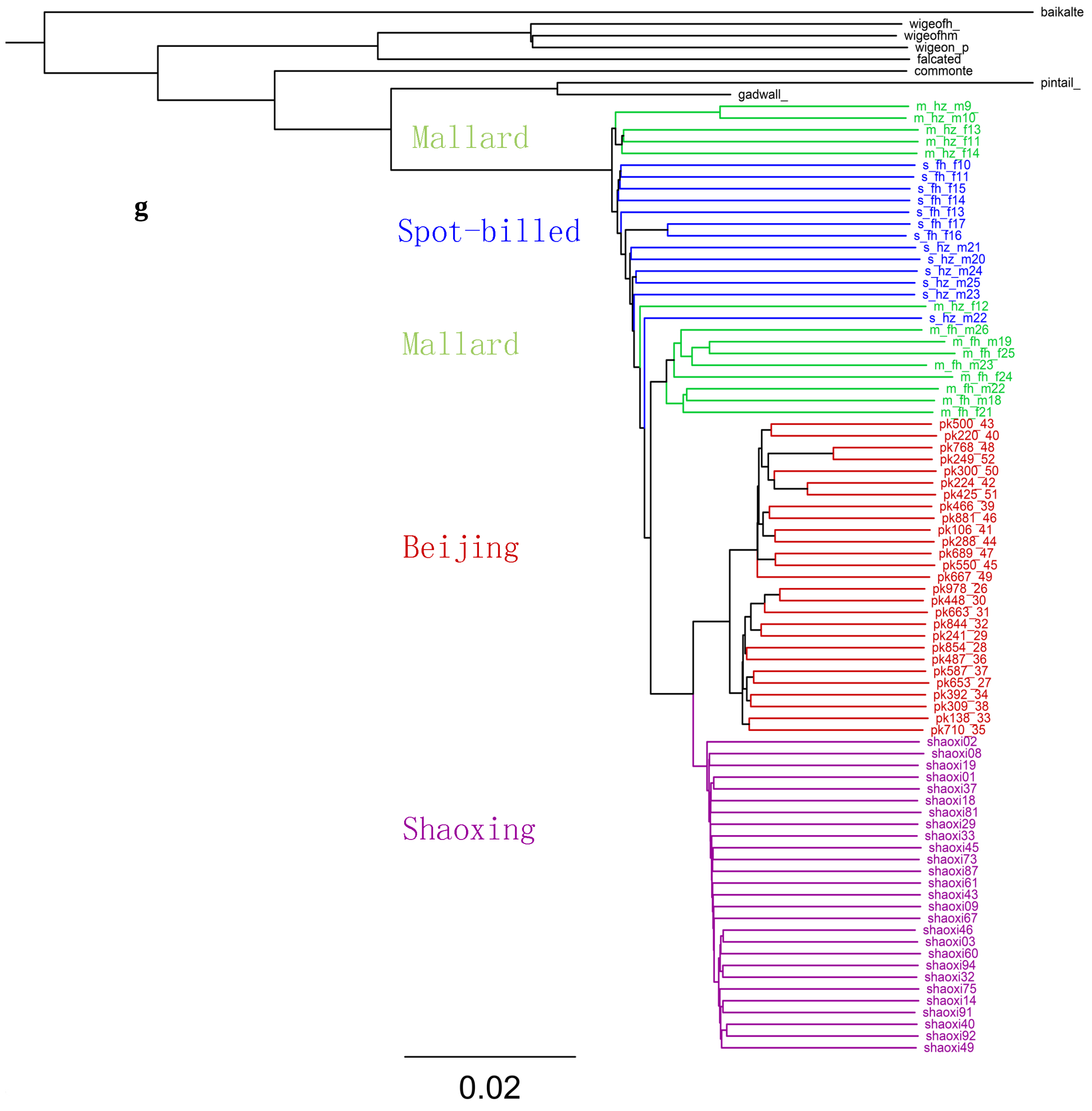
baikalte
wigeofh_
wigeofhm
wigeon_p
falcated
commonte
pintail_
gadwall_
Mallard
Spot-billed
Mallard
Beijing
Shaoxing
m_hz_m9
m_hz_m10
m_hz_f13
m_hz_f11
m_hz_f14
s_fh_f10
s_fh_f11
s_fh_f15
s_fh_f14
s_fh_f13
s_fh_f17
s_fh_f16
s_hz_m21
s_hz_m20
s_hz_m24
s_hz_m25
s_hz_m23
m_hz_f12
s_hz_m22
m_fh_m26
m_fh_m19
m_fh_f25
m_fh_m23
m_fh_f24
m_fh_m22
m_fh_m18
m_fh_f21
pk500_43
pk220_40
pk768_48
pk249_52
pk300_50
pk224_42
pk425_51
pk466_39
pk881_46
pk106_41
pk288_44
pk689_47
pk550_45
pk667_49
pk978_26
pk448_30
pk663_31
pk844_32
pk241_29
pk854_28
pk487_36
pk587_37
pk653_27
pk392_34
pk309_38
pk138_33
pk710_35
shaoxi02
shaoxi08
shaoxi19
shaoxi01
shaoxi37
shaoxi18
shaoxi81
shaoxi29
shaoxi33
shaoxi45
shaoxi73
shaoxi87
shaoxi61
shaoxi43
shaoxi09
shaoxi67
shaoxi46
shaoxi03
shaoxi60
shaoxi94
shaoxi32
shaoxi75
shaoxi14
shaoxi91
shaoxi40
shaoxi92
shaoxi49
0.02

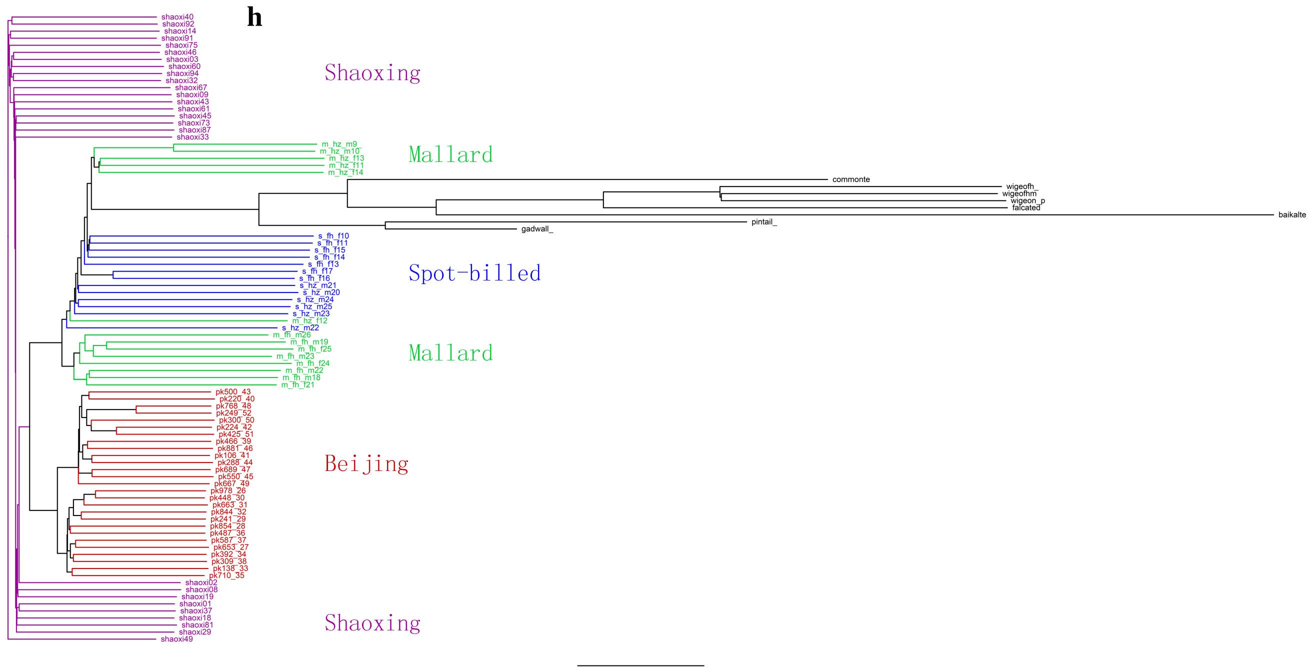

**Fig. S12. Distribution of genetic variation and characteristics for two wild and eight domestic duck populations.**

Mallard, Spot-billed, Beijing, Shaoxing, Gaoyou, Jinding, Campbell, Lianchengbai, Shanma and Cherry valley are represented by Ma, Sp, Bj, Sx, Gy, Jd, Cp, Lc, Sm, Cv. (**a**) Total number of SNPs. (**b**) Total number of mallard and spot-billed specific SNPs segregating in each domestic population. (**c**) Genetic diversity was measured by nucleotide diversity π and Watterson's estimator ϴ. (**d**) Distribution of heterozygosity. (**e**) LD decay measured by $r^2$ estimator against distance (50 kb up and 10 kb down) between the SNPs in mallard, spot-billed, Beijing and Shaoxing populations. Different way to display IBS tree: **(f)** root on the midpoint of mallard (green), spot-billed (blue), Beijing (red), Shaoxing (purple) **(g)** root on the midpoint of mallard, spot-billed, Beijing, Shaoxing and other 6 wild species (black) **(h)** root on an arbitrary point of mallard, spot-billed, Beijing, Shaoxing and other 6 wild species (black).

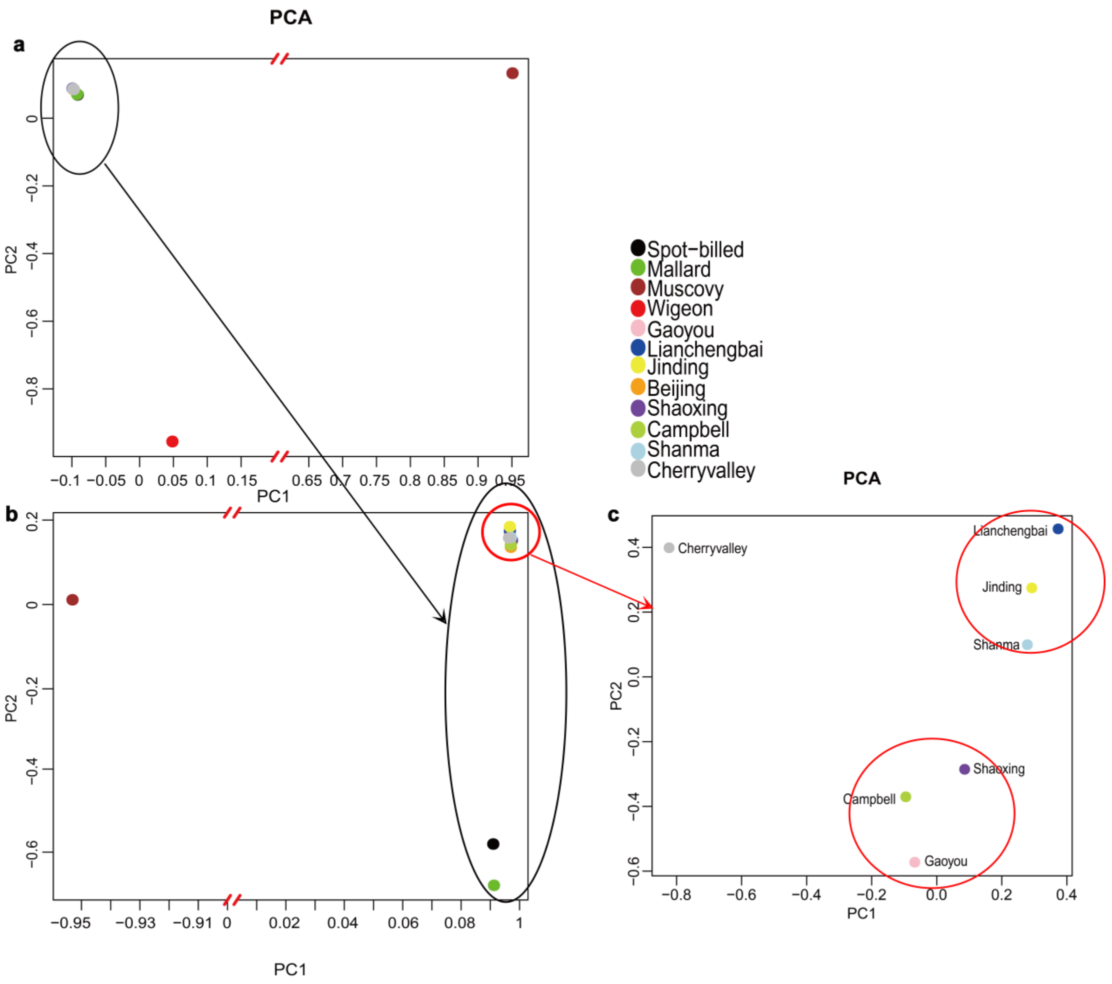
PCA
a
b
c
PC1
PC2
Spot−billed
Mallard
Muscovy
Wigeon
Gaoyou
Lianchengbai
Jinding
Beijing
Shaoxing
Campbell
Shanma
Cherryvalley
Cherryvalley
Lianchengbai
Jinding
Shanma
Shaoxing
Campbell
Gaoyou

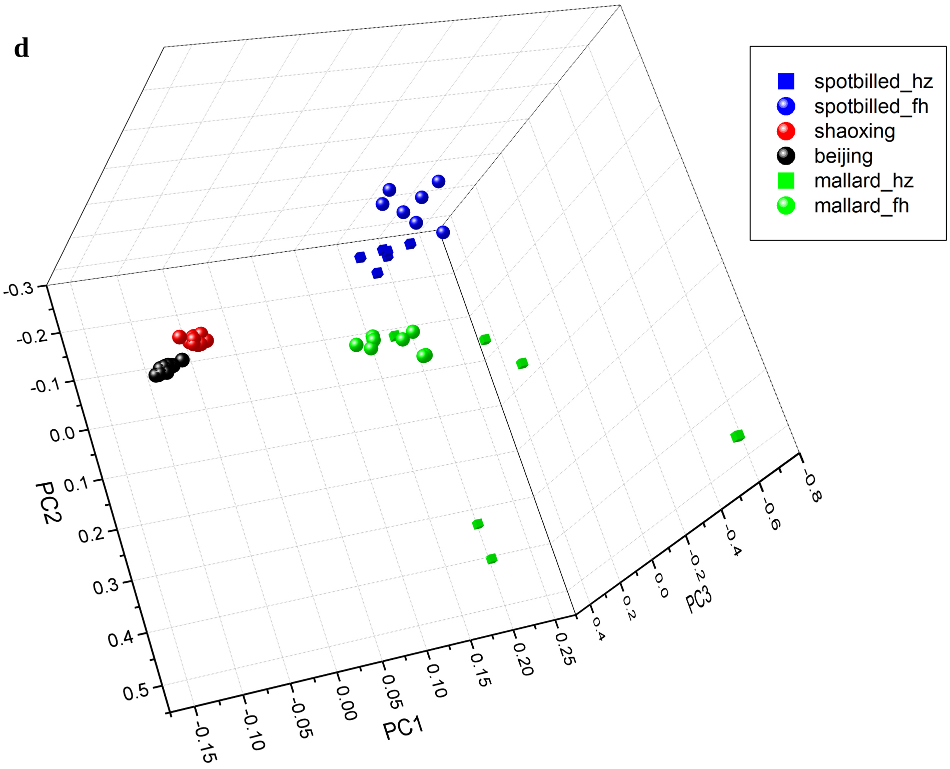
d
spotbilled_hz
spotbilled_fh
shaoxing
beijing
mallard_hz
mallard_fh
PC1
PC2
PC3

**Fig. S13. Principal component analysis of wild and domestic ducks using population as unit.**

(**a**) PCA analysis with genome wide SNPs set from 8 domestic duck breeds and three wild duck species, that having population data, together with Muscovy ducks. Two wild duck (mallard and spot-billed) and eight domestic duck populations were clustered in the top left, while the wigeon populations (consist of 3 individuals) and the muscovy duck population showing large distance with eight domestic duck populations. (**b**) PCA analysis on eight domestic duck populations and two wild duck (mallard and spot-billed) populations along with one muscovy duck population. It shows that 8 domestic duck populations cluster together and clearly separated from Mallard and Spot-billed in the second component. (**c**). PCA analysis on seven domestic duck populations. It shows that these populations clustered according to their geographical distributions. (d). 3D PCA on four populations. Random sample Shaoxing and Beijing to equal sample size of each as the two wild populations.

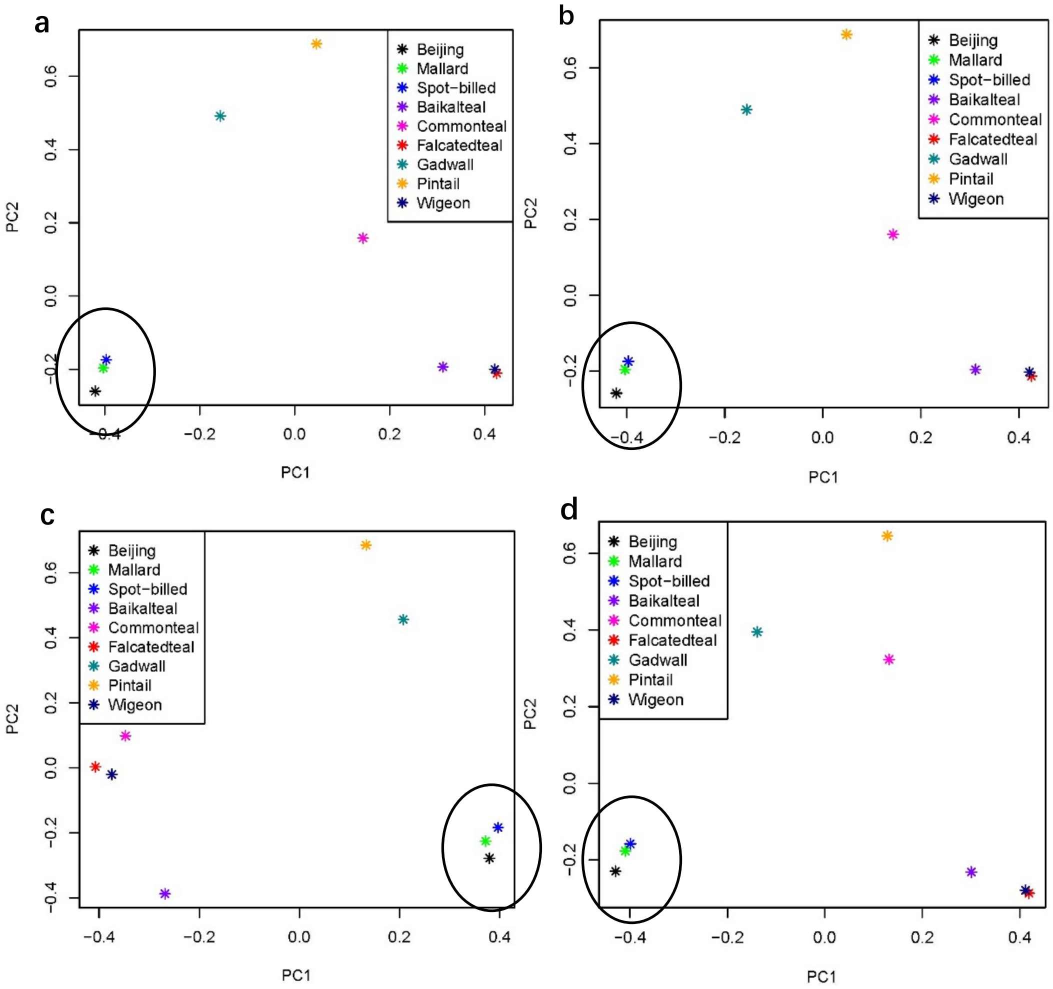

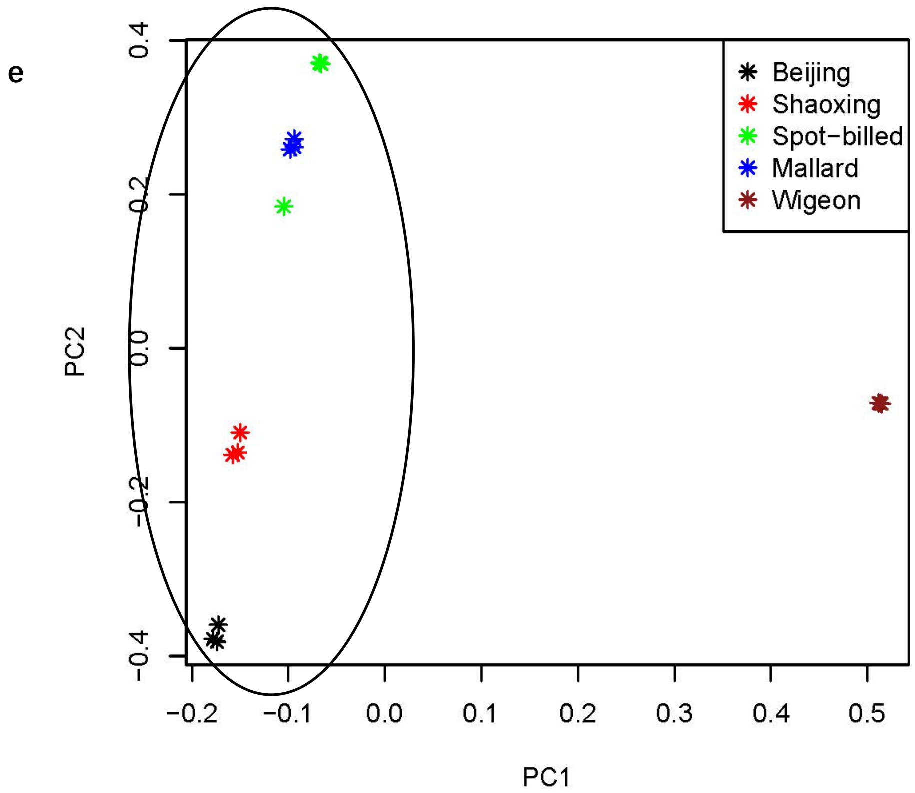

**Fig. S14. Principal component analysis of wild and domestic duck with same coverage and sample size for each population.**

(a-d) shows PCA analysis on eight wild species (one individuals each, random sample reads into 12× that is equal to Beijing individual) and a Beijing individual with different data process strategy. a) using LD pruned SNPs from all scaffolds of Beijing assemble. b) using LD pruned SNPs from the scaffolds regions which were homologous to chicken autosomes. c) using LD pruned SNPs from the scaffolds regions which were homologous to chicken Z chromosome. d) using LD pruned SNPs from all scaffolds of mallard duck genome assemble. e) 3 individuals for each of the 5 populations (mallard duck, Spot-billed, Beijig, Shaoxing and wigeon, random sample reads into 3× for each individuals).

Black cycles indicate spot-billed, mallard duck and domestic duck are clustered together.

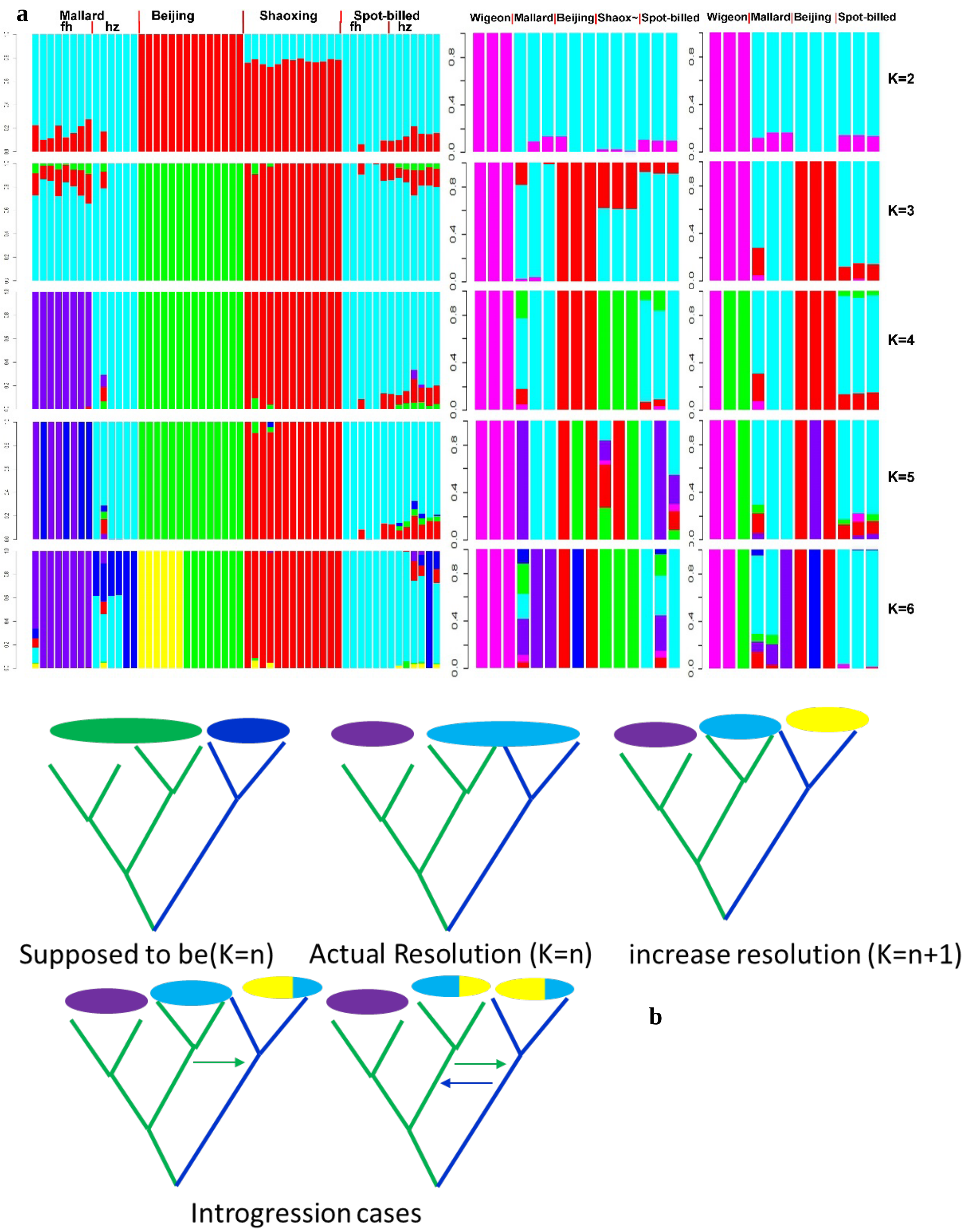

**Fig. S15. Genetic structure analysis with equal sample size and same coverage of each population for different populations combinations.**

a.) Samples in left panel were random sampled to 3-fold coverage. Beijing duck samples were from a designed population, which were from a same population and formed a high growth line and low growth line after several generations selection. Samples in middle panel were random sampled to 3-fold coverage and three samples for each of the 5 populations. Samples in the right panel were random sampled to 12-fold coverage and three samples for each of the 4 populations. K range from 2 to 6.

b.) clade (e.g. mallard colored by green, spot-billed colored by blue) identified by different ancestry (color of oval). Models to understand Fig. 2 d

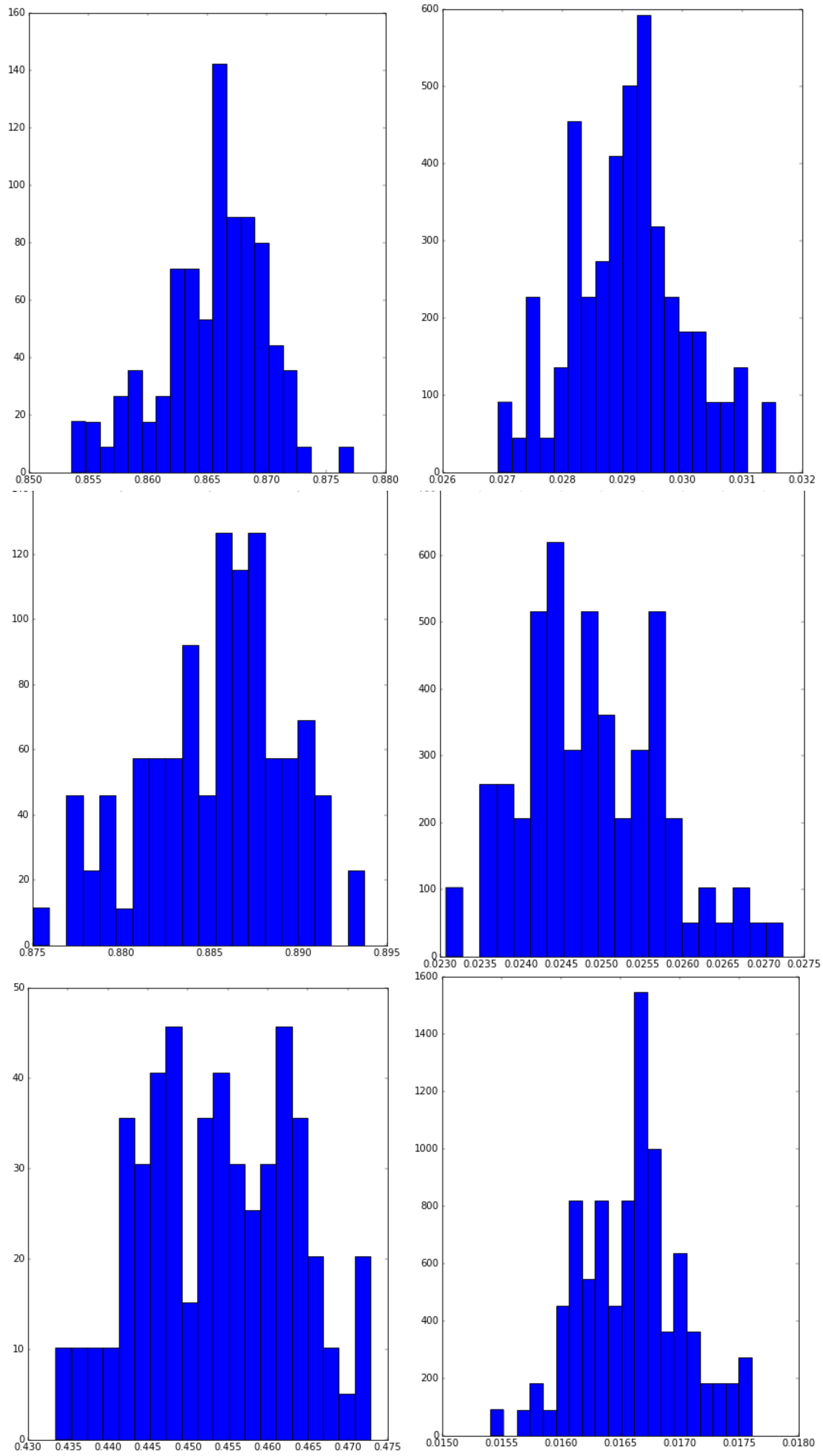

**Fig. S16. Split without migration model conventional bootstrap results.**

Panels from up to down corresponding to simulations of mallard and domestic duck divergence, spot-billed and domestic duck divergence, mallard duck and spot-billed duck divergence, separately. Left picture of each panels shows the distribution of s, whereas right one shows the distribution of Ts.

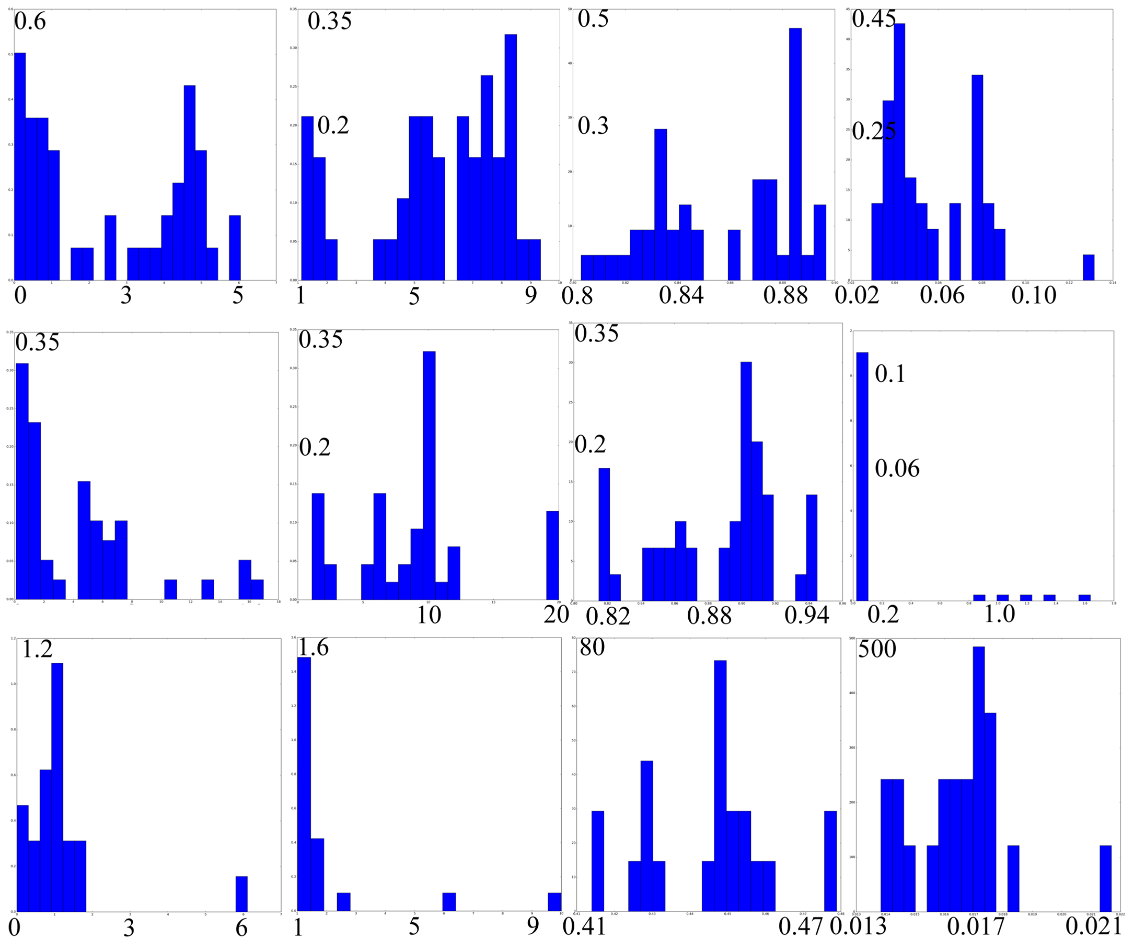

**Fig. S17. Split with migration model conventional bootstrap results.**

Panels from up to down corresponding to simulations of mallard and domestic duck divergence, spot-billed and domestic duck divergence, mallard duck and spot-billed duck divergence, separately. Columns from left to right corresponding the distributions of m12, m21, s, Ts parameters.

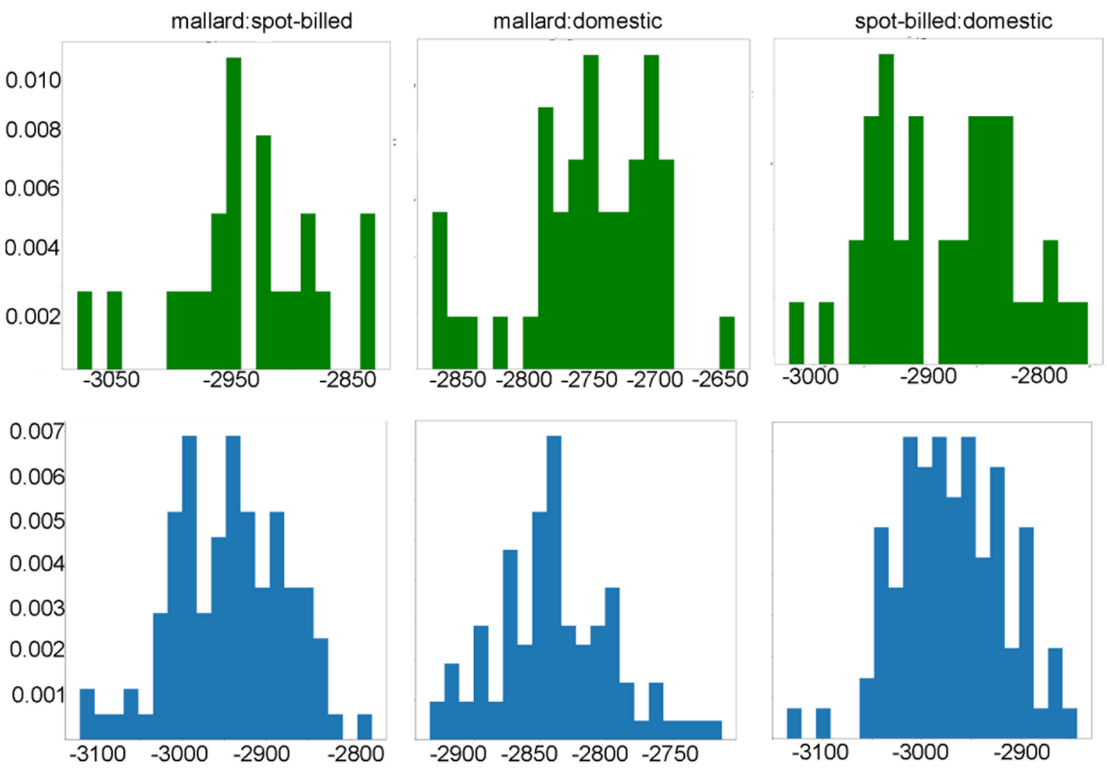

**Fig. S18. Likelihood Distribution from bootstrap results.** Distribution of likelihood produced by using model with migration/without migration are shown as blue picture/green picture, respectively. Simulation of divergence between mallard and spot-billed (M and SP), mallard and domestic (M and D), spot-billed and domestic (SP and D) are performed by both models each.

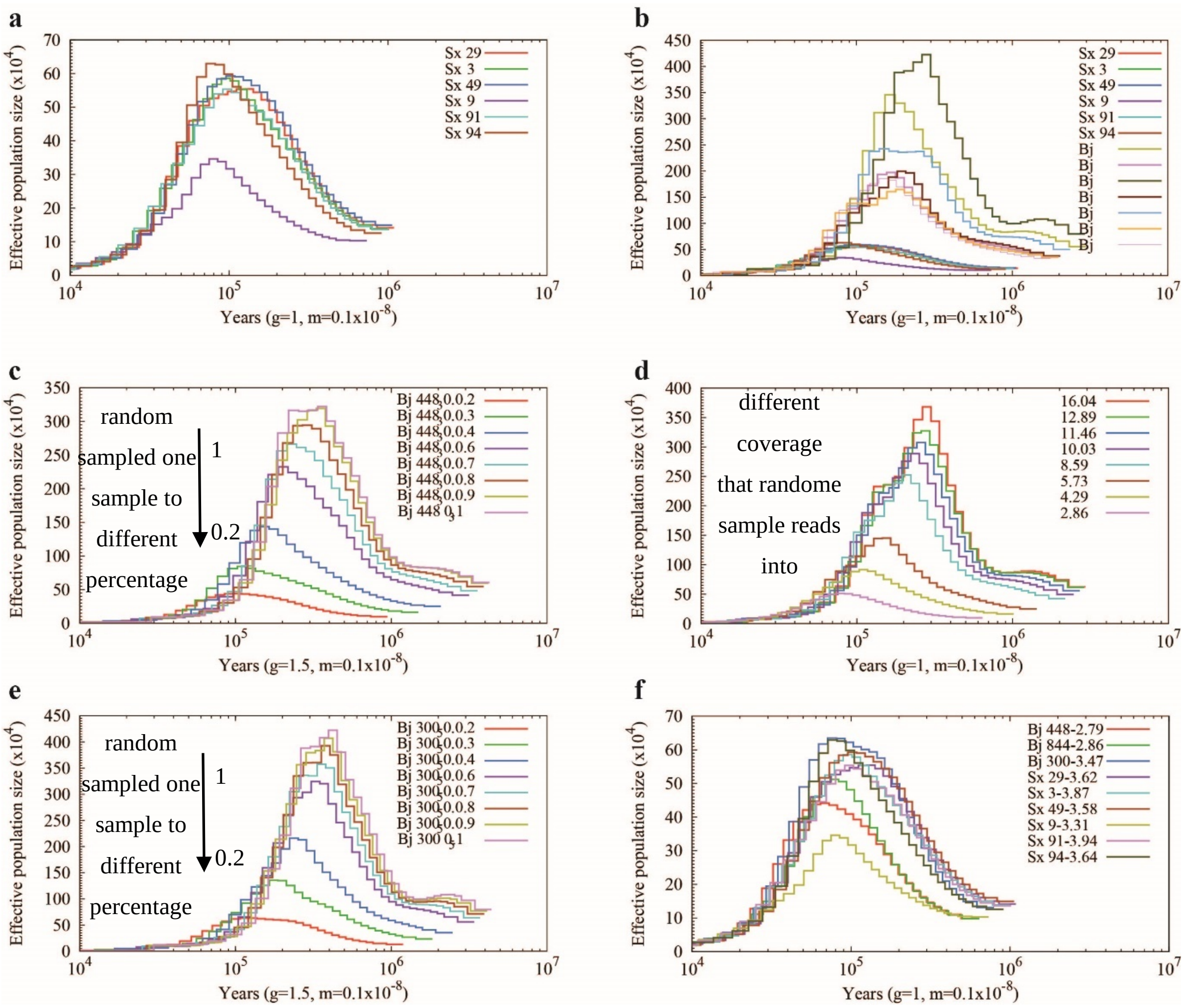

**Fig. S19. Comparison of PSMC between Beijing (Bj) and Shaoxing (Sx) ducks with different coverage data.**

(**a**) Demographic history of six Shaoxing ducks with a coverage of 2- to 3.5-fold. (**b**) Comparison demographic history of six Shaoxing (2- to 3.5-fold) and Beijing (13.5-fold) ducks. (**c-e**) PSMC analyses of Beijing based on different coverage of random sample reads. Sample ID in **c**, **d** and **e** are 44448_30, 43844_32, 41300_50, respectively. Coverage are represented in different color lines in legend. (**f**) PSMC curve of Beijing and Shaoxing ducks inferred with comparable coverage reads are similar.

**a**

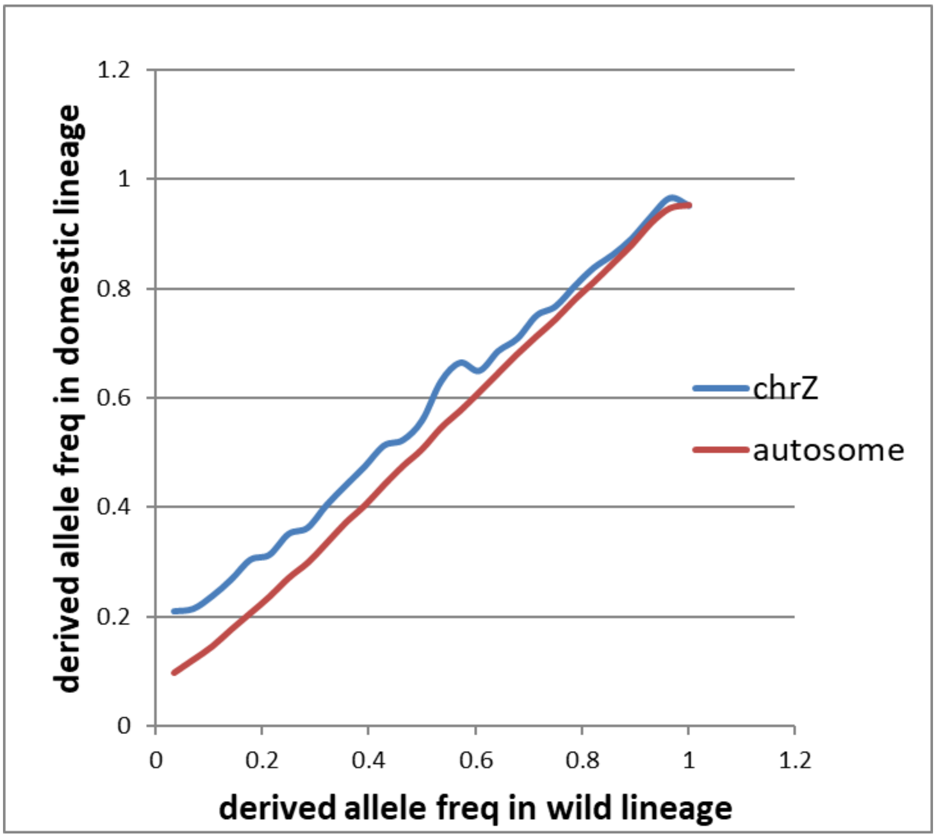
derived allele freq in domestic lineage
derived allele freq in wild lineage
chrZ
autosome
0
0.2
0.4
0.6
0.8
1
1.2

**b**

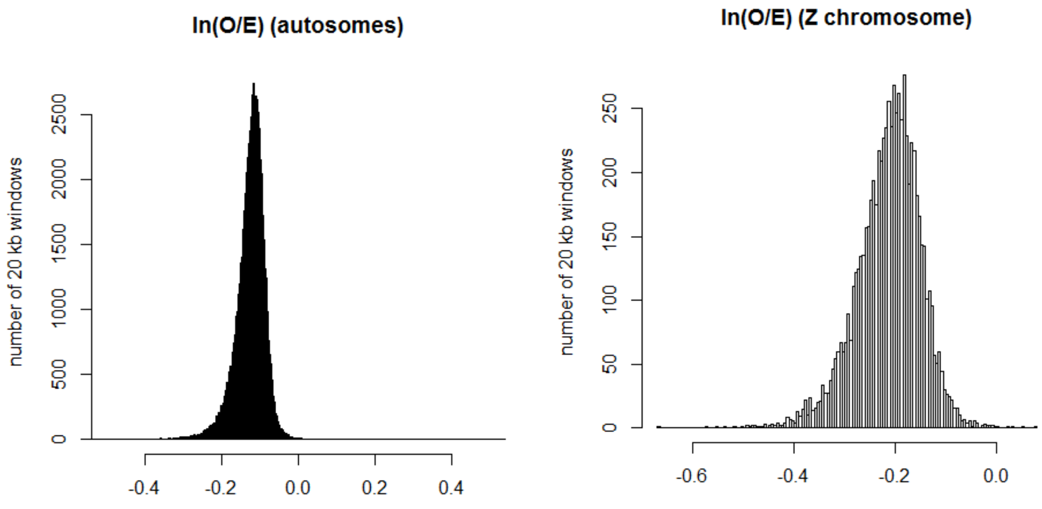
ln(O/E) (autosomes)
ln(O/E) (Z chromosome)
number of 20 kb windows
number of 20 kb windows
0
500
1000
1500
2000
2500
-0.4
-0.2
0.0
0.2
0.4
0
50
100
150
200
250
-0.6
-0.4
-0.2
0.0

c

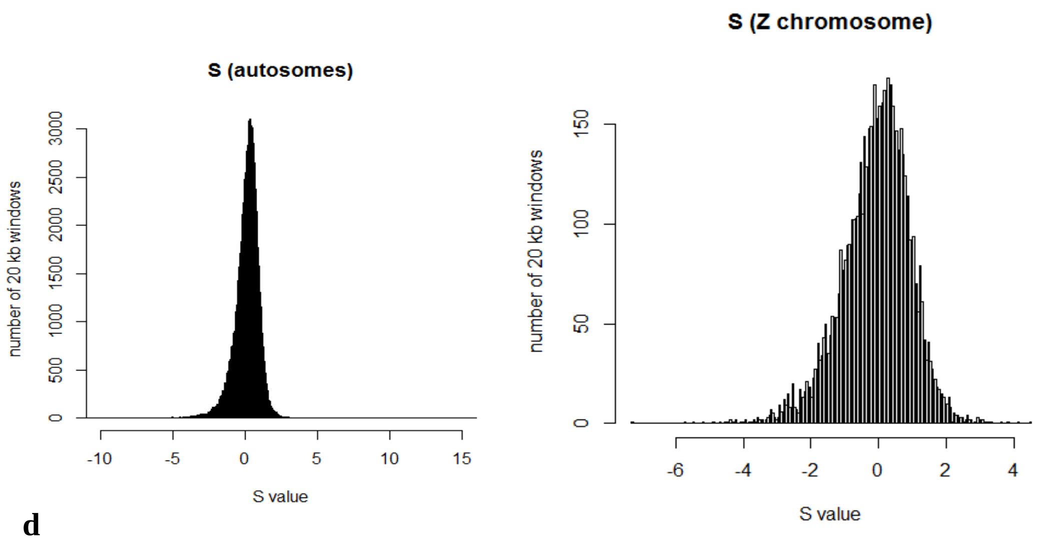
S (autosomes)
number of 20 kb windows
S value
S (Z chromosome)
number of 20 kb windows
S value

d

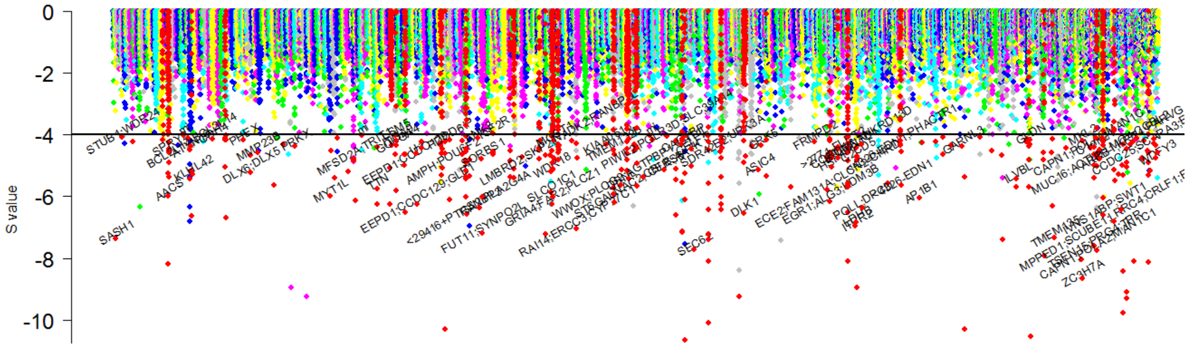
S value

e

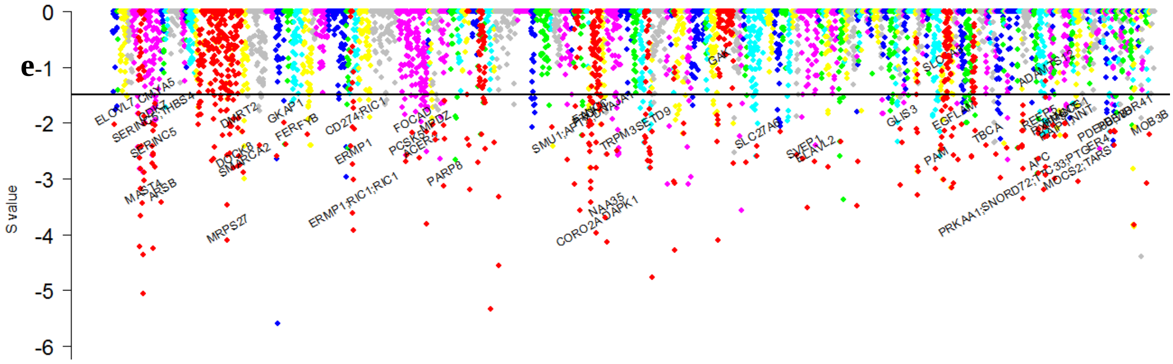
S value

f

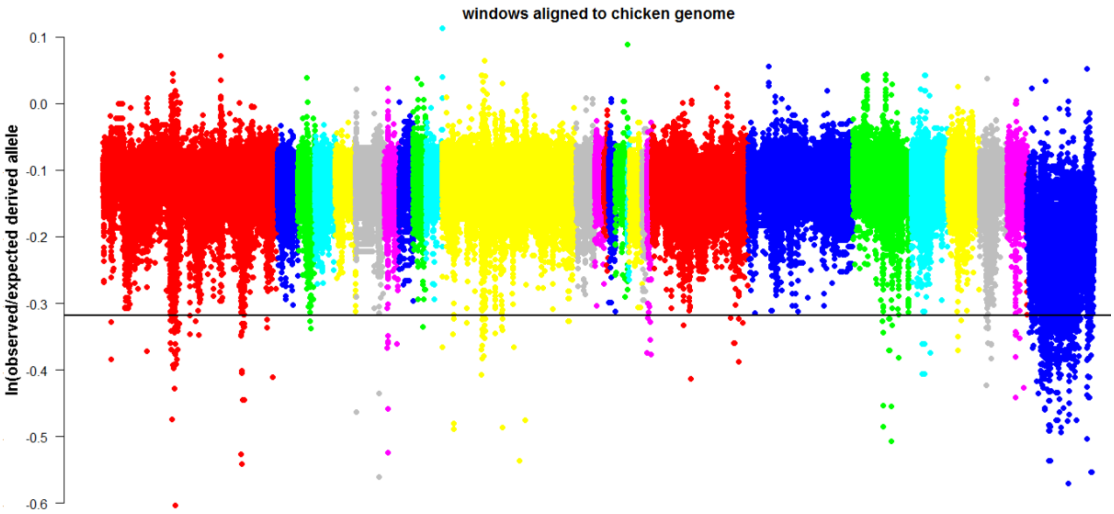
windows aligned to chicken genome
ln(observed/expected derived allele

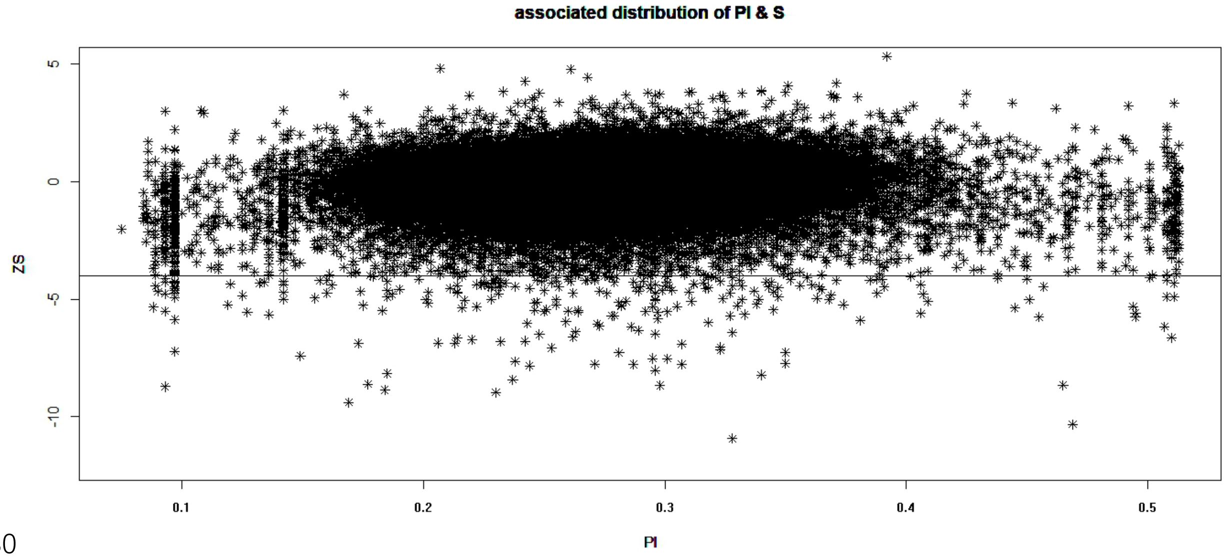

**Fig. S20. Whole genome screen for S score in wild duck with 20 kb window sliding by 10 kb.**
(**a**) Correlation of observed derived allele frequency between wild (combine mallard and spot-billed) and domestic duck (combine all 8 domestic duck breeds) lineage. (**b**) Distribution of the ln(observed/expected derived alleles frequency) for autosomes and Z-chromosome. (**c**) Distribution of the S score for autosomes and Z-chromosome. (**d**) The negative end of the S score distribution presented along duck autosomes. Genes that located in or near selected regions were showed its symbol. (**e**) The negative end of the S score distribution presented along duck Z-chromosome. (**f**) The S score distribution presented along scaffolds that is arranged according to their alignment to the chicken genome. (**g**) S Z-score of windows vs its nucleotide diversity. Dots blow horizontal lines represented windows of candidate ancient selection regions.

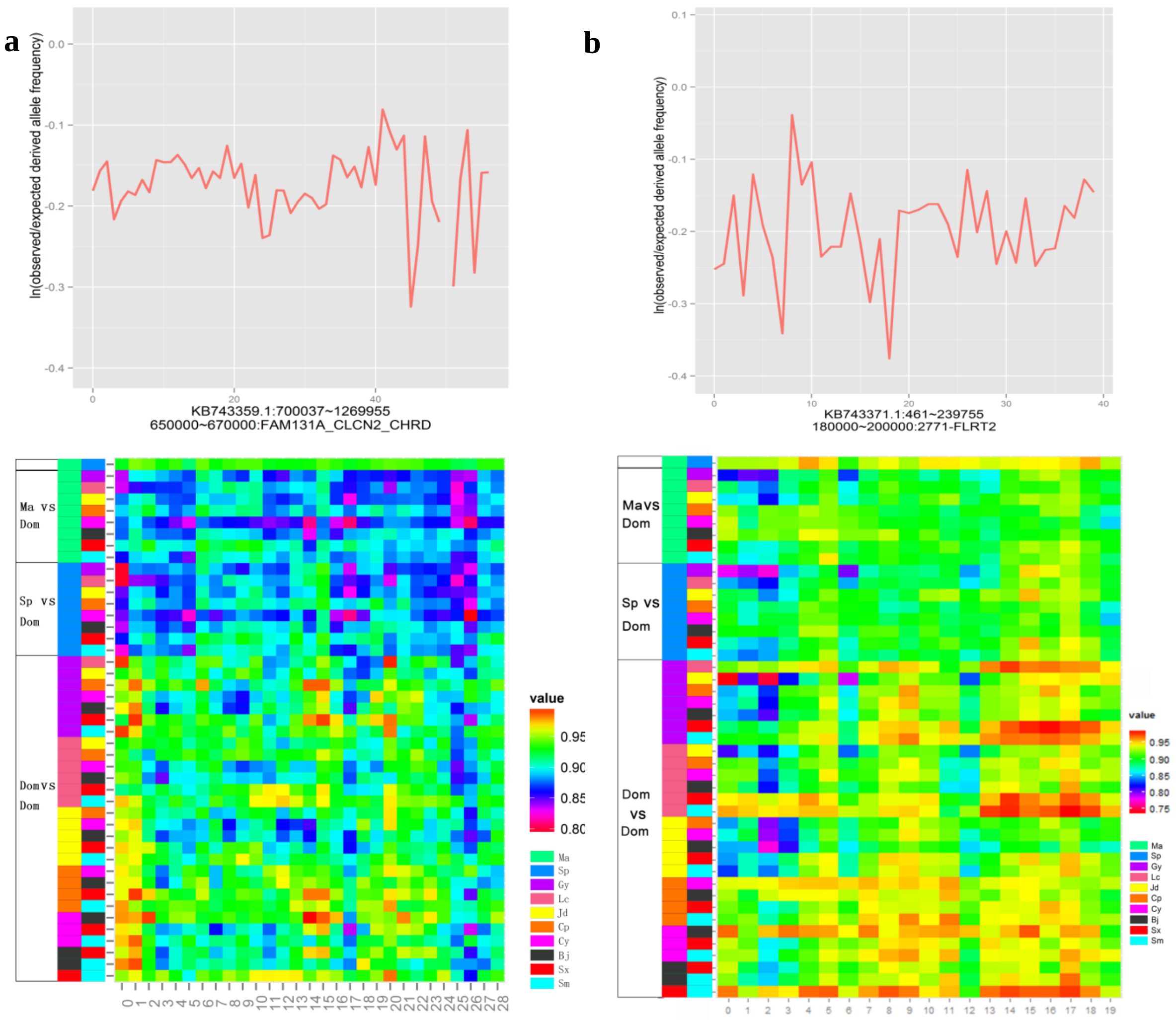
a
ln(observed/expected derived allele frequency)
KB743359.1:700037~1269955
650000~670000:FAM131A_CLCN2_CHRD
Ma vs Dom
Sp vs Dom
Dom vs Dom
value
Ma
Sp
Gy
Lc
Jd
Cp
Cy
Bj
Sx
Sm
b
ln(observed/expected derived allele frequency)
KB743371.1:461~239755
180000~200000:2771-FLRT2
Ma vs Dom
Sp vs Dom
Dom vs Dom
value

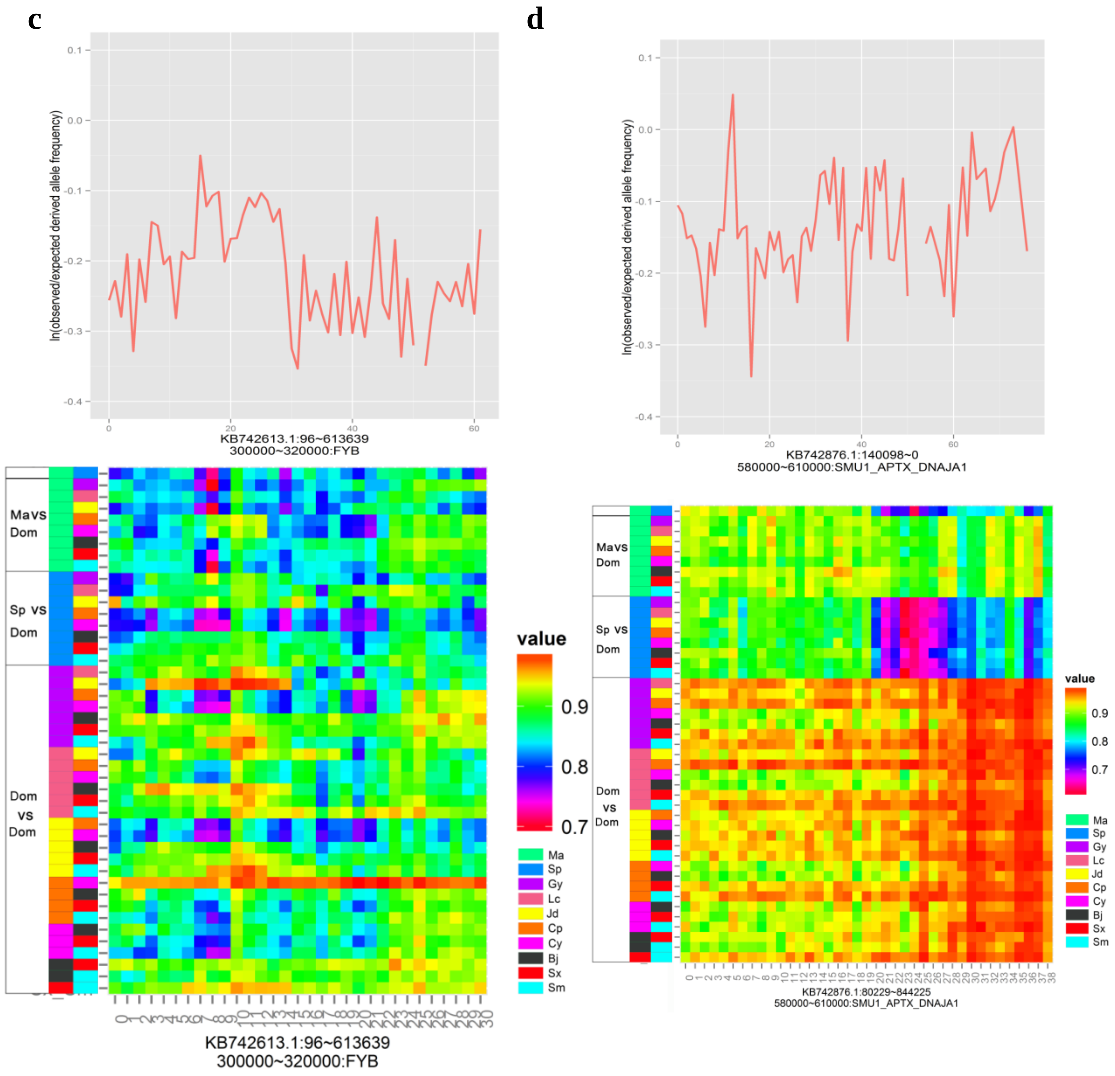

**Fig. S21. Details of four putative selective sweep regions in the early stage of the wild duck lineage.**

Each panel shows the distribution of the log-transformed value of the ratio for the observed/expected derive allele frequency using non-overlap 10 kb window consist (up) and scores between pairwise comparisons in 20 kb non-overlap windows (low). (**a**) *CHRD* locus. (**b**) *FLRT2* locus. (**c**) *FYB* locus. (**d**) *DNAJA1* locus.

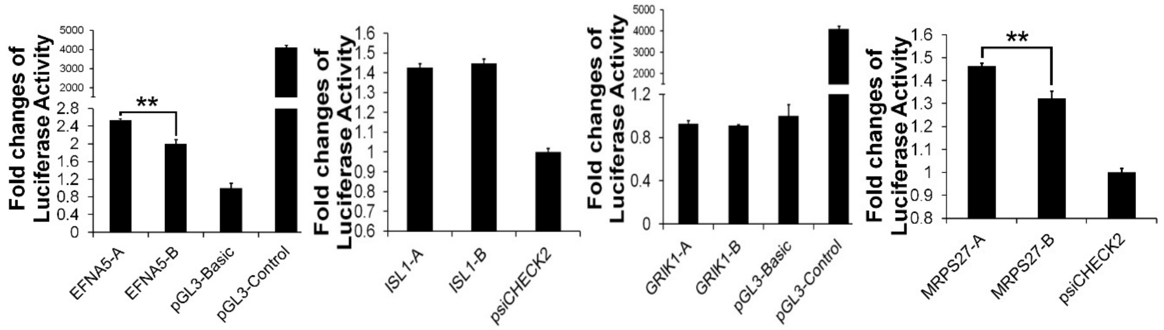

**Fig. S22. Effects of casual candidate mutation on luciferase reporter activity.** Comparison in luciferase activity of alleles in wild (spot-billed and mallard) and domestic (Beijing) duck in DF1 (chicken embryonic fibroblast) cells. The *ISL1* was under positive selection in early stage of the wild lineage. The *GRIK1* and *LYST* were under positive selection in the domestic lineage. “-A”, “-B” represent wild mutation types and domestic mutation types respectively. DF1 cells were transfected with recombinant pGL3 luciferase reporter plasmids and luciferase activity was measured at 24 hours after transfection.

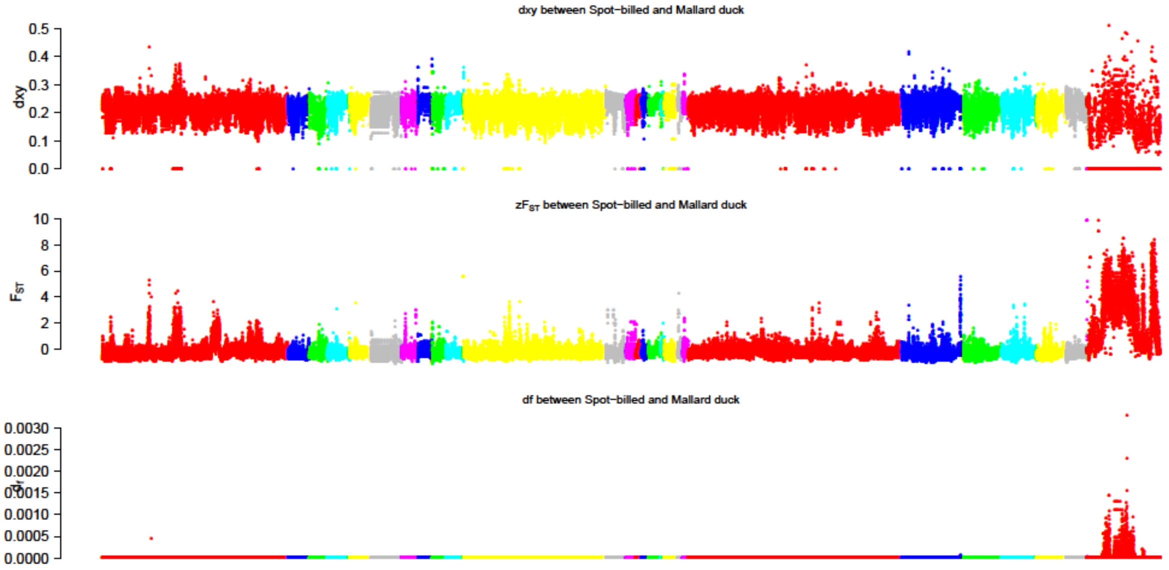

**Fig. S23. Distribution of $F_{ST}$ (fixation index), dxy and dfs (fixed difference sites) along Beijing duck scaffolds between mallard duck and spot-billed.**
$F_{ST}$ and dfs were counted with 40 kb window by step-size of 20 kb. The duck scaffolds that is shown in red and arranged at the right most are homologous to chicken Z chromosome and referred as duck Z chromosome.

**Fig. S24. Distribution of Tajima’D, positive end of z$F_{ST}$, dxy, number of SNPs per window, nucleotide diversity (π) and zHp along scaffolds in ducks.**
See “Fig. S23.docx”

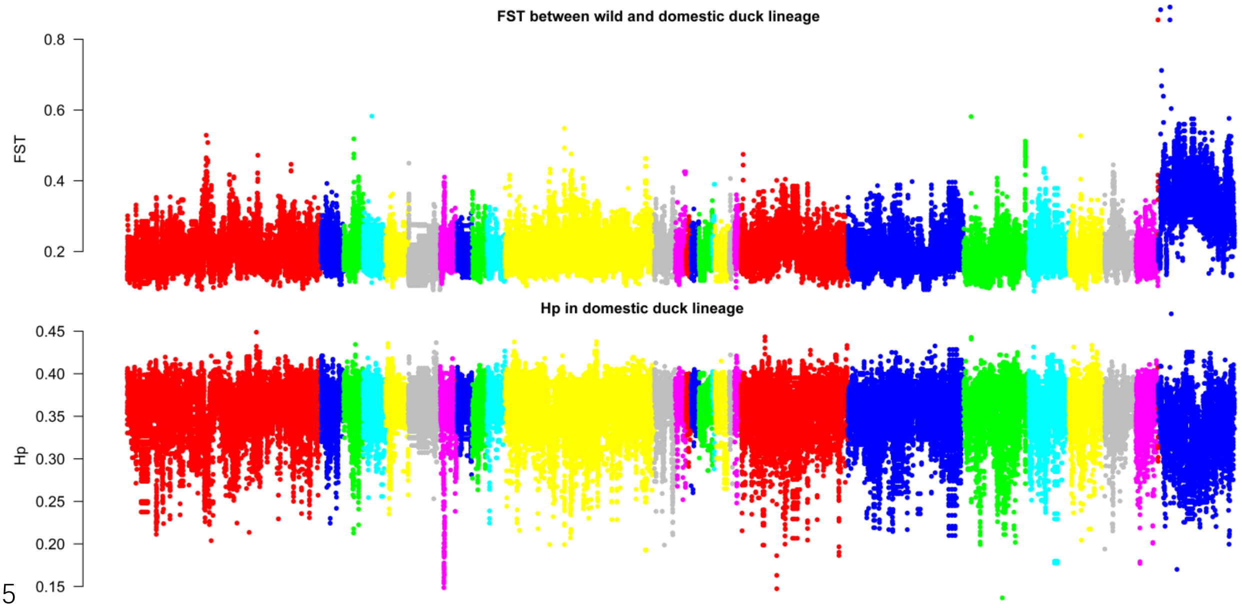

**Fig. S25. Distribution of $F_{ST}$ between the wild and the domestic duck lineage, and Hp in the domestic duck lineage.**

Both $F_{ST}$ and Hp were calculated in a 40 kb window with a sliding size of 20 kb along scaffolds that are arranged according their alignment to chicken autosomes and sex-chromosomes.

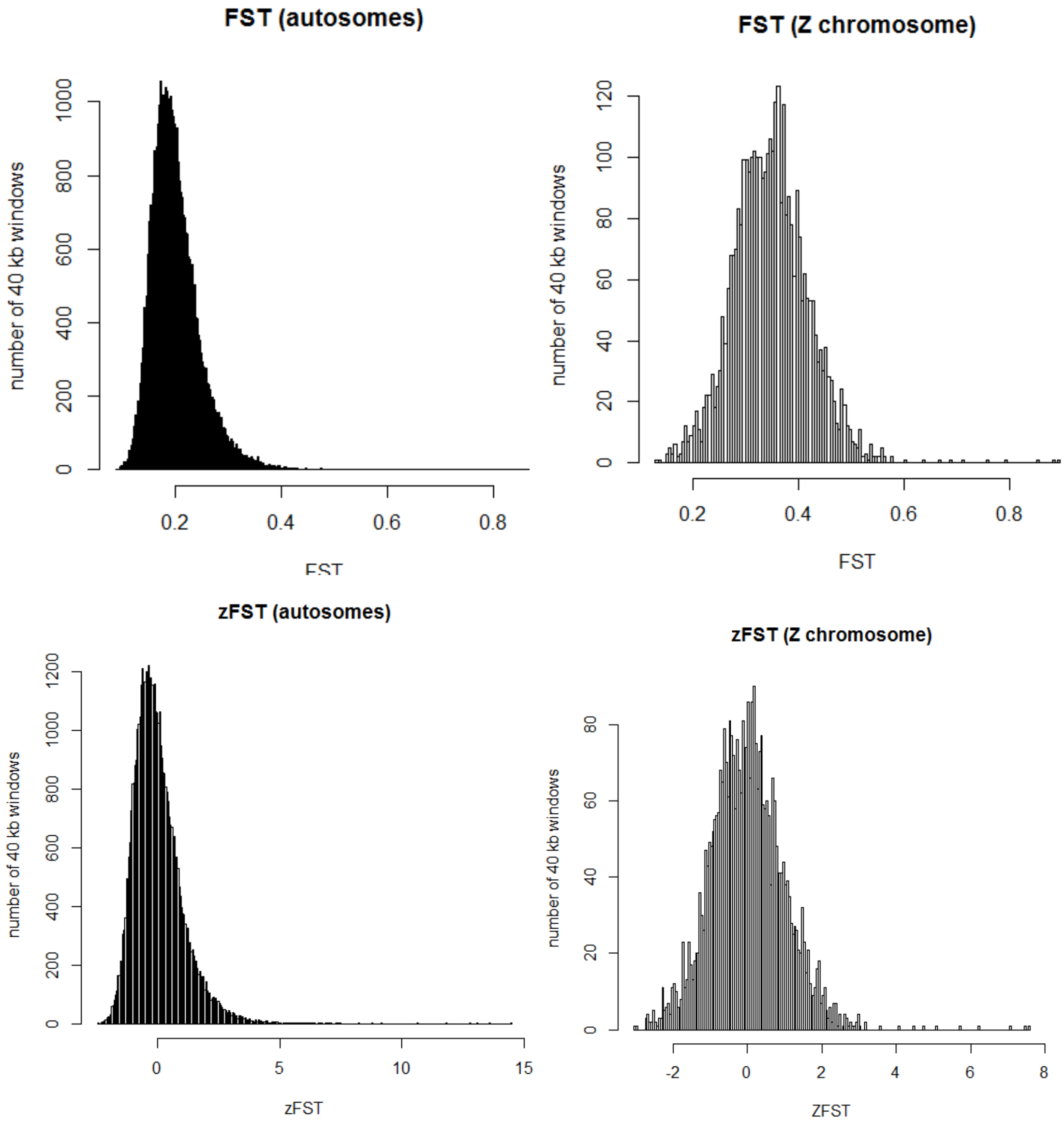

**Fig. S26. Distribution of $F_{ST}$/z$F_{ST}$ between the wild and the domestic duck lineage.**

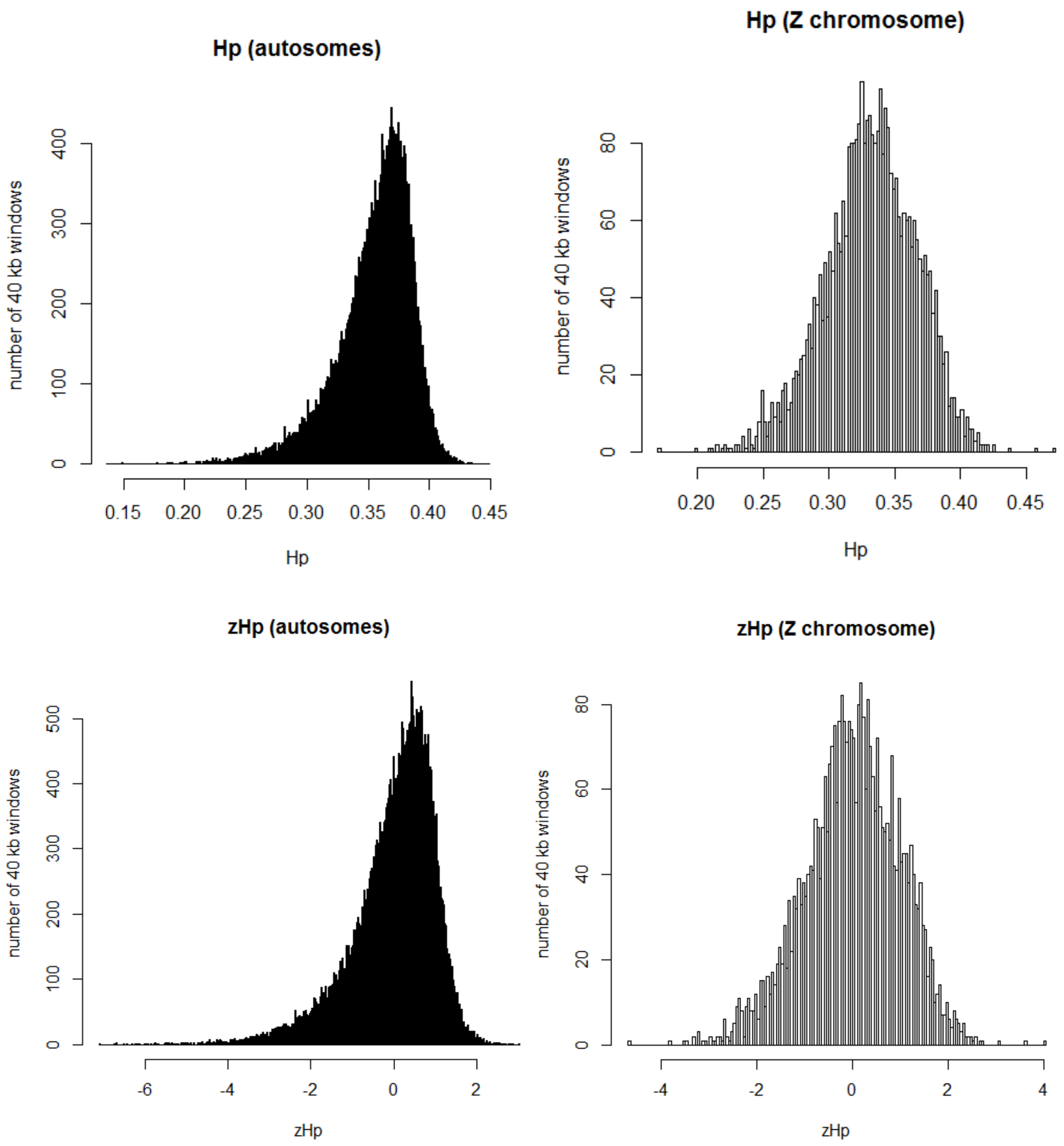

**Fig. S27. Distribution of Hp/zHp in the domestic duck lineage.**

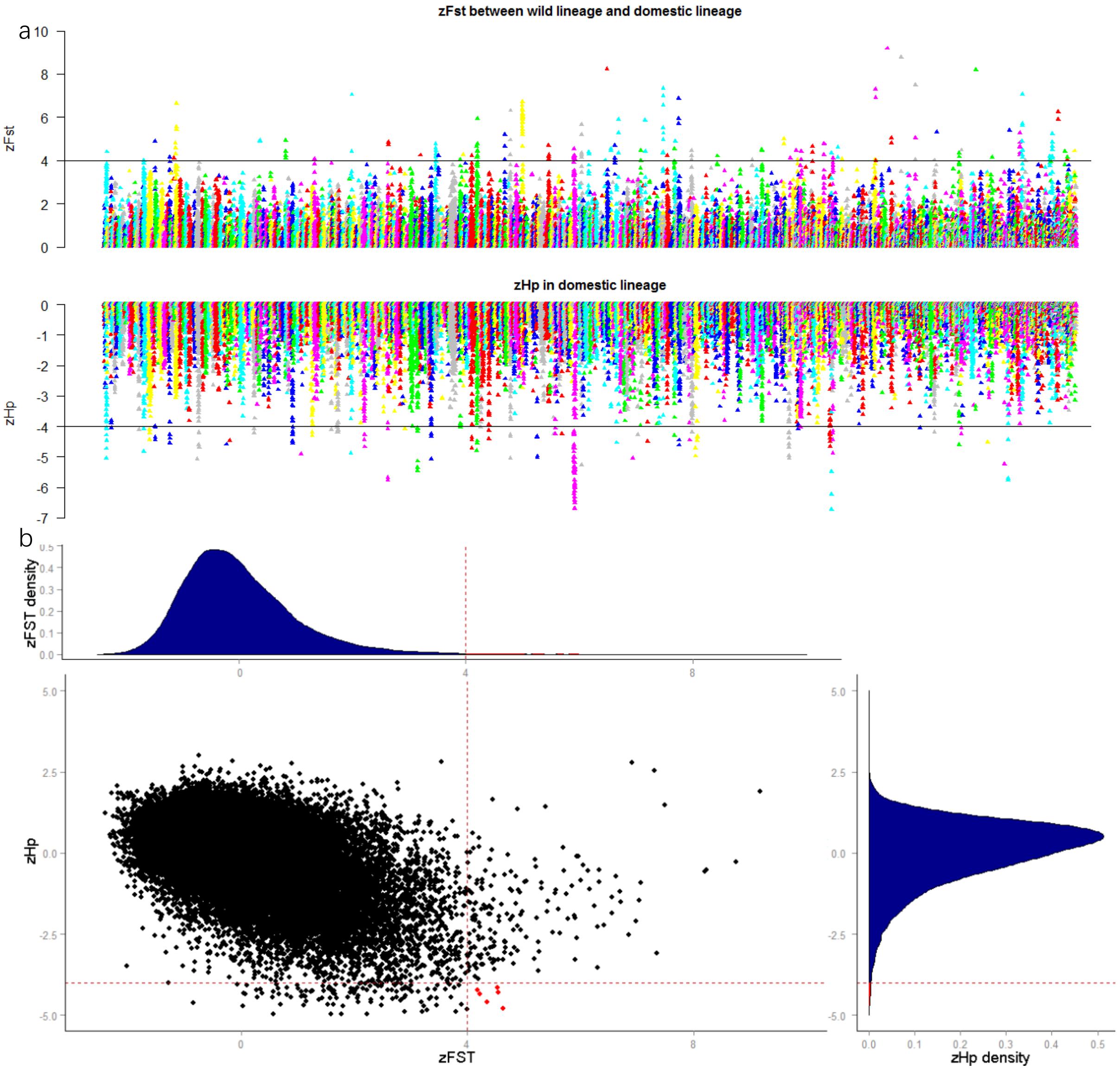

**Fig. S28. Distribution of z$F_{ST}$ and zHp in ducks.**

z$F_{ST}$ and zHp were counted with 40 kb window by a step size of 20 kb. (**a**) The positive end of the z$F_{ST}$ between wild lineage and domestic lineage and negative end of zHp in domestic lineage distribution plotted along scaffolds. (**b**) Distribution of z$F_{ST}$ and zHp. Red dash line indicate threshold and the windows identified as selection regions by both zHp and z$F_{ST}$ were show in red dots. (scaffolds homologous to chicken Z chromosome are not shown)

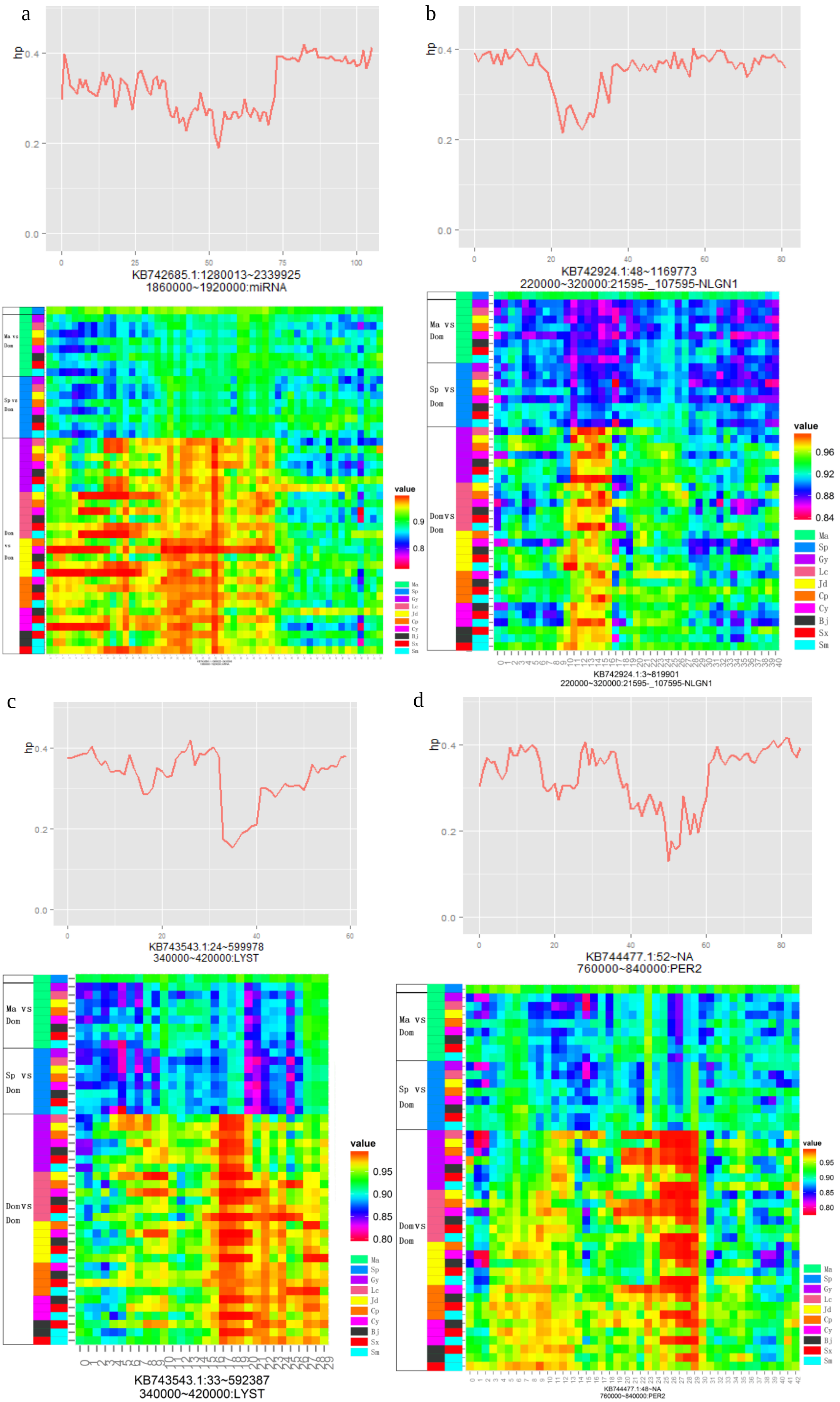
a
hp
KB742685.1:1280013~2339925
1860000~1920000:miRNA
b
KB742924.1:48~1169773
220000~320000:21595-_107595-NLGN1
value
Ma
Sp
Gy
Lc
Jd
Cp
Cy
Bj
Sx
Sm
Ma vs Dom
Sp vs Dom
Dom vs Dom
KB742924.1:3~819901
220000~320000:21595-_107595-NLGN1
c
KB743543.1:24~599978
340000~420000:LYST
KB743543.1:33~592387
340000~420000:LYST
d
KB744477.1:52~NA
760000~840000:PER2
KB744477.1:48~NA
760000~840000:PER2

**Fig. S29. Details of four putative selective sweep regions in the domestic duck lineage.**

Each panel shows distribution of the hp value using non-overlap 10 kb window (upside) and scores between pairwise comparisons in 20 kb non-overlap windows (downside). (**a**) *miR-3598-5p* locus. (**b**) *NLGN1* locus. (**c**) *LYST* locus. (**d**) *PER2* locus.

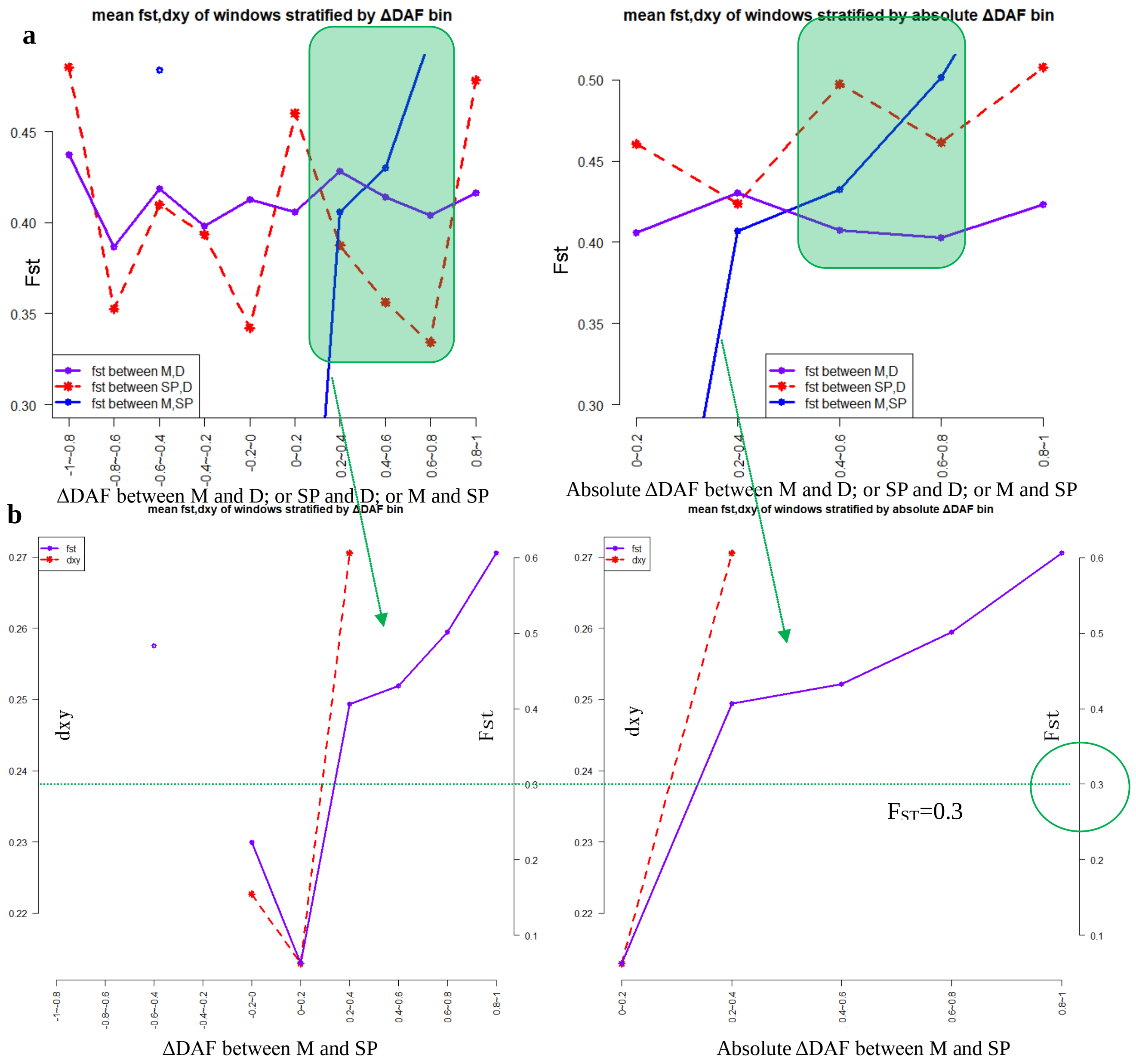

**Fig. S30. Observation on genomic divergence values stratified by ΔDAF provide evidence of introgression between domestic and wild ducks.**

**(a)** Mean $F_{ST}$ between domestic and wild duck populations decreased in high ΔDAF bins may due to introgression. Blue lines (corresponding to purple line in (**b**)) ΔDAF measured between mallard and spot-billed indicate lack of introgression evidence, when comparing with introgression cases that ΔDAF measured between wild (red dash line: spot-billed, purple line: mallard) and domestic ducks. “M”, “SP” and “D” indicate mallard, spot-billed and domestic duck, respectively. **(b)** Mean $F_{ST}$ (purple lines), dxy (red dash line) of windows stratified by ΔDAF (absolute ΔDAF in right) bins between mallard and spot-billed duck populations. dxy is not available in higher ΔDAF bins due to fixed SNPs are exclude in dxy calculation.

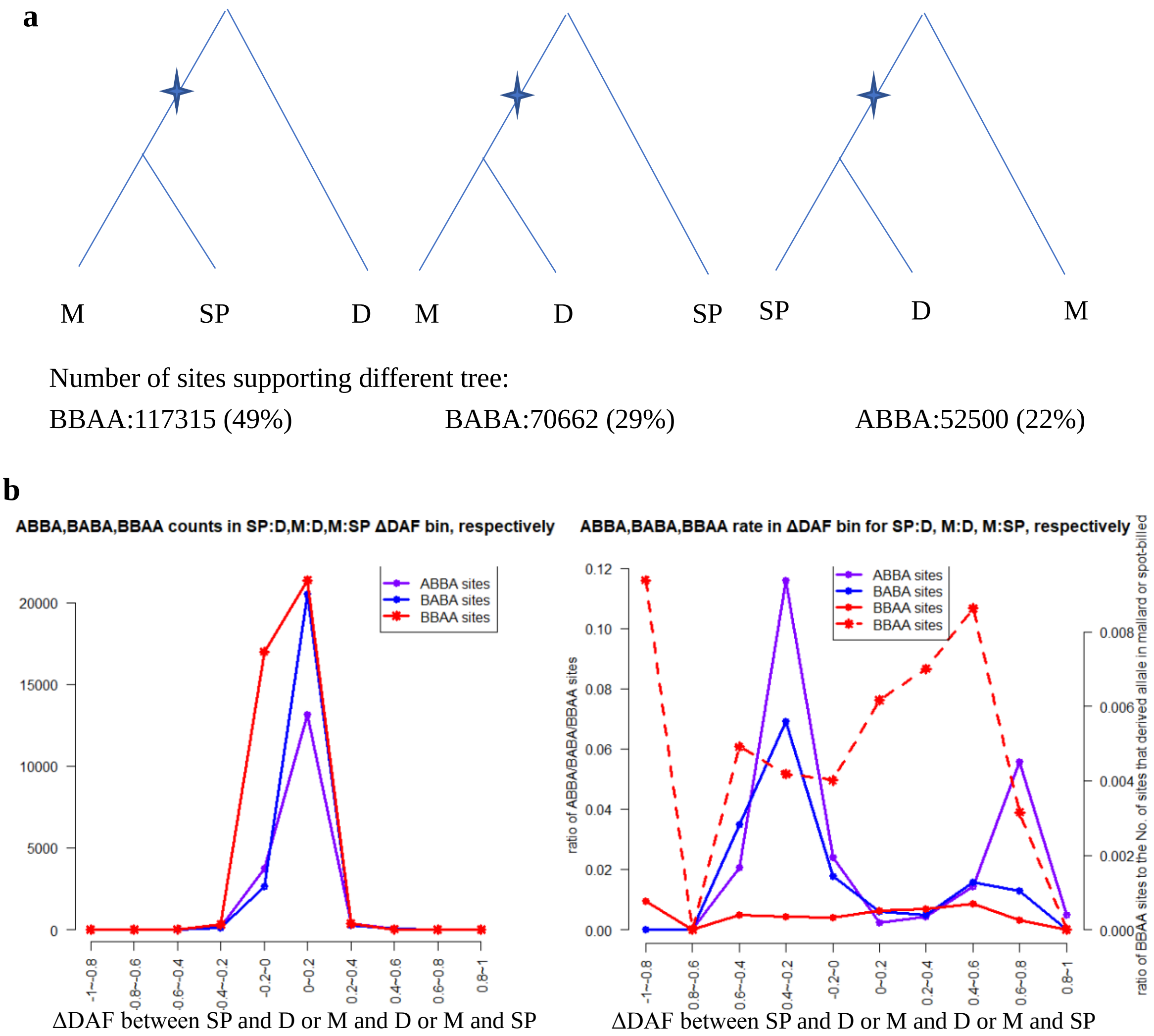

**Fig. S31. Absent evidence of introgression between mallard duck and spot-billed duck population based on ABBA-BABA analysis of mallard, spot-billed and domestic duck.**

**(a)** Number of sites supporting different trees is indicated both as a percentage and as actual numbers. **(b)** Left panel, number of ABBA, BABA and BBAA sites were stratified by ΔDAF between spot-billed: domestic (purple line), mallard: domestic (blue line) and mallard: spot-billed (red line), respectively. Right panel, BBAA rate (red line) were measured as the BBAA counts divided by number of sites that derived allele present in either mallard or spot-billed population in each bins (ABBA (purple line), BABA (blue line) similarly). The red dash line is corresponding to right y-axis. "M", "SP" and "D" indicate mallard, spot-billed and domestic duck, respectively.

**Supplementary Tables**

**Table S1: Summary for whole-genome resequencing**

| Group | Species/breeds | Sample number | Sequence depth | Total data (Gb) | Read length | Sequence strategy | SNP number |
|---|---|---|---|---|---|---|---|
| Wild duck | Mallard | 14 | 14.1× | 218.2 | 125 | Individual | 32,982,331 |
| | Spot-billed | 13 | 14.5× | 209.5 | 125 | Individual | 33,581,513 |
| | Wigeon | 3 | 20× | 75.6 | 125 | Individual | 21,393,824 |
| | Gadwall | 1 | 20× | 17.0 | 125 | Individual | 10,756,767 |
| | Pintail | 1 | 20× | 25.7 | 125 | Individual | 8,551,170 |
| | Falcata teal | 1 | 20× | 22.3 | 125 | Individual | 7,658,519 |
| | Common teal | 1 | 20× | 22.1 | 125 | Individual | 9,146,292 |
| | Baikal teal | 1 | 20× | 23.2 | 125 | Individual | 8,500,280 |
| Indigenous Chinese duck breeds | Beijing | 27 | 13.5× | 400.7 | 125 | Individual | 14,748,500 |
| | Shaoxing | 27 | 2~3.5× | 95.1 | 100 | Individual | 15,414,890 |
| | Gaoyou | 10 | 4.7× | 56.9 | 100 | Pooling | 9,190,980 |
| | Shanma | 10 | 4.0× | 50.4 | 100 | Pooling | 10,185,166 |
| | Lianchengbai | 10 | 4.7× | 57.6 | 100 | Pooling | 8,978,662 |
| | Jinding | 10 | 4.4× | 55.5 | 100 | Pooling | 8,617,344 |
| Commercial breeds | Cherry valley | 10 | 7.5× | 94.5 | 100 | Pooling | 9,615,871 |
| | Campbell | 10 | 4.5× | 57.3 | 100 | Pooling | 9,821,965 |
| Outgroup | Muscovy | 10 | 5.24× | 57.9 | 100 | Pooling | -- |
| | Taihu goose | 10 | 3.5× | 39.7 | 100 | Pooling | -- |

**Table S2: Statistics of the reads in mallard and spot-billed genome assemblies**

| Paired-end libraries (bp) | Paired-end insert size | Number of libraries | Average read length (bp) | Total data (Gb) | Sequence coverage (X) | Physical coverage (X) |
|---|---|---|---|---|---|---|
| mallard | 250bp | 3 | 150_150 | 89.76 | 70.81 | 59.01 |
| | 500bp | 2 | 100_100 | 46.18 | 36.94 | 92.35 |
| | 800bp | 2 | 100_100 | 57.4 | 45.92 | 183.68 |
| | 2K | 3 | 49_49 | 27.37 | 21.9 | 446.94 |
| | 5K | 2 | 49_49 | 16.32 | 13.06 | 666.33 |
| | 10K | 3 | 49_49 | 11.67 | 9.34 | 953.06 |
| | 20K | 2 | 49_49 | 13.75 | 11 | 2,244.90 |
| Total | / | 17 | / | 262.45 | 208.97 | 4,646.27 |
| spot-billed | 250bp | 3 | 150_150 | 95.64 | 76.51 | 63.76 |
| | 500bp | 2 | 100_100 | 63.39 | 50.71 | 126.78 |
| | 800bp | 2 | 100_100 | 45.31 | 36.25 | 145 |

| | | | | | | |
|---|---|---|---|---|---|---|
| | 2K | 3 | 49_49 | 26.2 | 20.96 | 427.76 |
| | 5K | 2 | 49_49 | 17.11 | 13.69 | 698.47 |
| | 10K | 3 | 49_49 | 15.37 | 12.3 | 1,255.10 |
| | 20K | 2 | 49_49 | 12.17 | 9.74 | 1,987.76 |
| Total | / | 17 | / | 275.19 | 220.16 | 4,704.63 |

**Table S3: Statistics of the three *de novo* ducks assemblies**

| Category | Contig | | | Scaffold | | |
|---|---|---|---|---|---|---|
| | Beijing Duck | Mallard | Spot-billed | Beijing Duck | Mallard | Spot-billed |
| Total length(kb) | 1,069,961 | 1,176,969 | 1,187,333 | 1,105,049 | 1,270,808 | 1,317,385 |
| Total number | 227,597 | 172,189 | 194,934 | 78,487 | 61,591 | 67,685 |
| Longest size(bp) | 263,737 | 484,574 | 673,122 | 5,998,093 | 17,658,973 | 25,320,044 |
| N50 size(bp) | 26,138 | 38,271 | 33,061 | 1,233,631 | 2,489,142 | 2,054,876 |
| N50 number | 11,193 | 7,876 | 9,194 | 268 | 125 | 154 |
| N90 size(bp) | 3,085 | 5,381 | 4,386 | 195,458 | 145,390 | 102,224 |
| N90 number | 53,850 | 38,114 | 45,269 | 1,097 | 870 | 1,035 |

**Table S4: Comparison in length and coverage of the mallard and the spot-billed genome using seven Beijing duck BAC sequences**

| BACs name | Length of BACs (bp) | mallard coverage | spot-billed coverage |
|---|---|---|---|
| ndae_TLR1 | 67,954 | 85.91% | 70.03% |
| ndbaa_TLR15 | 79,704 | 98.50% | 98.25% |
| ndbab_TLR5 | 72,128 | 98.41% | 98.32% |
| ndbac_1_TLR3 | 76,489 | 97.53% | 96.49% |
| ndbad_2_TLR7 | 97,799 | 95.64% | 87.51% |
| ndbae_TLR2 | 75,080 | 98.82% | 93.59% |
| ndbag_TLR4 | 77,042 | 97.79% | 97.98% |

**Table S5: Evaluation of the mallard genome coverage using duck EST sequences**

| Dataset | Number | Total length (bp) | Covered by assembly (%) | with >90% sequence in one scaffold | | with >50% sequence in one scaffold | |
|---|---|---|---|---|---|---|---|
| | | | | Number | Percent (%) | Number | Percent (%) |
| >0bp | 319,996 | 98,257,350 | 99.02 | 294,845 | 92.14 | 315,689 | 98.65 |

| Dataset | Number | Total length (bp) | Covered by assembly (%) | with >90% sequence in one scaffold | | with >50% sequence in one scaffold | |
|---|---|---|---|---|---|---|---|
| | | | | Number | Percent (%) | Number | Percent (%) |
| >200bp | 144,337 | 73,939,509 | 99.53 | 135,799 | 94.08 | 143,128 | 99.16 |
| >500bp | 43,896 | 43,155,260 | 99.84 | 41,595 | 94.76 | 43,687 | 99.52 |
| >1000bp | 13,652 | 22,939,026 | 99.99 | 12,831 | 93.99 | 13,613 | 99.71 |

**Table S6: Evaluation of the spot-billed genome coverage using duck EST sequences**

| Dataset | Number | Total length (bp) | Covered by assembly (%) | with >90% sequence in one scaffold | | with >50% sequence in one scaffold | |
|---|---|---|---|---|---|---|---|
| | | | | Number | Percent (%) | Number | Percent (%) |
| >0bp | 319,996 | 98,257,350 | 99.01 | 294,324 | 91.98 | 315,566 | 98.62 |
| >200bp | 144,337 | 73,939,509 | 99.53 | 135,565 | 93.92 | 143,053 | 99.11 |
| >500bp | 43,896 | 43,155,260 | 99.85 | 41,531 | 94.61 | 43,670 | 99.49 |
| >1000bp | 13,652 | 22,939,026 | 99.99 | 12,802 | 93.77 | 13,606 | 99.66 |

**Table S7: Type and proportion of transposable elements(TEs) in the Mallard, Spot-billed, Beijing duck and chicken genome**

| TE type | mallard length (bp) | % genome | Spot-billed length (bp) | % genome | Beijing duck length (bp) | % genome | Chicken length (bp) | % genome |
|---|---|---|---|---|---|---|---|---|
| DNA | 9,479,699 | 0.75 | 9,438,606 | 0.71 | 5,590,903 | 0.51 | 13,872,990 | 1.33 |
| LINE | 98,060,422 | 7.71 | 101,177,399 | 7.68 | 63,146,176 | 5.71 | 116,778,031 | 11.15 |
| SINE | 1,389,649 | 0.1 | 1,274,941 | 0.09 | 892,886 | 0.08 | 573,610 | 0.05 |
| LTR | 52,195,974 | 4.1 | 48,381,075 | 3.67 | 21,255,312 | 1.92 | 47,374,647 | 4.53 |
| Other | 21,136 | 0 | 12,303 | 0 | 3,201 | 0 | 3,338 | 0 |
| Unknown | 1,100,812 | 0.08 | 3,402,377 | 0.25 | 4,600,535 | 0.41 | 3,351,756 | 0.32 |
| Total | 136,134,153 | 10.71 | 139,051,537 | 10.55 | 80,740,993 | 7.31 | 149,208,388 | 14.25 |

**Table S8: Gene annotation of the mallard and spot-billed reference gene sets using five protein databases**

| Database | Mallard | | Spot-billed | |
|---|---|---|---|---|
| | Number | Percent (%) | Number | Percent (%) |
| Total | 21,056 | 100 | 21,123 | 100 |
| InterPro | 15,335 | 72.83 | 15,308 | 72.47 |
| GO | 12,817 | 60.87 | 12,815 | 60.67 |
| KEGG | 15,878 | 75.41 | 15,803 | 74.81 |
| Swissprot | 17,551 | 83.35 | 17,531 | 82.99 |
| TrEMBL | 18,371 | 87.25 | 18,327 | 86.76 |
| All annotated | 18,467 | 87.70 | 18,426 | 87.23 |

| | | | | |
|---|---|---|---|---|
| Unannotated | 2,589 | 12.29 | 2,697 | 12.77 |

**Table S9: Gene family of two mammals and seven birds clustering by TreeFam**

| Species | Total genes | Unclustered genes | Families | Unique families | Ave. genes per family |
|---|---|---|---|---|---|
| mallard | 21,056 | 2,478 | 14,541 | 30 | 1.28 |
| Spot-billed | 21,123 | 2,604 | 14,530 | 42 | 1.27 |
| Beijing duck | 20,629 | 5,381 | 13,498 | 18 | 1.13 |
| *G. gallus* | 15,508 | 493 | 13,351 | 6 | 1.12 |
| *S. camelus* | 16,178 | 326 | 11,579 | 105 | 1.37 |
| *M. gallopavo* | 14,123 | 455 | 12,414 | 7 | 1.10 |
| *F. albico* | 15,303 | 716 | 13,284 | 33 | 1.10 |
| *H. sapiens* | 20,238 | 1,825 | 15,036 | 117 | 1.22 |
| *M. musculus* | 22,627 | 1,549 | 15,407 | 139 | 1.37 |

**Table S10: Over represented GO terms (p-value <0.001) of wild duck expanded gene families**

See file “Table_S9.xlsx”

**Table S11: Tests for population mixture of mallard, spot-billed, Beijing and Shaoxing ducks**

| Source 1 | Source 2 | Target | f3 | Standard error | Z | SNPs |
|---|---|---|---|---|---|---|
| Mallard | Spot-billed | Beijing | 0.007886 | 0.000748 | 10.546 | 9,997 |
| Mallard | Spot-billed | Shaoxing | 0.008662 | 0.000814 | 10.643 | 9,943 |
| Mallard | Spot-billed | Mix Beijing, Shaoxing | 0.007279 | 0.000305 | 23.834 | 2,930,914 |
| Mallard_hz | Spot-billed_hz | Beijing | 0.008656 | 0.001076 | 8.045 | 9,975 |
| Mallard_hz | Spot-billed_hz | Shaoxing | 0.010200 | 0.000683 | 14.930 | 9,907 |
| Mallard_fh | Spot-billed_fh | Beijing | 0.007320 | 0.000603 | 12.147 | 9,986 |
| Mallard_fh | Spot-billed_fh | Shaoxing | 0.007380 | 0.000988 | 7.470 | 9,921 |

Considering the extensive substructure exist in wild ducks which reflect those individuals may trace their recent ancestry to different populations even species, we assume different clusters combinations to source 1 and source 2, and combine different domestic breeds to one target population, to minimize the population effect. All the Z were significantly greater than zero.

**Table S12: Divergence without migration inferred parameters.**

| combinations | Parameter[a] | Conventional bootstrap 95% confidence interval | Conventional bootstrap 95% confidence interval (convert to real) | Maximum likelihood |
|---|---|---|---|---|
| Domestic, mallard duck | s (Nmallard) | 0.85658-0.874169 | 1567694-1637111 | 1595689 |
| | 1-s(Ndomestic) | 0.125831-0.143419 | 233297-265236 | - |

| | | | | |
|---|---|---|---|---|
| | Ts | 0.0272-0.031008 | 100818-114733 | 102427 |
| Domestic, spot-billed duck | s(Nspot-bill) | 0.8776821-0.893130 | 1660302-1731619 | 1659603 |
| | 1-s(Ndomestic) | 0.10687-0.122315 | 205404-233549 | - |
| | Ts | 0.02321-0.02653 | 89220-101346 | 91700 |
| Spot-billed, mallard duck | s(Nmallard) | 0.4352336-0.472079 | 949555-1031148 | 969384 |
| | 1-s(Nspot~) | 0.52792-0.564766 | 1146439- 1239046 | - |
| | Ts | 0.015755-0.0174258 | 68583-76294 | 69432 |

[a] see model code above.

**Table S13: Summary of selective sweep regions in the wild duck lineage**

These selective sweep regions occurred during their population in conjunction with or shortly after, their population divergence from domestic lineage (S<-4 for autosomes, S<-1.5 for Z-chromosomes). See file “Table_S13.xlsx”

**Table S14: Enriched gene ontology of positively selected genes in the wild duck lineage.**
See file “Table_S14.xlsx”

**Table S15: Enriched gene ontology in biological process terms of positively selected genes in wild duck lineage.**
See file “Table_S15.xlsx”

**Table S16: Information of variant sites being performed luciferase assay to evaluate the activity.**

| Gene symbol | Variant position | Catalog of variant position | ΔAF | AF in wild | AF in domestic |
|---|---|---|---|---|---|
| *EFNA5* | KB744645.1: 114292,114293,114301 | 10 kb* | 0.949,0.916,0.740 | 0,0.916,0 | 0.949,0,0.741 |
| *EFNA5* | KB744645.1:58363 | $1^{th}$ intron | 0.809 | 0.835 | 0.026 |
| *ISL1* | KB743873.1:25803 (indel) | 3’ UTR | 0.875 | 0.875 | 0 |
| *MRPS27* | KB742554.1: 2151394,2151434,2151445 | 3’ UTR | 0.960,0.419,0.417 | 0,0.419,0.419 | 0.960,0.419,0.419 |
| *GRIK1* | KB742951.1:543269 | $2^{th}$ intron | 0.730 | 0.151 | 0.881 |
| *LYST* | KB743543.1:403502 | $5^{th}$ intron | 0.611 | 0.685 | 0.074 |

Note: 10 kb* means 10 kb downstream of the gene.

**Table S17: Mean values of population genomic parameters for Z-chromosome and autosomes in mallard and spot-billed**

| Index | | Autosomes | | Z-chromosome | |
|---|---|---|---|---|---|
| | | mean | SD | Mean | SD |
| Pi | Mallard | 0.007520 | 0.004199 | 0.003397 | 0.002495 |
| | Spot-billed | 0.007607 | 0.004321 | 0.003020 | 0.002625 |
| SNPs | Mallard | 1,123 | 572 | 494 | 348 |
| | Spot-billed | 1,150 | 600 | 440 | 376 |
| Tajima's D | Mallard | -0.525593 | 0.435471 | -0.257150 | 0.445641 |
| | Spot-billed | -0.625759 | 0.420070 | -0.348016 | 0.492465 |
| $F_{ST}$ | | 0.049303 | 0.034864 | 0.259623 | 0.150430 |

**Table S18: Mean values of genomic parameters inside/outside high $F_{ST}$ (between mallard and spot-billed ducks) regions in autosomes**

| Species | | pi (π) | | No. of SNPs | | Tajima's D | |
|---|---|---|---|---|---|---|---|
| | | mean | sd | mean | sd | mean | sd |
| Mallard | Inside | 0.002421 | 0.00217898 | 360 | 305 | -0.61100 | 0.579419 |
| | Outside | 0.007595 | 0.00418171 | 1,134 | 569 | -0.52064 | 0.433172 |
| Spot-billed | Inside | 0.002331 | 0.00196792 | 334 | 280 | -0.47211 | 0.656325 |
| | Outside | 0.007683 | 0.00430549 | 1,161 | 597 | -0.6228 | 0.415501 |

**Table S19: Summary of domestic candidate regions (zHp<-4 for autosomes, zHp<-2 for Z-chromosomes)**
See file “Table_S19.xlsx”

**Table S20: Summary of high $F_{ST}$ regions between the wild and domestic duck lineage.**
See file “Table_S20.xlsx”

**Table S21: Enriched gene ontology of positively selected genes in/close to domestic candidate regions.**
See file “Table_S21.xlsx”

**Table S22: Enriched gene ontology in biological process terms of positively selected genes in/close to domestic candidate regions.**
See file “Table_S22.xlsx”

**Table S23. Averaged all ΔDAF (absolute ΔDAF) of windows inside/outside high differentiation regions.**

| Mean value across SNPs | high differentiation regions | Genomic background |
|---|---|---|
| absolute ΔDAF | 0.0385672 | 0.03672404 |

| | | |
|---|---|---|
| ΔDAF | 0.0370516 | 0.03497989 |

#### Reference

Anderson, R. M., Lawrence, A. R., Stottmann, R. W., Bachiller, D., & Klingensmith, J. (2002). Chordin and noggin promote organizing centers of forebrain development in the mouse. *Development, 129*(21), 4975-4987.

Ashburner, M., Ball, C. A., Blake, J. A., Botstein, D., Butler, H., Cherry, J. M., . . . Sherlock, G. (2000). Gene ontology: tool for the unification of biology. The Gene Ontology Consortium. *Nat Genet, 25*(1), 25-29. doi:10.1038/75556

Axelsson, E., Ratnakumar, A., Arendt, M. L., Maqbool, K., Webster, M. T., Perloski, M., . . . Lindblad-Toh, K. (2013). The genomic signature of dog domestication reveals adaptation to a starch-rich diet. *Nature, 495*(7441), 360-364. doi:10.1038/nature11837

Barber, M. R., Aldridge, J. R., Jr., Webster, R. G., & Magor, K. E. (2010). Association of RIG-I with innate immunity of ducks to influenza. *Proc Natl Acad Sci U S A, 107*(13), 5913-5918. doi:10.1073/pnas.1001755107

Benson, G. (1999). Tandem repeats finder: a program to analyze DNA sequences. *Nucleic Acids Res, 27*(2), 573-580.

Birney, E., Clamp, M., & Durbin, R. (2004). GeneWise and Genomewise. *Genome Res, 14*(5), 988-995. doi:10.1101/gr.1865504

Chen, N. (2004). Using RepeatMasker to identify repetitive elements in genomic sequences. *Curr Protoc Bioinformatics, Chapter 4*, Unit 4 10. doi:10.1002/0471250953.bi0410s05

Danecek, P., Auton, A., Abecasis, G., Albers, C. A., Banks, E., DePristo, M. A., . . . Genomes Project Analysis, G. (2011). The variant call format and VCFtools. *Bioinformatics, 27*(15), 2156-2158. doi:10.1093/bioinformatics/btr330

de Manuel, M., Kuhlwilm, M., Frandsen, P., Sousa, V. C., Desai, T., Prado-Martinez, J., . . . Marques-Bonet, T. (2016). Chimpanzee genomic diversity reveals ancient admixture with bonobos. *Science, 354*(6311), 477-481. doi:10.1126/science.aag2602

DePristo, M. A., Banks, E., Poplin, R., Garimella, K. V., Maguire, J. R., Hartl, C., . . . Daly, M. J. (2011). A framework for variation discovery and genotyping using next-generation DNA sequencing data. *Nat Genet, 43*(5), 491-498. doi:10.1038/ng.806

Edgar, R. C. (2004). MUSCLE: multiple sequence alignment with high accuracy and high throughput. *Nucleic Acids Res, 32*(5), 1792-1797. doi:10.1093/nar/gkh340

Ellegren, H., Smeds, L., Burri, R., Olason, P. I., Backstrom, N., Kawakami, T., . . . Wolf, J. B. (2012). The genomic landscape of species divergence in Ficedula flycatchers. *Nature, 491*(7426), 756-760. doi:10.1038/nature11584

Elsik, C. G., Mackey, A. J., Reese, J. T., Milshina, N. V., Roos, D. S., & Weinstock, G. M. (2007). Creating a honey bee consensus gene set. *Genome Biol, 8*(1), R13. doi:10.1186/gb-2007-8-1-r13

Freedman, A. H., Gronau, I., Schweizer, R. M., Ortega-Del Vecchyo, D., Han, E., Silva, P. M., . . . Novembre, J. (2014). Genome sequencing highlights the dynamic early history of dogs. *PLoS Genet, 10*(1), e1004016. doi:10.1371/journal.pgen.1004016

Green, R. E., Krause, J., Briggs, A. W., Maricic, T., Stenzel, U., Kircher, M., . . . Paabo, S. (2010). A draft sequence of the Neandertal genome. *Science, 328*(5979), 710-722. doi:10.1126/science.1188021

Groenen, M. A., Archibald, A. L., Uenishi, H., Tuggle, C. K., Takeuchi, Y., Rothschild, M. F., . . . Megens, H.-J. (2012). Analyses of pig genomes provide insight into porcine demography and evolution. *Nature, 491*(7424), 393-398.

Han, F., Lamichhaney, S., Grant, B. R., Grant, P. R., Andersson, L., & Webster, M. T. (2017). Gene flow, ancient polymorphism, and ecological adaptation shape the genomic landscape of divergence among Darwin's finches. *Genome Res, 27*(6), 1004-1015. doi:10.1101/gr.212522.116

Harr, B. (2006). Genomic islands of differentiation between house mouse subspecies. *Genome Research, 16*(6), 730-737.

Harris, R. S. (2007). *Improved pairwise alignment of genomic DNA*: The Pennsylvania State University.

Huang, Y., Li, Y., Burt, D. W., Chen, H., Zhang, Y., Qian, W., . . . Li, J. (2013). The duck genome and transcriptome provide insight into an avian influenza virus reservoir species. *Nature genetics, 45*(7), 776-783.

Huang, Y., Li, Y., Burt, D. W., Chen, H., Zhang, Y., Qian, W., . . . Li, N. (2013). The duck genome and transcriptome provide insight into an avian influenza virus reservoir species. *Nat Genet, 45*(7), 776-783. doi:10.1038/ng.2657

Huang, Y., Temperley, N. D., Ren, L., Smith, J., Li, N., & Burt, D. W. (2011). Molecular evolution of the vertebrate TLR1 gene family--a complex history of gene duplication, gene conversion, positive selection and co-evolution. *BMC Evol Biol, 11*, 149. doi:10.1186/1471-2148-11-149

Huelsenbeck, J. P., & Ronquist, F. (2001). MRBAYES: Bayesian inference of phylogenetic trees. *Bioinformatics, 17*(8), 754-755.

Hunter, S., Apweiler, R., Attwood, T. K., Bairoch, A., Bateman, A., Binns, D., . . . Yeats, C. (2009). InterPro: the integrative protein signature database. *Nucleic Acids Res, 37*(Database issue), D211-215. doi:10.1093/nar/gkn785

Jurka, J., Kapitonov, V. V., Pavlicek, A., Klonowski, P., Kohany, O., & Walichiewicz, J. (2005). Repbase Update, a database of eukaryotic repetitive elements. *Cytogenet Genome Res, 110*(1-4), 462-467. doi:10.1159/000084979

Kanehisa, M., & Goto, S. (2000). KEGG: kyoto encyclopedia of genes and genomes. *Nucleic Acids Res, 28*(1), 27-30.

Karlsson, E. K., Baranowska, I., Wade, C. M., Salmon Hillbertz, N. H. C., Zody, M. C., Anderson, N., . . . Lindblad-Toh, K. (2007). Efficient mapping of mendelian traits in dogs through genome-wide association. *Nature genetics, 39*(11), 1321-1328. doi:10.1038/ng.2007.10

Kent, W. J. (2002). BLAT--the BLAST-like alignment tool. *Genome Res, 12*(4), 656-664. doi:10.1101/gr.229202. Article published online before March 2002

Kofler, R., Orozco-terWengel, P., De Maio, N., Pandey, R. V., Nolte, V., Futschik, A., . . . Schlotterer, C. (2011). PoPoolation: a toolbox for population genetic analysis of next generation sequencing data from pooled individuals. *PLoS One, 6*(1), e15925. doi:10.1371/journal.pone.0015925

Lavretsky, P., Dacosta, J. M., Hernandez-Banos, B. E., Engilis, A., Jr., Sorenson, M. D., & Peters, J. L. (2015). Speciation genomics and a role for the Z chromosome in the early stages of divergence between Mexican ducks and mallards. *Mol Ecol, 24*(21), 5364-5378. doi:10.1111/mec.13402

Li, C., Jiao, S., Wang, G., Gao, Y., Liu, C., He, X., . . . Wang, H. (2015). The Immune Adaptor ADAP Regulates Reciprocal TGF-beta1-Integrin Crosstalk to Protect from Influenza Virus Infection. *PLoS Pathog, 11*(4), e1004824. doi:10.1371/journal.ppat.1004824

Li, H., Handsaker, B., Wysoker, A., Fennell, T., Ruan, J., Homer, N., . . . Genome Project Data Processing, S. (2009). The Sequence Alignment/Map format and SAMtools. *Bioinformatics, 25*(16), 2078-2079. doi:10.1093/bioinformatics/btp352

Li, M., Tian, S., Jin, L., Zhou, G., Li, Y., Zhang, Y., . . . Li, R. (2013). Genomic analyses identify distinct patterns of selection in domesticated pigs and Tibetan wild boars. *Nat Genet, 45*(12), 1431-1438. doi:10.1038/ng.2811

Liedvogel, M., Maeda, K., Henbest, K., Schleicher, E., Simon, T., Timmel, C. R., . . . Mouritsen, H. (2007). Chemical Magnetoreception: Bird Cryptochrome 1a Is Excited by Blue Light and Forms Long-Lived Radical-Pairs. *PLoS One, 2*(10). doi:ARTN e1106

10.1371/journal.pone.0001106

McPherson, J. D., Marra, M., Hillier, L., Waterston, R. H., Chinwalla, A., Wallis, J., . . . Lehrach, H. (2001). A physical map of the human genome. *Nature, 409*(6822), 934-941. doi:10.1038/35057157

Palm, E. C., Newman, S. H., Prosser, D. J., Xiao, X., Ze, L., Batbayar, N., . . . Takekawa, J. Y. (2015). Mapping migratory flyways in Asia using dynamic Brownian bridge movement models. *Movement ecology, 3*(1), 3.

Pfeifer, B., Wittelsburger, U., Ramos-Onsins, S. E., & Lercher, M. J. (2014). PopGenome: an efficient Swiss army knife for population genomic analyses in R. *Mol Biol Evol, 31*(7), 1929-1936. doi:10.1093/molbev/msu136

Prado-Martinez, J., Sudmant, P. H., Kidd, J. M., Li, H., Kelley, J. L., Lorente-Galdos, B., . . . Marques-Bonet, T. (2013). Great ape genetic diversity and population history. *Nature, 499*(7459), 471-475. doi:10.1038/nature12228

Purcell, S., Neale, B., Todd-Brown, K., Thomas, L., Ferreira, M. A., Bender, D., . . . Sham, P. C. (2007). PLINK: a tool set for whole-genome association and population-based linkage analyses. *Am J Hum Genet, 81*(3), 559-575. doi:10.1086/519795

Ruan, J., Li, H., Chen, Z., Coghlan, A., Coin, L. J., Guo, Y., . . . Durbin, R. (2008). TreeFam: 2008 Update. *Nucleic Acids Res, 36*(Database issue), D735-740. doi:10.1093/nar/gkm1005

Schmitt, P., Gueguen, Y., Desmarais, E., Bachere, E., & de Lorgeril, J. (2010). Molecular diversity of antimicrobial effectors in the oyster Crassostrea gigas. *BMC Evol Biol, 10*, 23. doi:10.1186/1471-2148-10-23

Tamura, K., Peterson, D., Peterson, N., Stecher, G., Nei, M., & Kumar, S. (2011). MEGA5: molecular evolutionary genetics analysis using maximum likelihood, evolutionary distance, and maximum parsimony methods. *Mol Biol Evol, 28*(10), 2731-2739. doi:10.1093/molbev/msr121

Trapnell, C., Pachter, L., & Salzberg, S. L. (2009). TopHat: discovering splice junctions with RNA-Seq. *Bioinformatics, 25*(9), 1105-1111. doi:10.1093/bioinformatics/btp120

Wang, G. D., Zhai, W., Yang, H. C., Fan, R. X., Cao, X., Zhong, L., . . . Zhang, Y. P. (2013). The genomics of selection in dogs and the parallel evolution between dogs and humans. *Nat Commun, 4*, 1860. doi:10.1038/ncomms2814

Wu, F., Kaczynski, T. J., Sethuramanujam, S., Li, R., Jain, V., Slaughter, M., & Mu, X. (2015). Two transcription factors, Pou4f2 and Isl1, are sufficient to specify the retinal ganglion cell fate. *Proc Natl Acad Sci U S A, 112*(13), E1559-1568. doi:10.1073/pnas.1421535112

Wu, L. Q., & Dickman, J. D. (2012). Neural correlates of a magnetic sense. *Science, 336*(6084), 1054-1057. doi:10.1126/science.1216567

Xu, Z., & Wang, H. (2007). LTR_FINDER: an efficient tool for the prediction of full-length LTR retrotransposons. *Nucleic Acids Res, 35*(Web Server issue), W265-268. doi:10.1093/nar/gkm286

Yang, J., Lee, S. H., Goddard, M. E., & Visscher, P. M. (2011). GCTA: a tool for genome-wide complex trait analysis. *Am J Hum Genet, 88*(1), 76-82. doi:10.1016/j.ajhg.2010.11.011

Yang, Z. (1997). PAML: a program package for phylogenetic analysis by maximum likelihood. *Comput Appl Biosci, 13*(5), 555-556.

Yu, X. J., Zheng, H. K., Wang, J., Wang, W., & Su, B. (2006). Detecting lineage-specific adaptive evolution of brain-expressed genes in human using rhesus macaque as outgroup. *Genomics, 88*(6), 745-751. doi:10.1016/j.ygeno.2006.05.008

Zhao, S., Zheng, P., Dong, S., Zhan, X., Wu, Q., Guo, X., . . . Wei, F. (2013). Whole-genome sequencing of giant pandas provides insights into demographic history and local adaptation. *Nat Genet, 45*(1), 67-71. doi:10.1038/ng.2494
