## supplemental FigS24 for "Genomic analyses reveal the origin of domestic ducks and identify different genetic underpinnings of wild ducks"

Scaffolds from KB742382.1 to KB742485.1

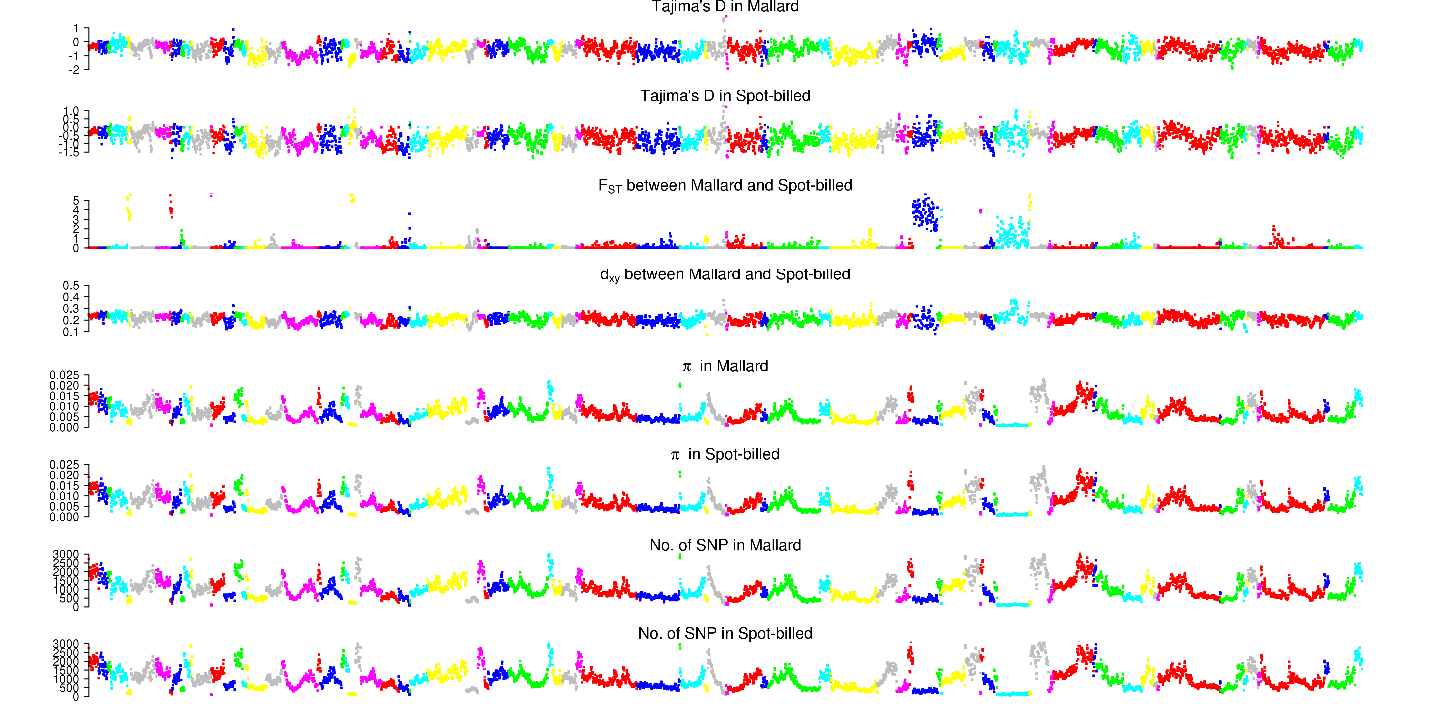

Scaffolds from KB742486.1 to KB742605.1

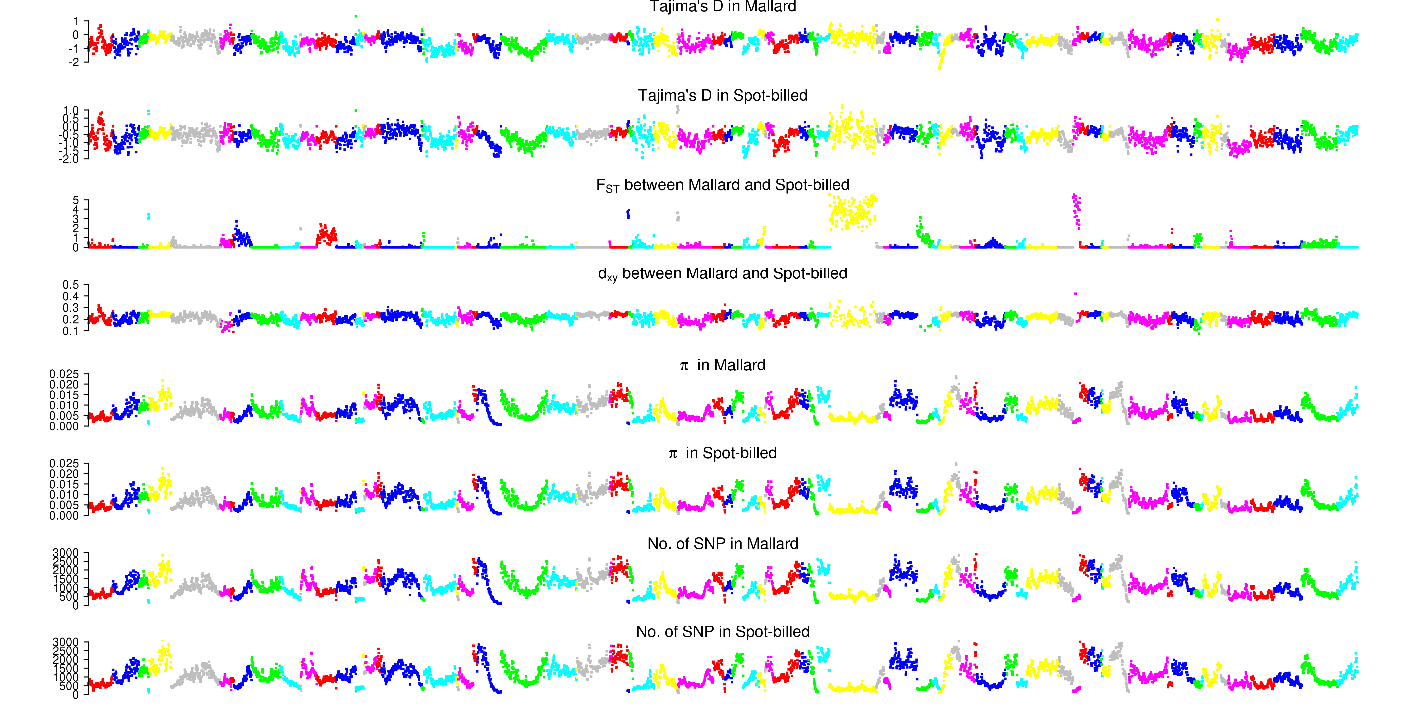

Scaffolds from KB742608.1 to KB742730.1

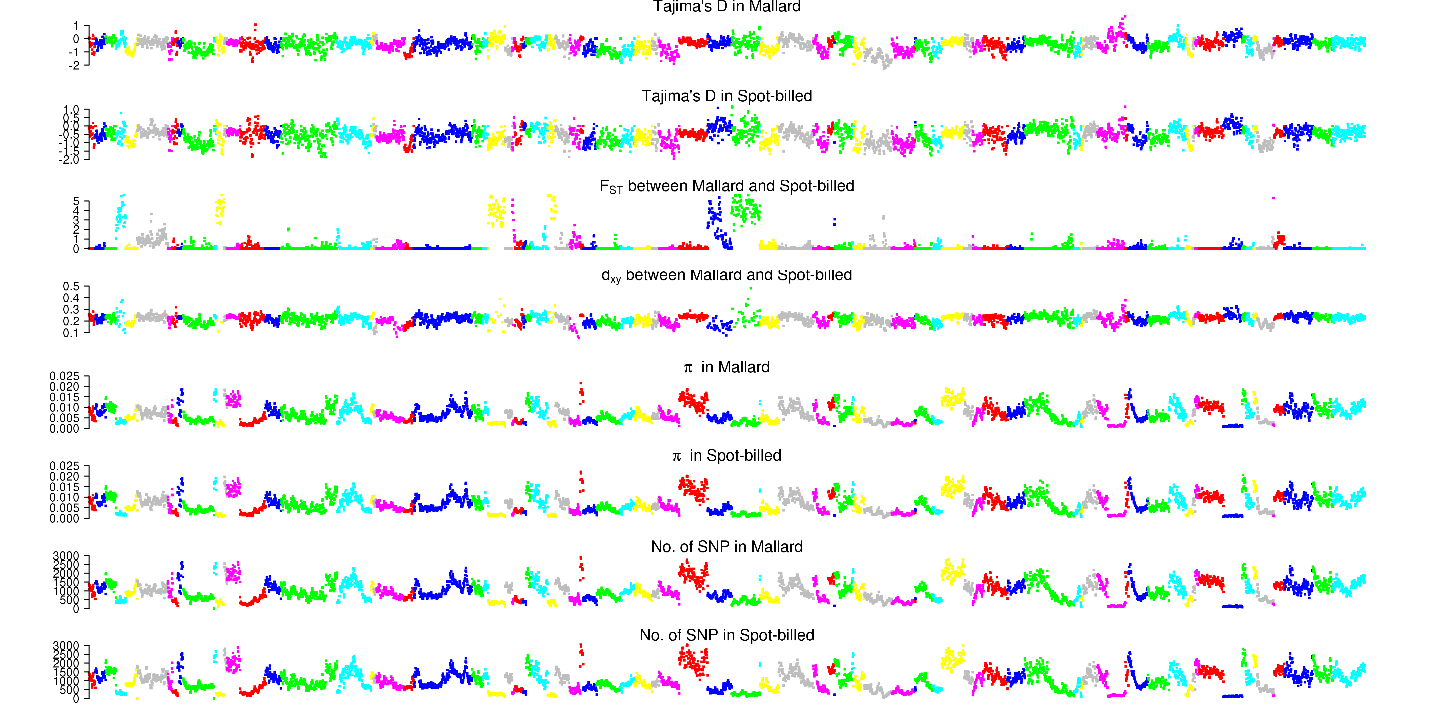

Scaffolds from KB742733.1 to KB742856.1

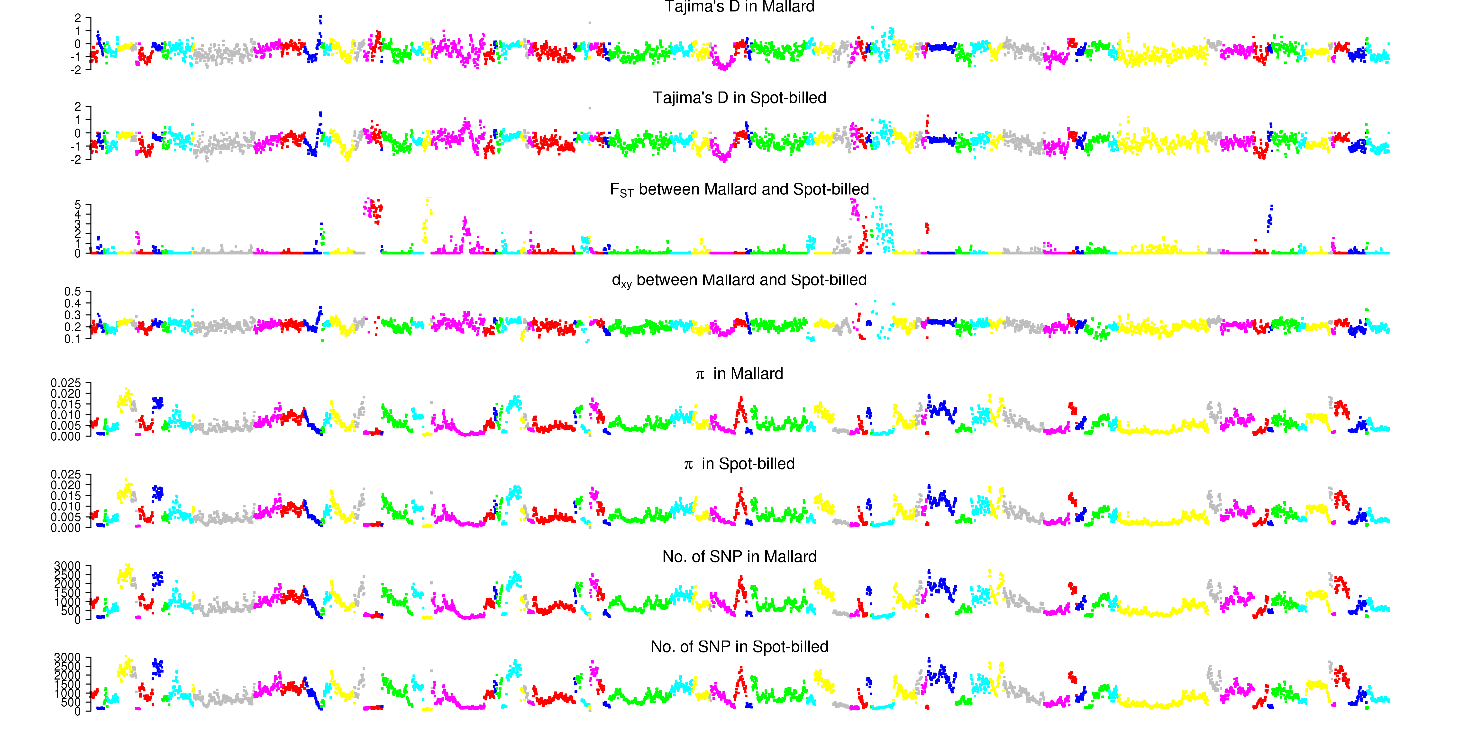

Scaffolds from KB742860.1 to KB742974.1

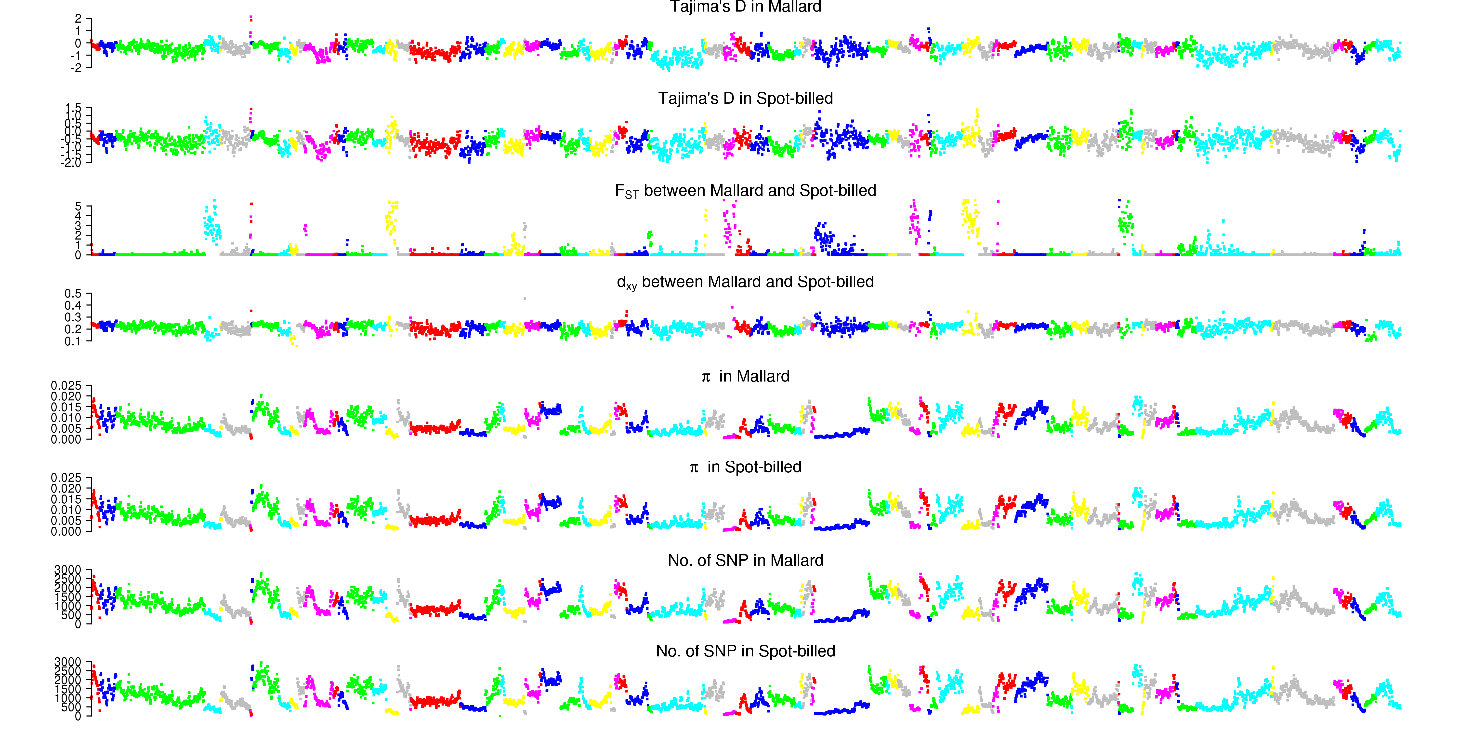

Scaffolds from KB742976.1 to KB743103.1

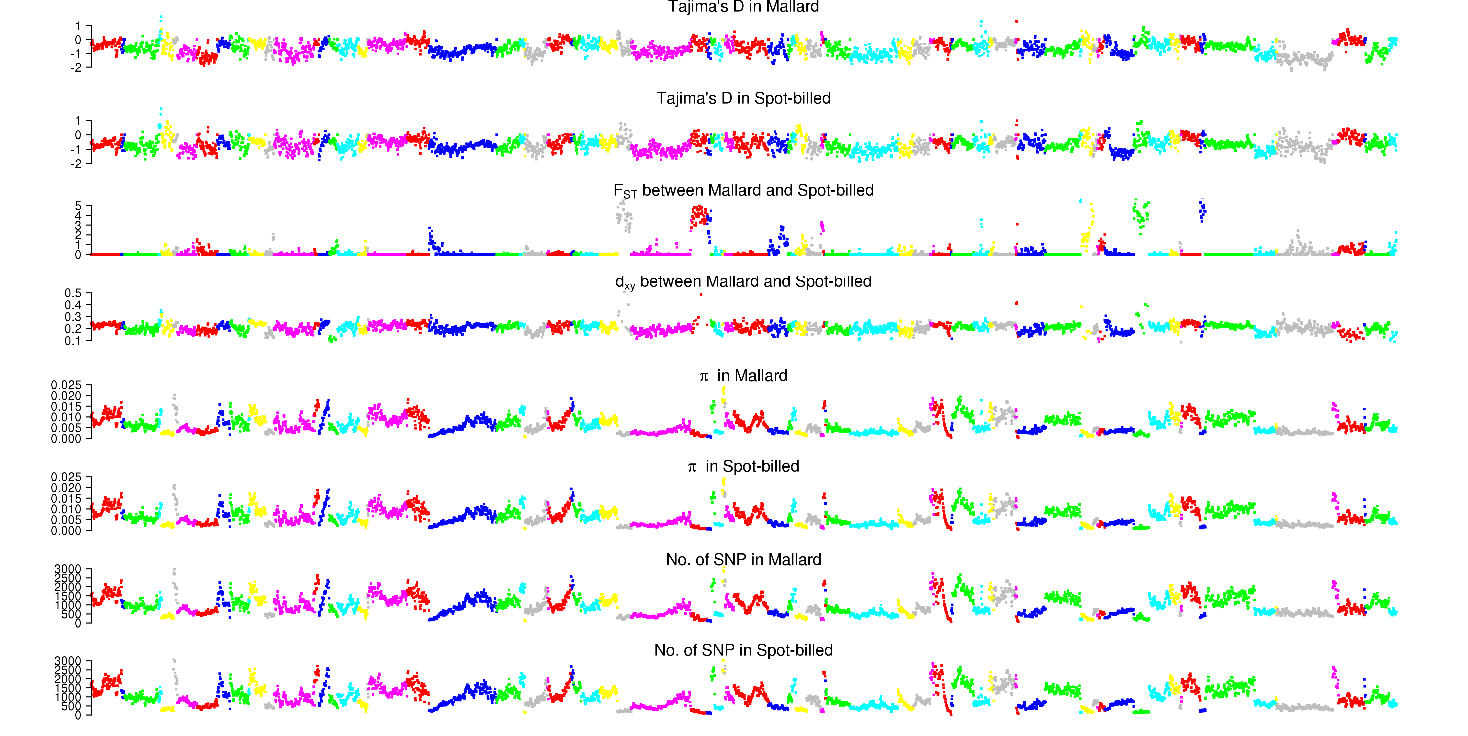

Scaffolds from KB743106.1 to KB743245.1

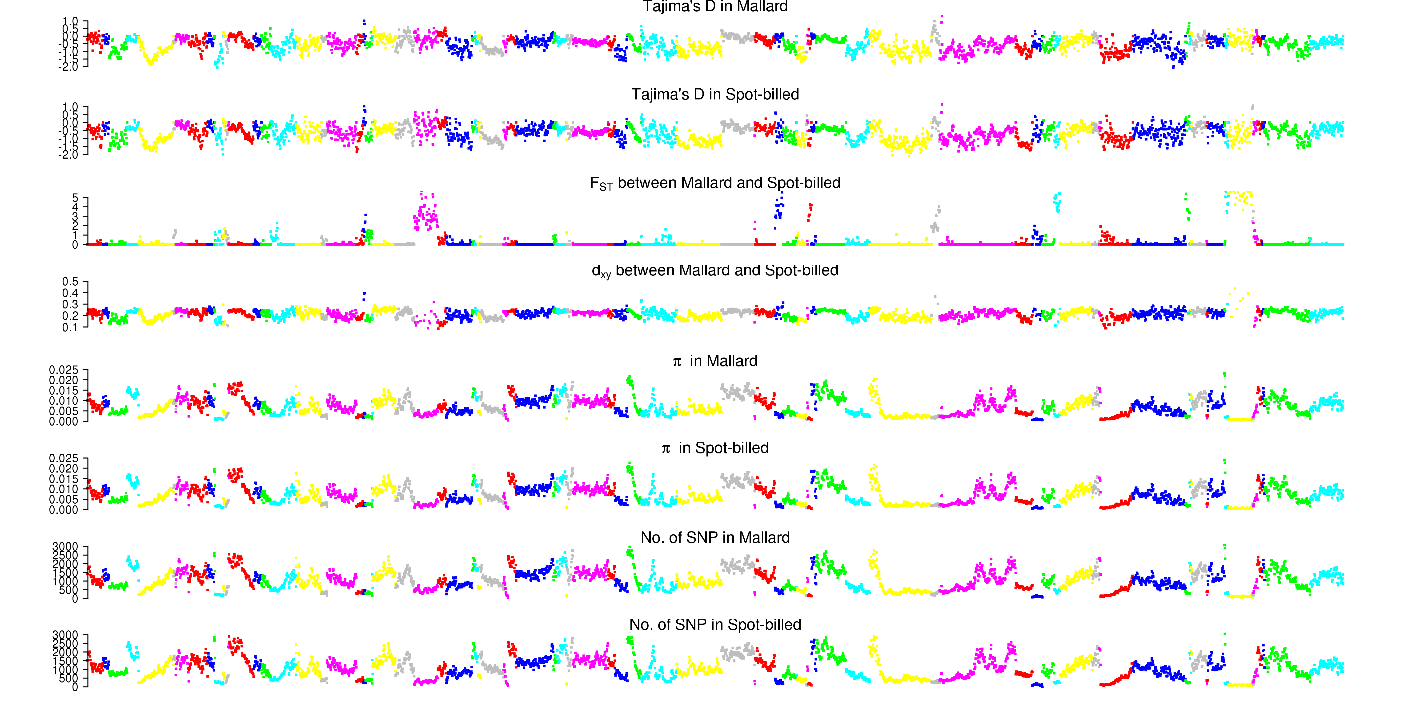

Scaffolds from KB743248.1 to KB743387.1

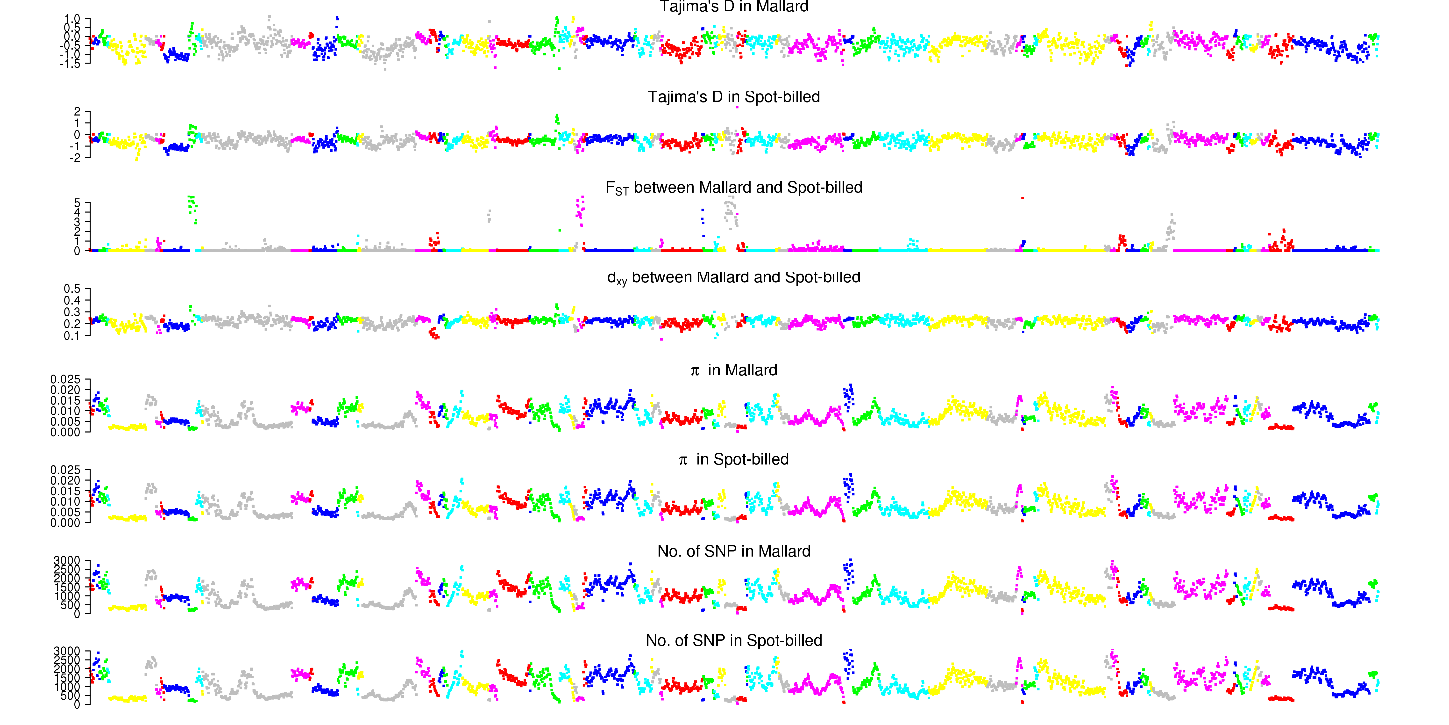

Scaffolds from KB743391.1 to KB743525.1

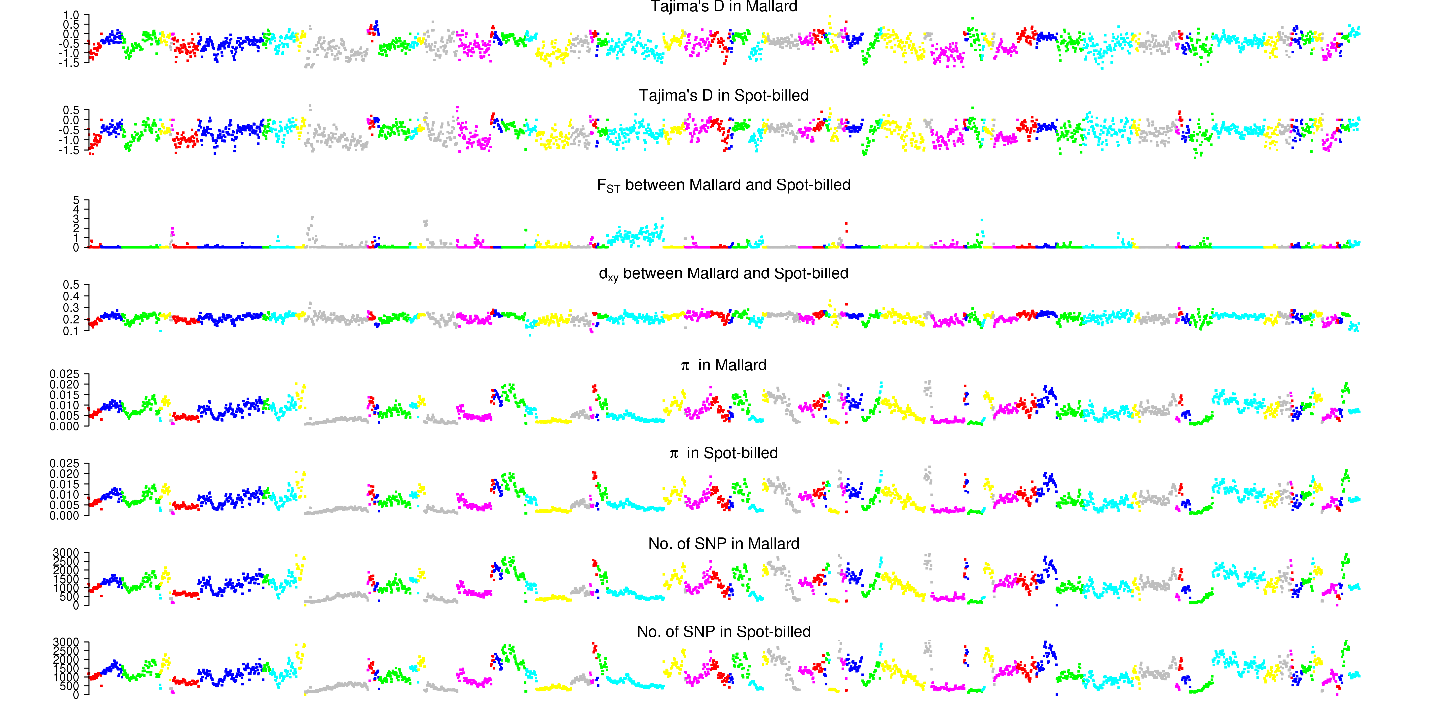

Scaffolds from KB743527.1 to KB743672.1

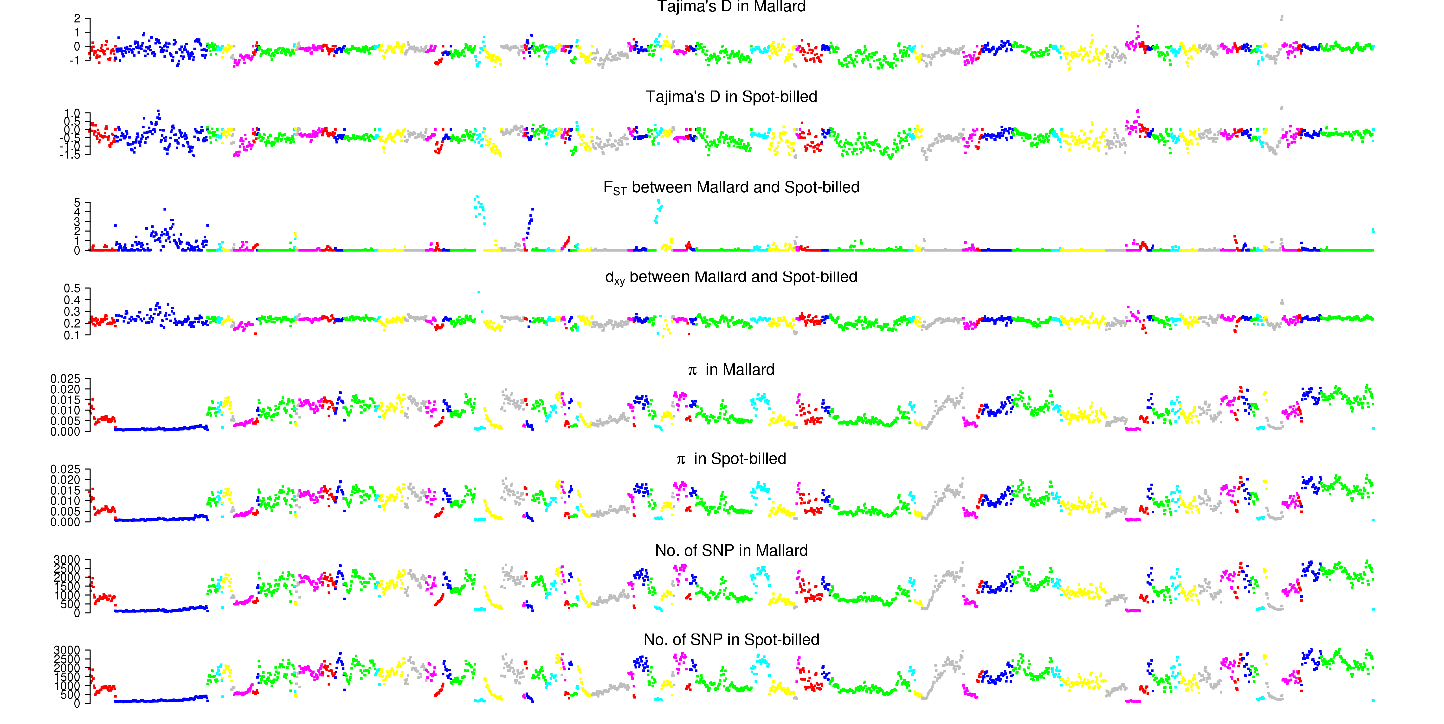

Scaffolds from KB743674.1 to KB743833.1

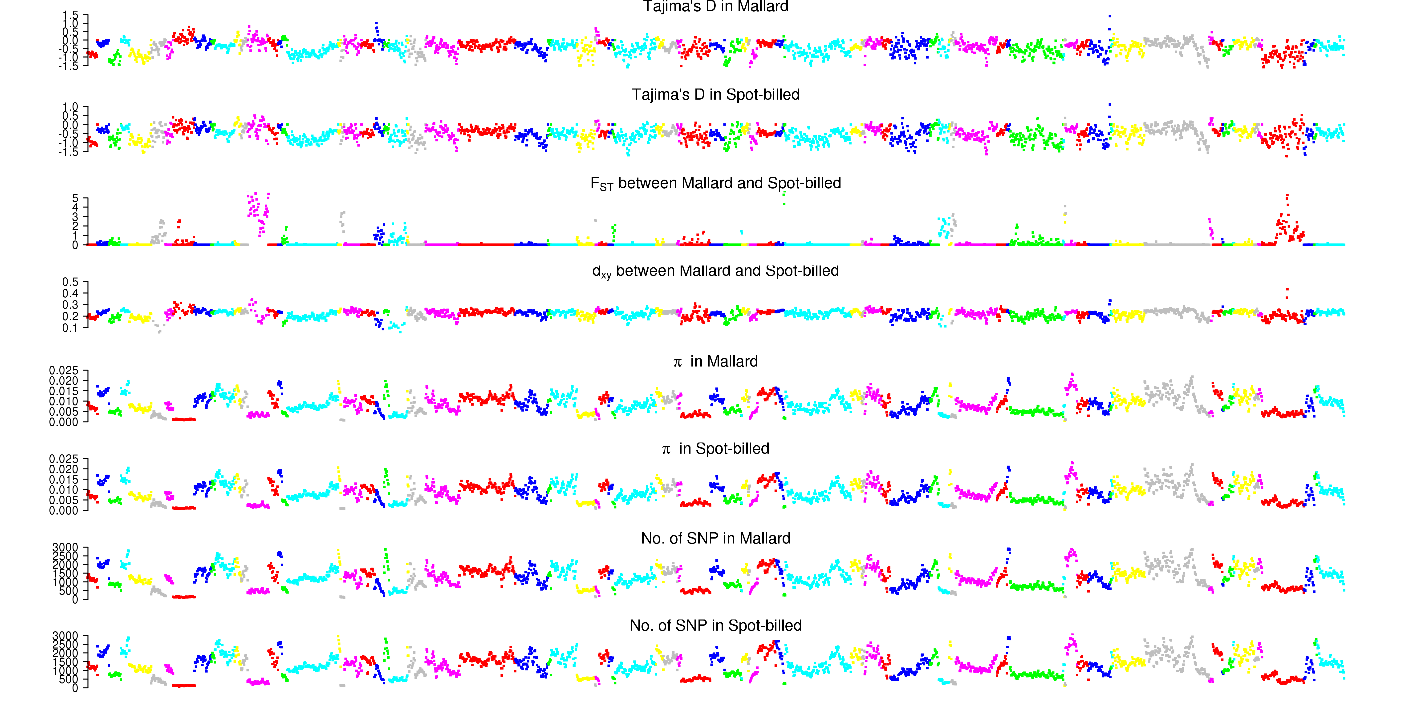

Scaffolds from KB743839.1 to KB744029.1

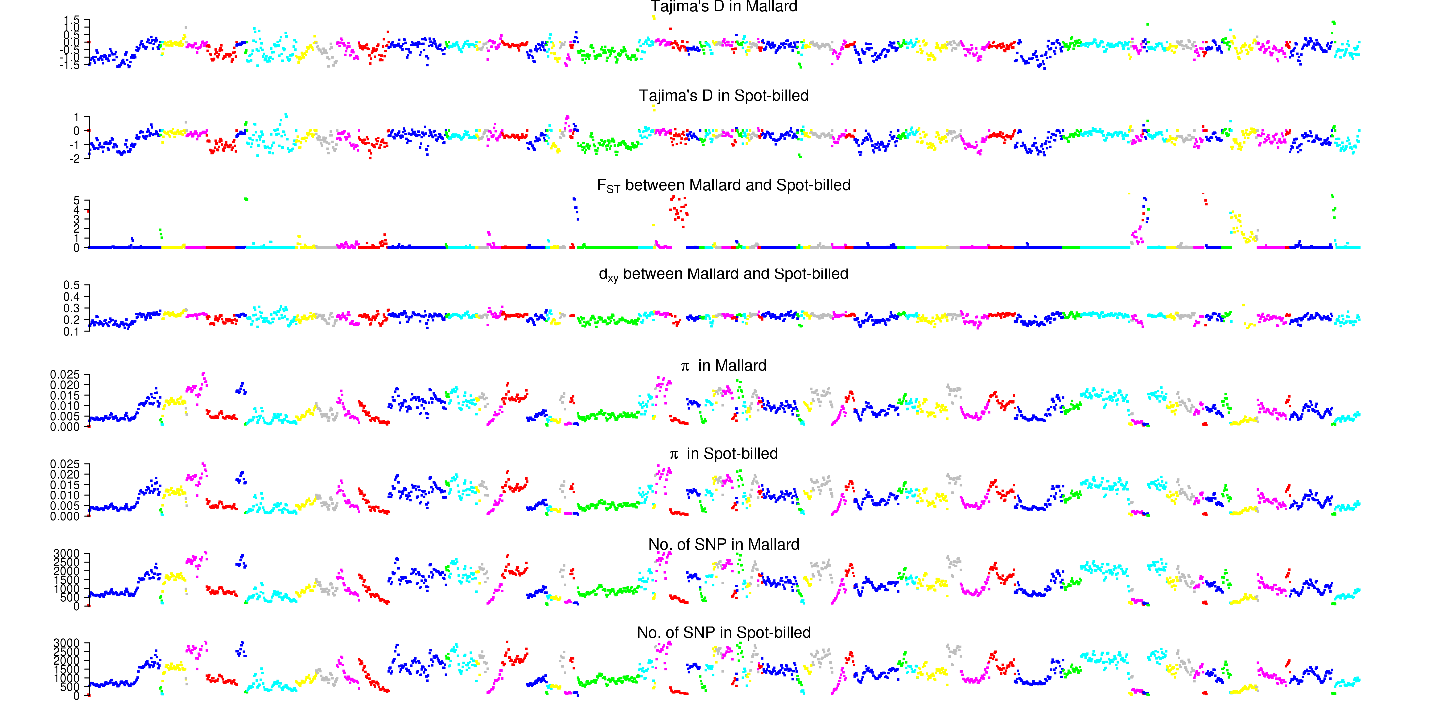

Scaffolds from KB744032.1 to KB744232.1

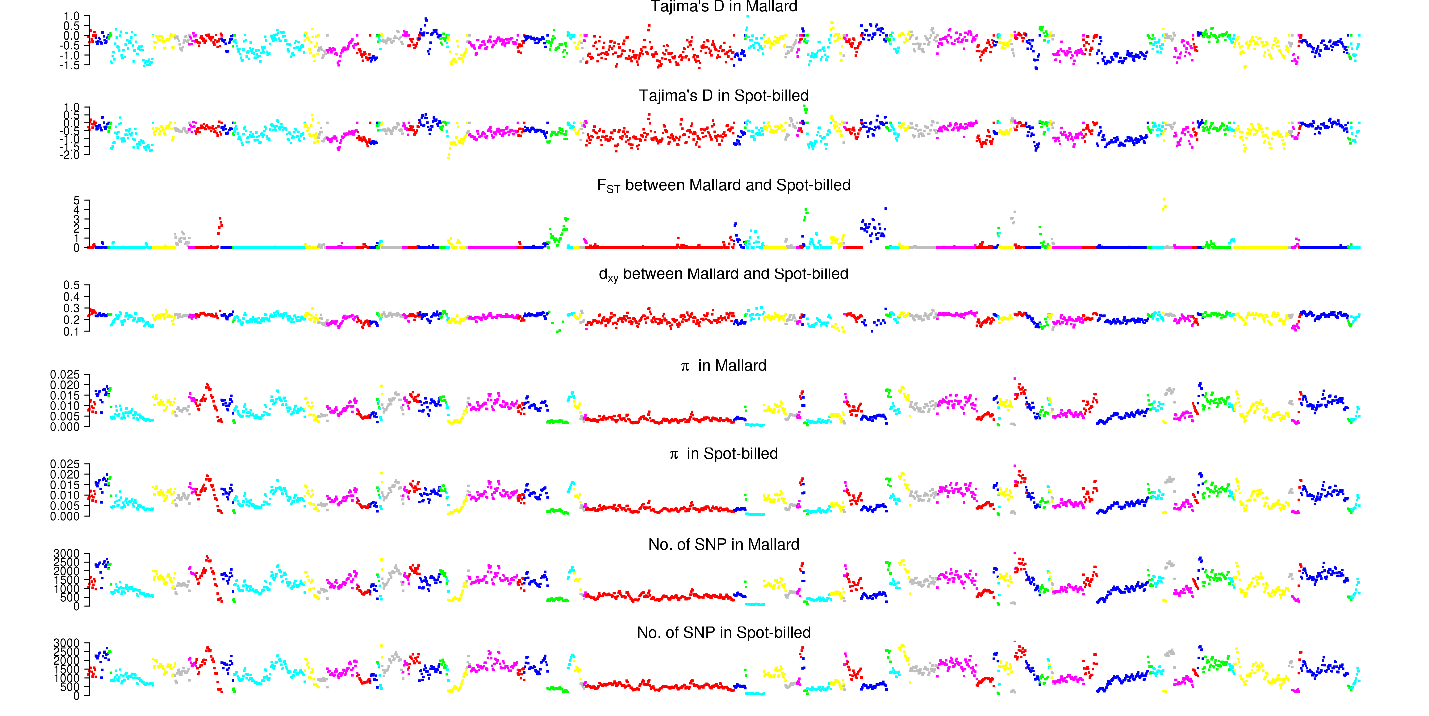

Scaffolds from KB744235.1 to KB744466.1

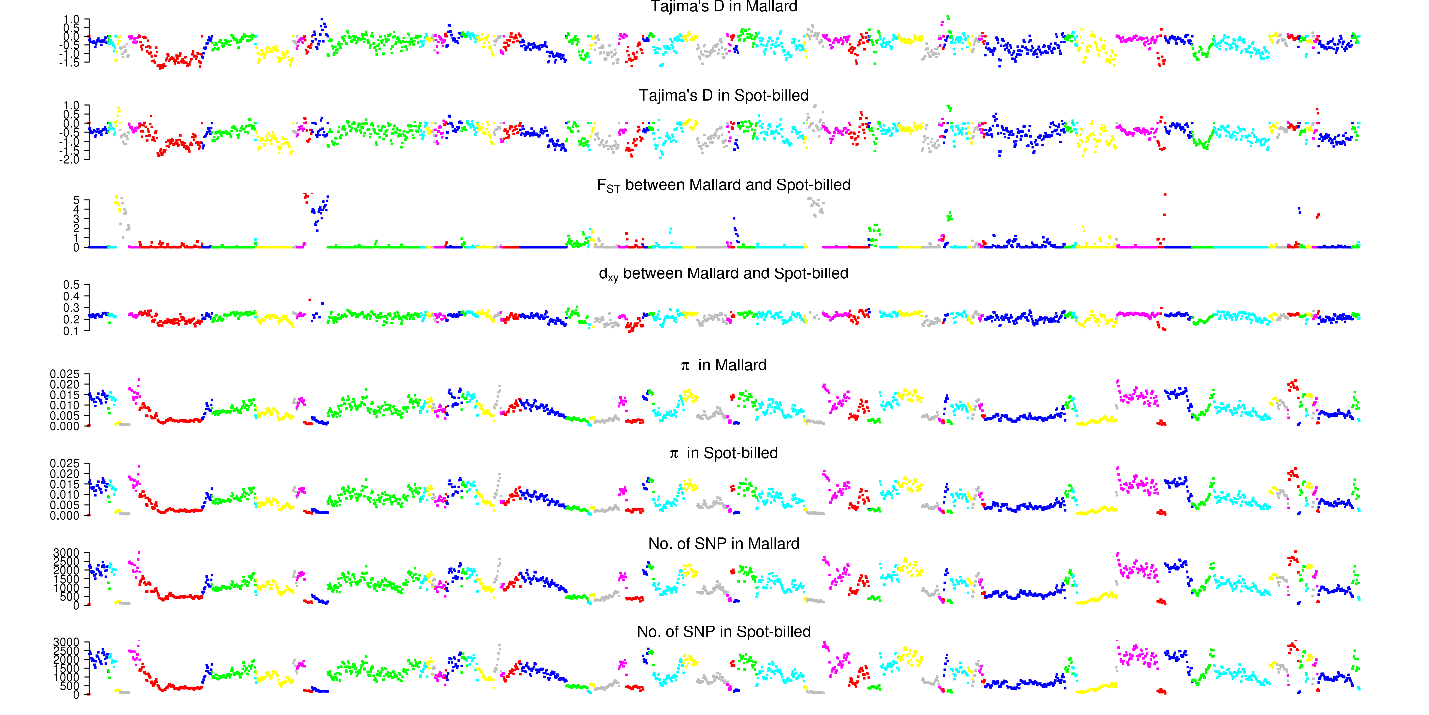

Scaffolds from KB744475.1 to KB744721.1

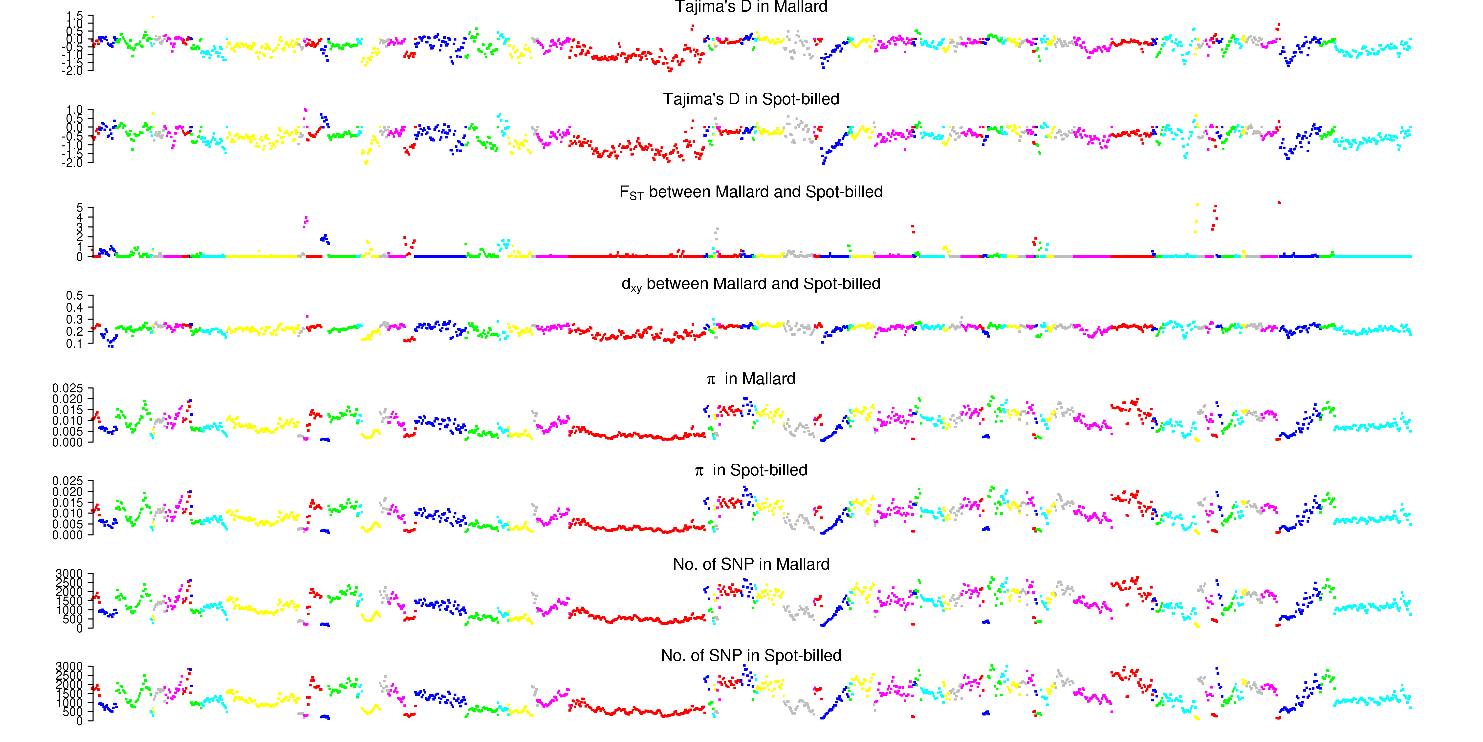

Scaffolds from KB744725.1 to KB745078.1
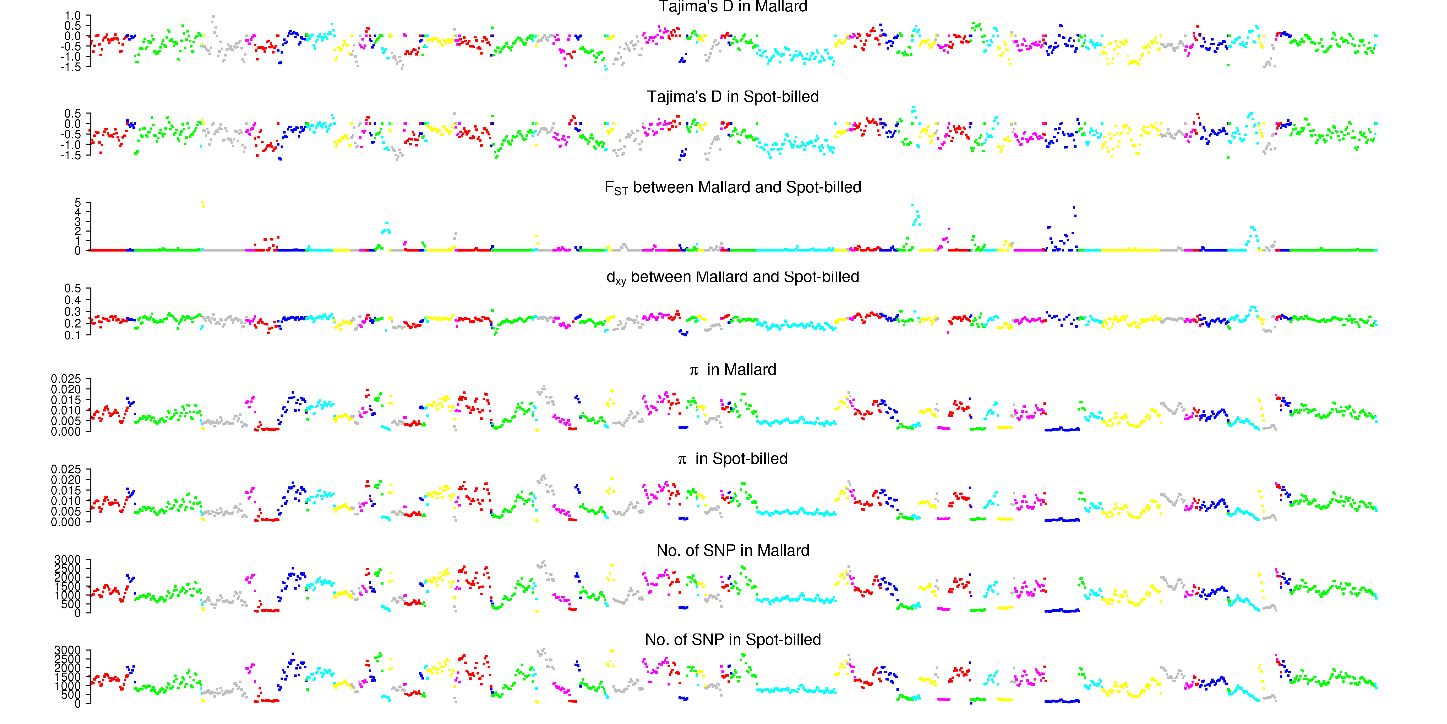

Scaffolds from KB745085.1 to KB745764.1

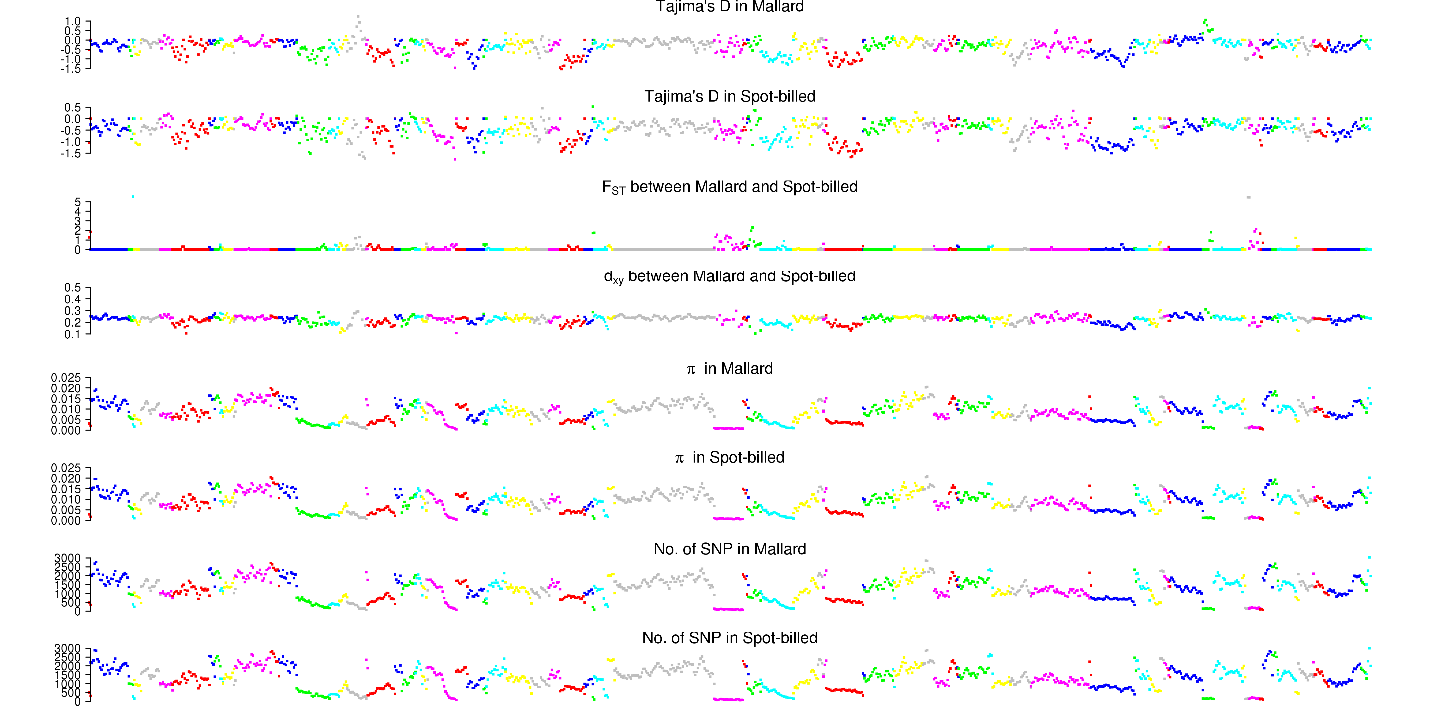

Scaffolds from KB745806.1 to KB747284.1
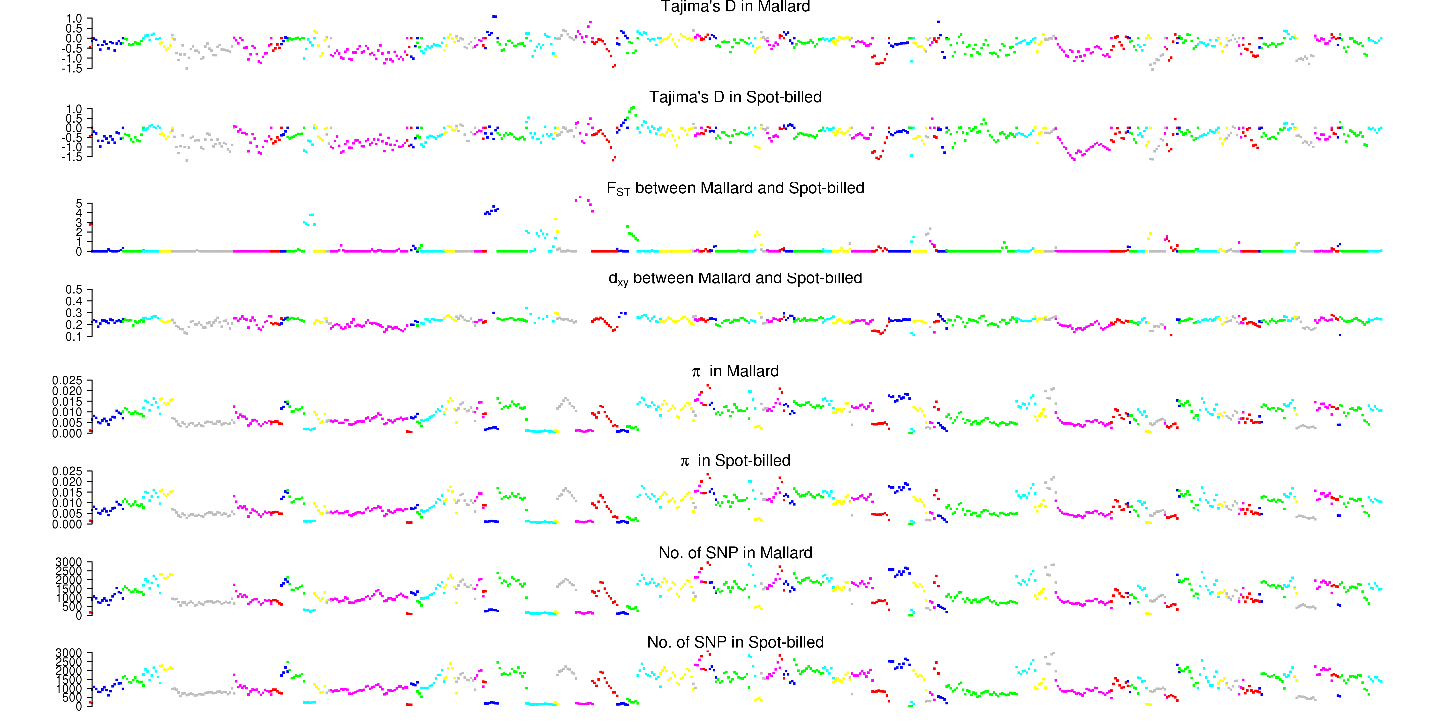

### Fig. S23. Distribution of Tajima’D, positive end of z*F_ST_*, dxy, number of SNPs per window, nucleotide diversity (π) and zHp along scaffolds in ducks.

These numbers were counted with 40 kb window by step size of 20 kb.
